## Additional file 1 for "Methods detecting rhythmic gene expression are biologically relevant only for strong signal"

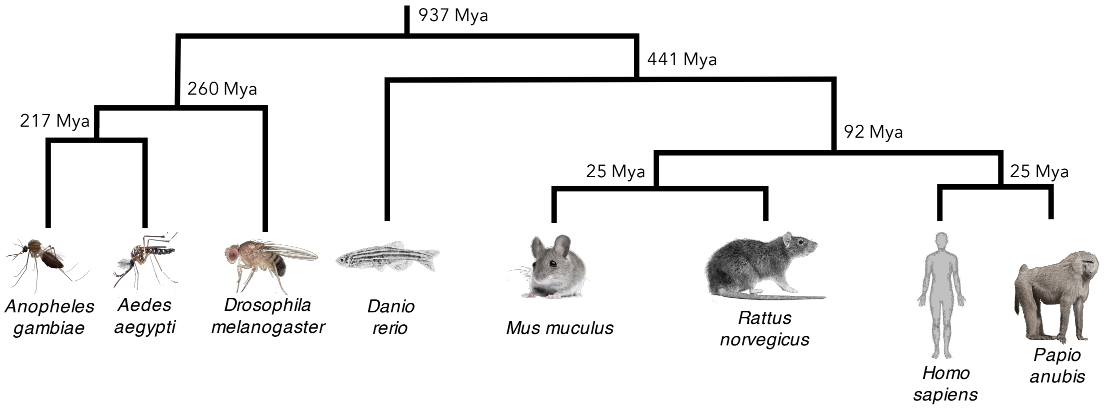


**Fig S1.** Phylogenetic tree of species studied.

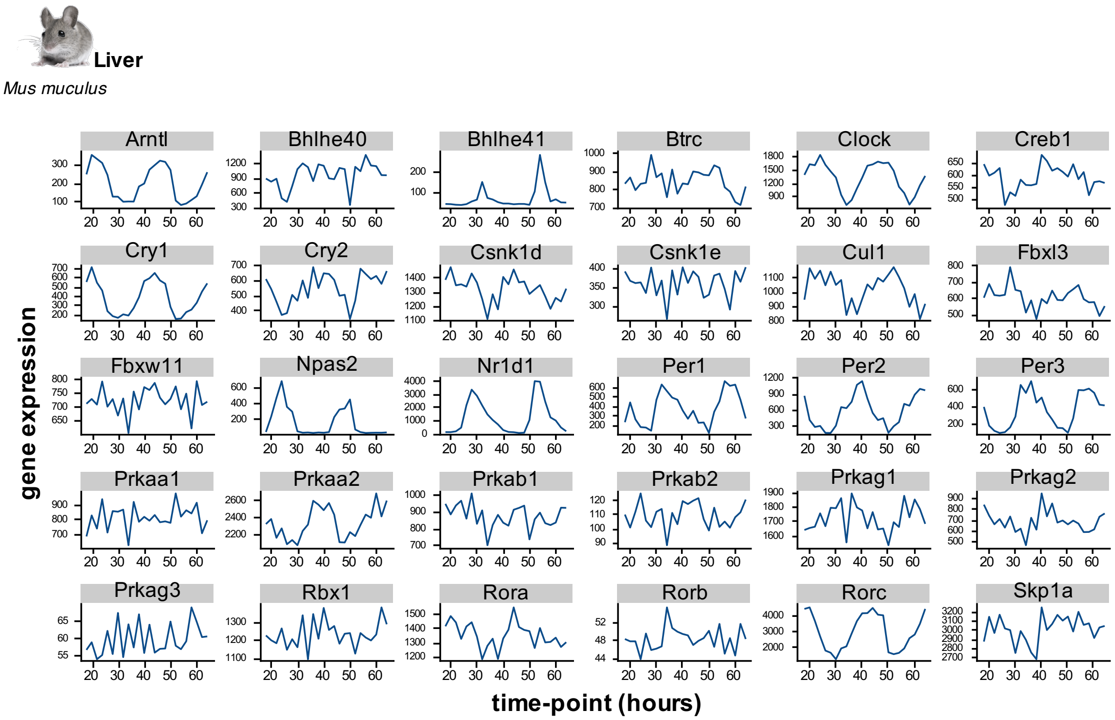


**Fig S2.** Expression over time of some known circadian genes in liver from mouse experiment data [2] (microarray).

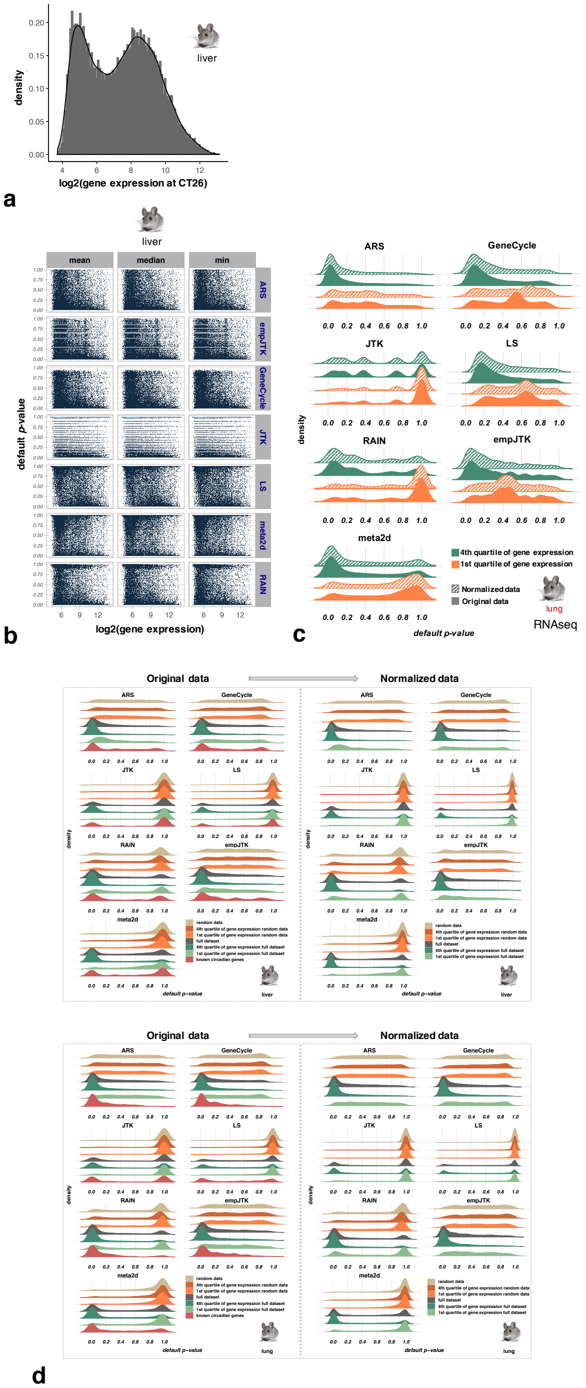


**Fig S3.** **a)** Typical bimodal density distribution of gene expression at time-point CT26 from mouse liver data (microarray).
**b)** Gene expression per time-point, calculated with the median, the mean, or the minimum of time-points, as a function of default $p$-values obtained for the seven methods applied to mouse liver data.
**c)** Methods applied to original vs normalized gene expression values of mouse lung data (RNAseq) produce the same distributions of $p$-values within highly expressed genes, or within lowly expressed genes.
**d)** Density distribution of default $p$-values obtained for the seven methods applied to original vs normalized mouse liver and lung data, sub-categorized in seven sub-groups (See the main text).

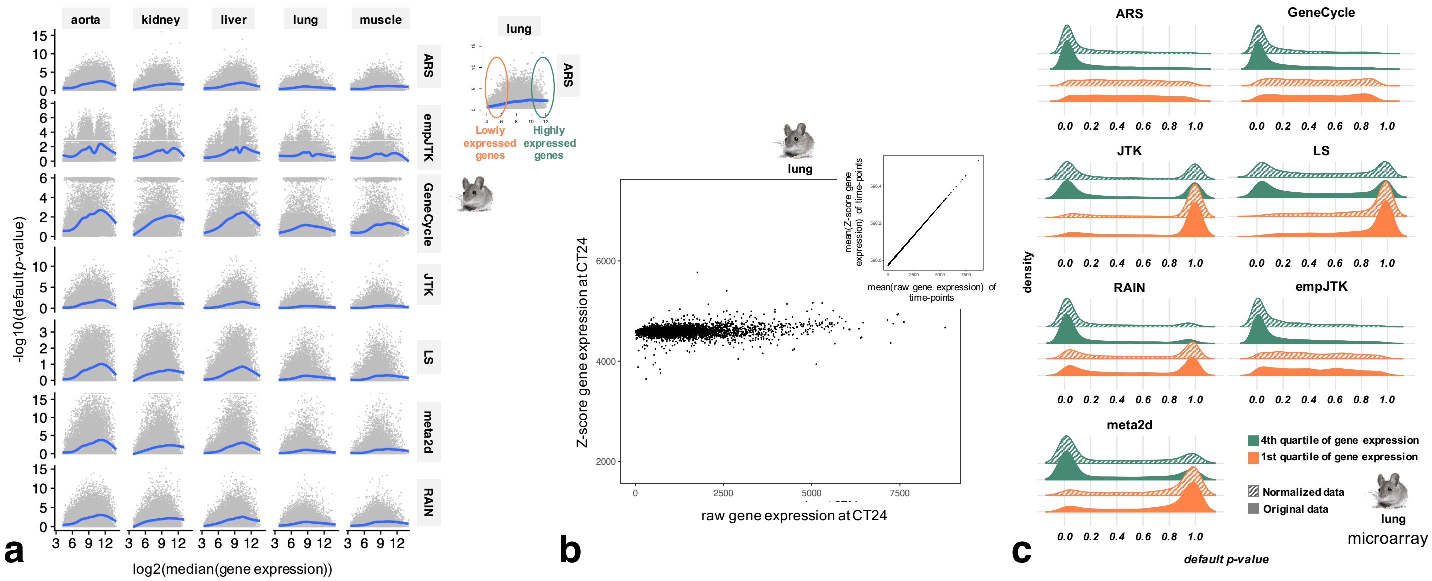


**Fig S4.** Detected rhythmic genes are enriched in highly expressed genes.

**a)** Scatter plot showing the relation between the expression level of genes (median of time-points) and their default $p$-values. The blue line is the smoothed conditional mean. Higher expression levels imply a higher power to detect rhythmic patterns.

**b)** Control of the normalization by Z-score of gene expression values at a given time-point (CT24).

**c)** Methods applied to original vs normalized gene expression values produce the same distributions of $p$-values within highly expressed genes, or within lowly expressed genes. Particularly, the normalization of gene expression values does not allow to recover rhythmicity within lowly expressed genes.

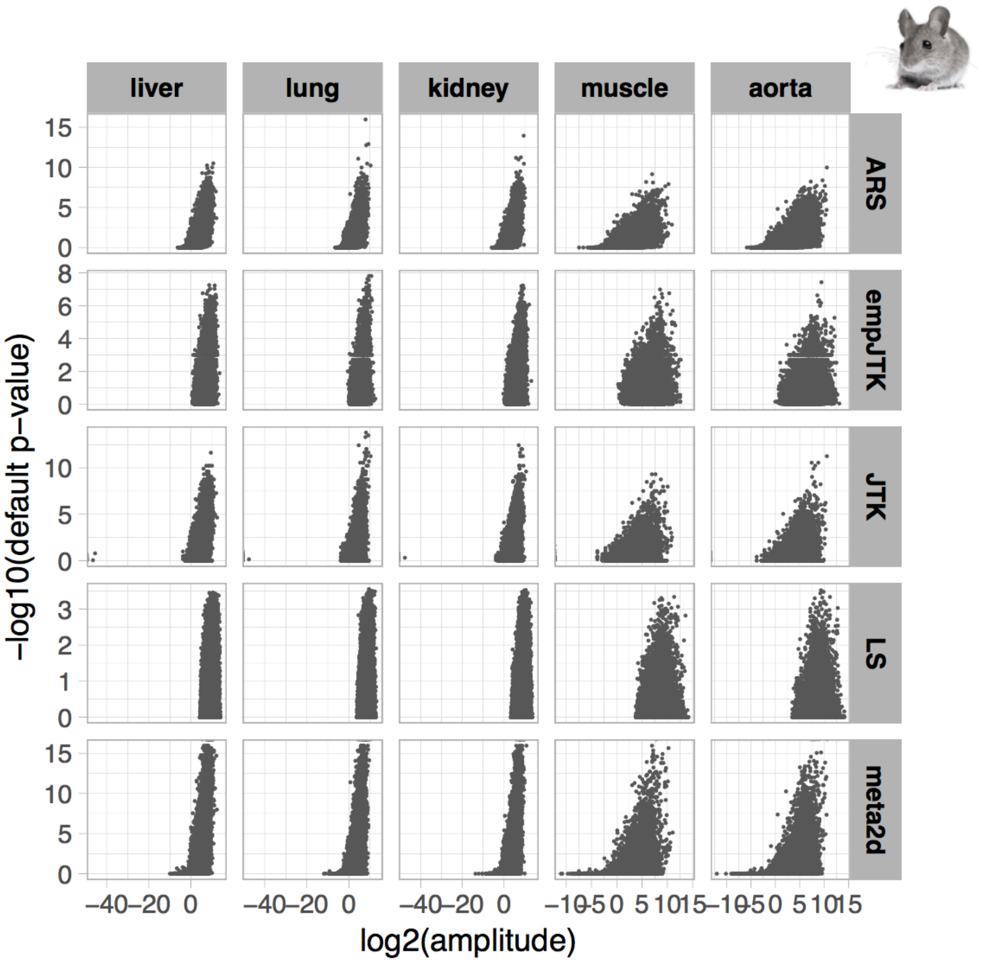


**Fig S5.** Scatter-plots showing the amplitude of gene expression as a function their default $p$-values obtained for the seven methods applied to five mouse tissues (microarray).

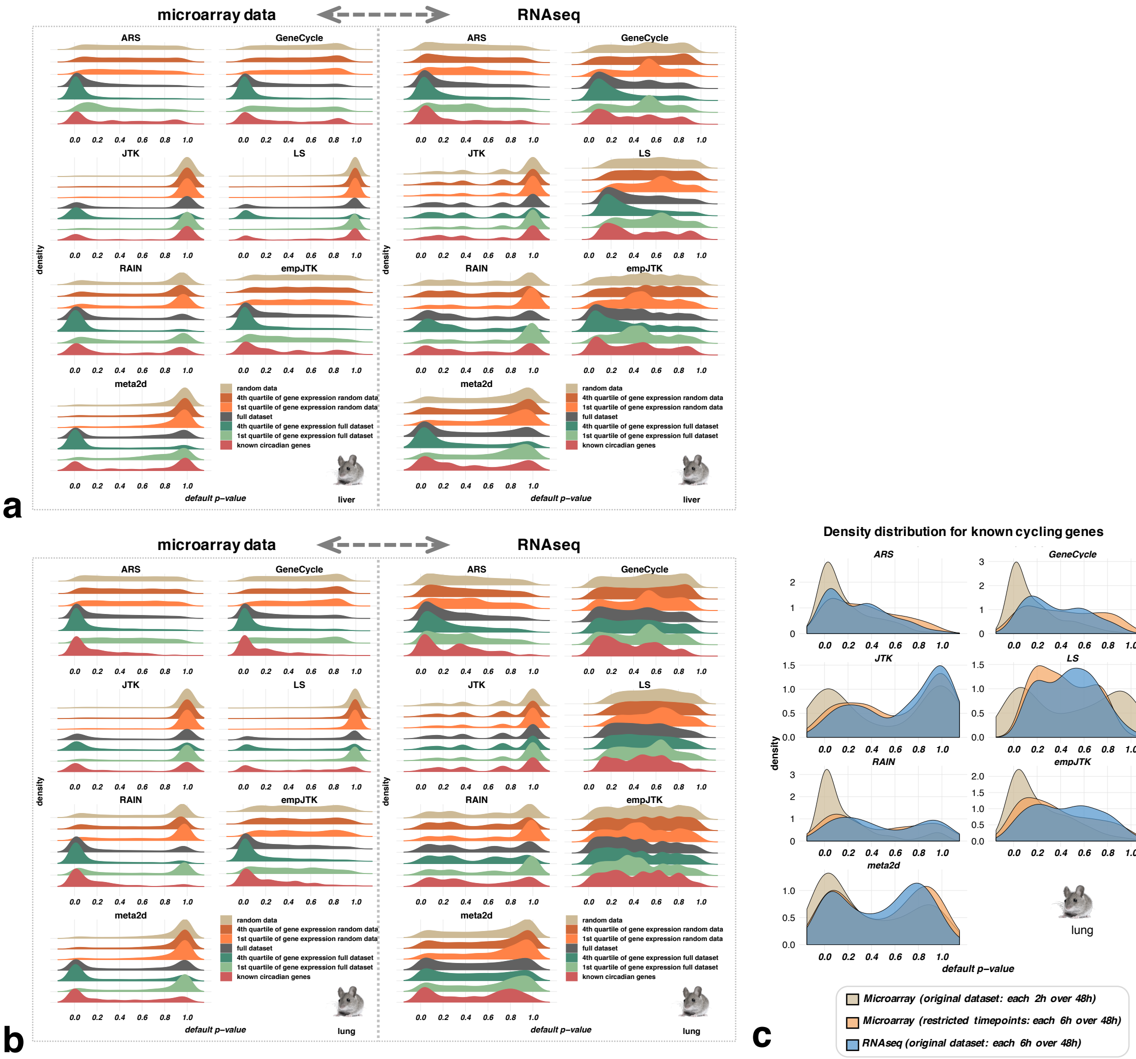


**Fig S6. a)b)** *p*-values distributions obtained from microarray vs RNAseq in mouse liver (**a**) and lung (**b**).
**c)** The restriction of microarray time-series to the same time-points as in the RNAseq series applied to known cycling genes produces similar *p*-value distributions to those obtained with RNAseq.

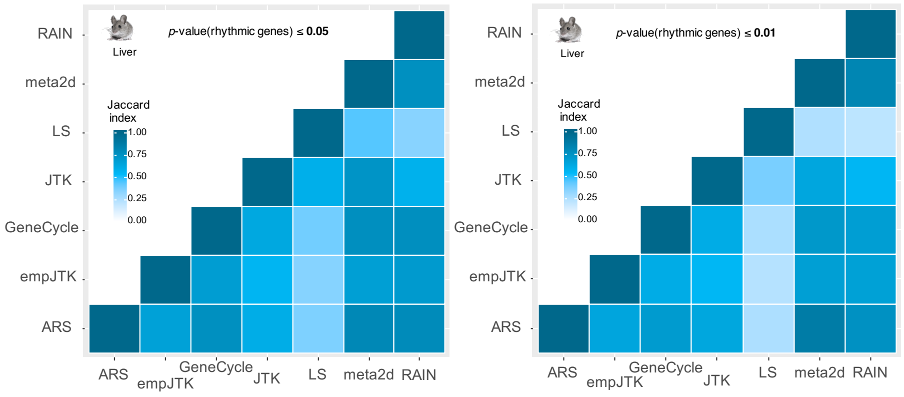


**Fig S7.** Heatmaps of Jaccard-indices comparing the similarity of genes called rhythmic (default $p$-value $\leq 0.05$ or 0.01) between the methods applied to mouse liver dataset (microarray).

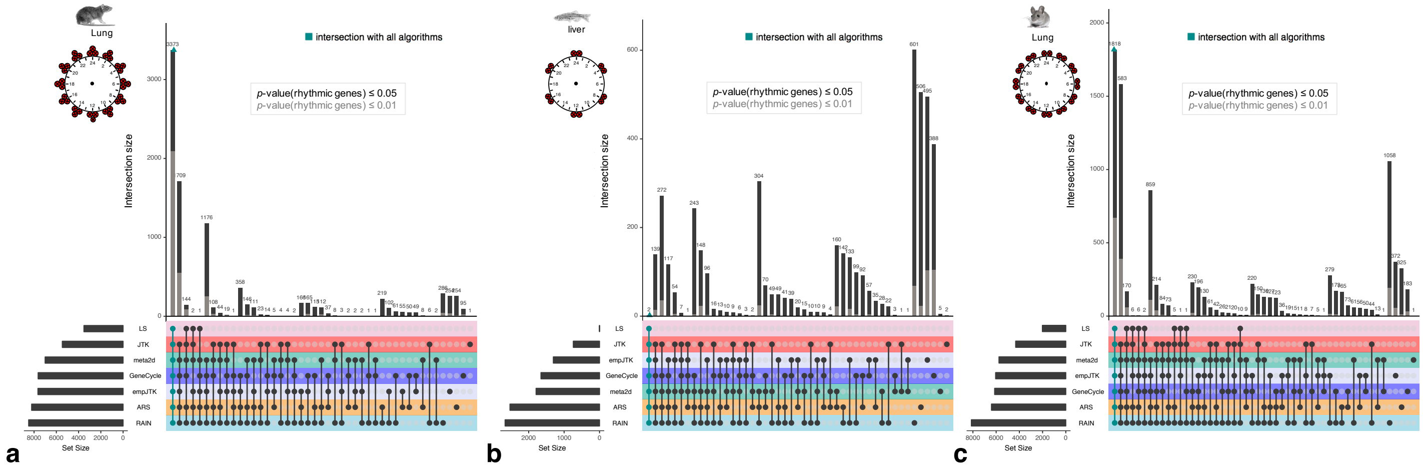


**Fig S8.** Upset diagram for mouse lung dataset (microarray) (**a**), zebrafish liver dataset (**b**), and mouse lung dataset (**c**) for the $p$-value thresholds of 0.05 (black) or 0.01 (grey) for calling genes rhythmic.

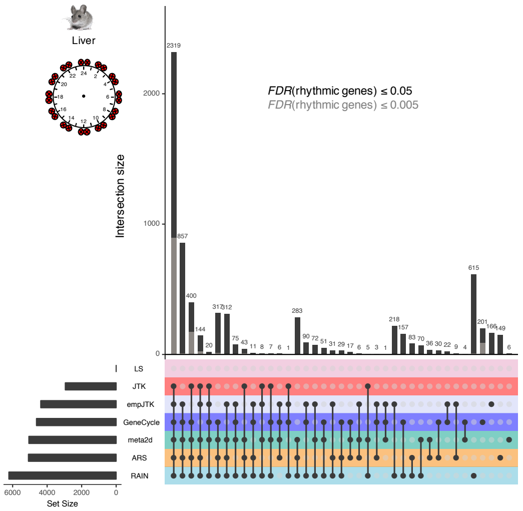


**Fig S9**. Using a very low false positive tolerance, all methods except LS overlap largely.
Upset diagram for mouse liver dataset (microarray) for the FDR thresholds of 0.05 (black) or 0.005 (grey) for calling genes rhythmic. FDR correspond to the false discovery rate adjustment of default $p$-values using `p.adjust` R function.

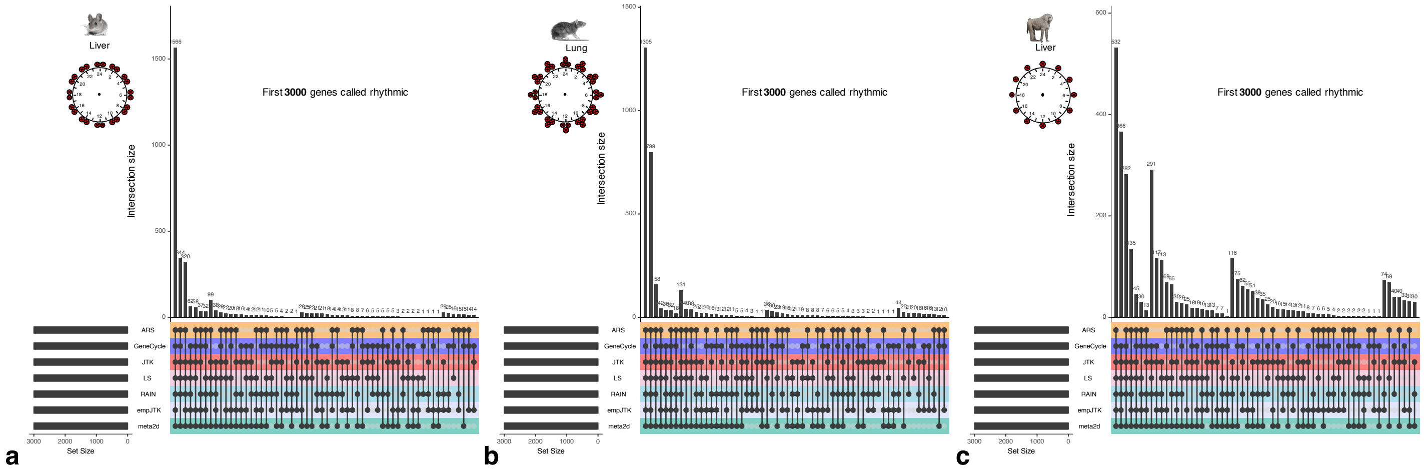


**Fig S10.** Upset diagram for mouse liver dataset (microarray) (**a**), rat lung dataset (**b**), and baboon liver dataset (**c**) for the first 3000 genes detected rhythmic for each method.

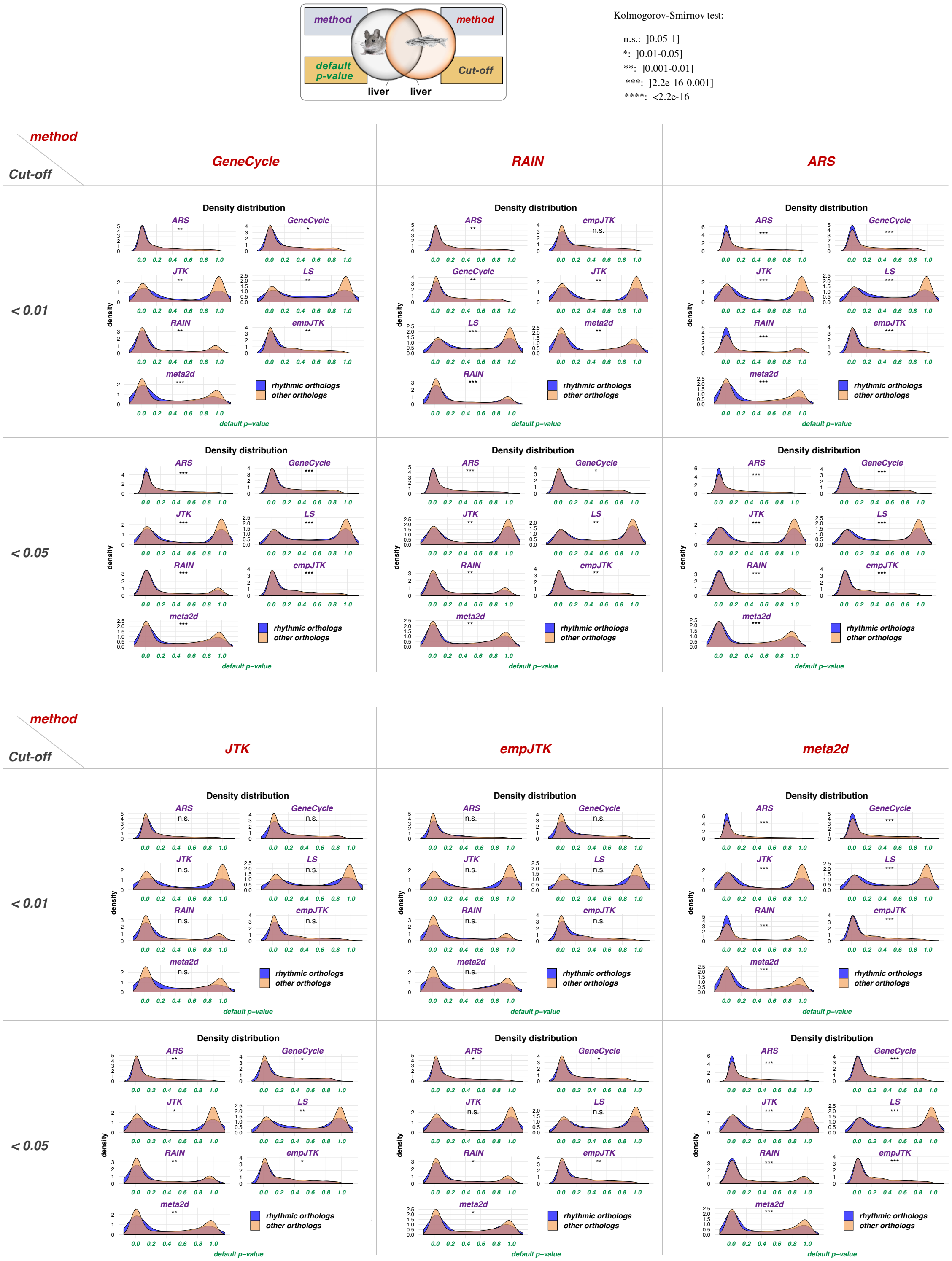


**Fig S11.** Default $p$-values density distribution of rhythmic orthologs vs non-rhythmic orthologs obtained for the seven methods applied to mouse liver data. Rhythmic orthologs are mouse-zebrafish 1:1 orthologs detected rhythmic in zebrafish liver each rhythm detection method (in red) and a $p$-value threshold of 0.01 or 0.05.

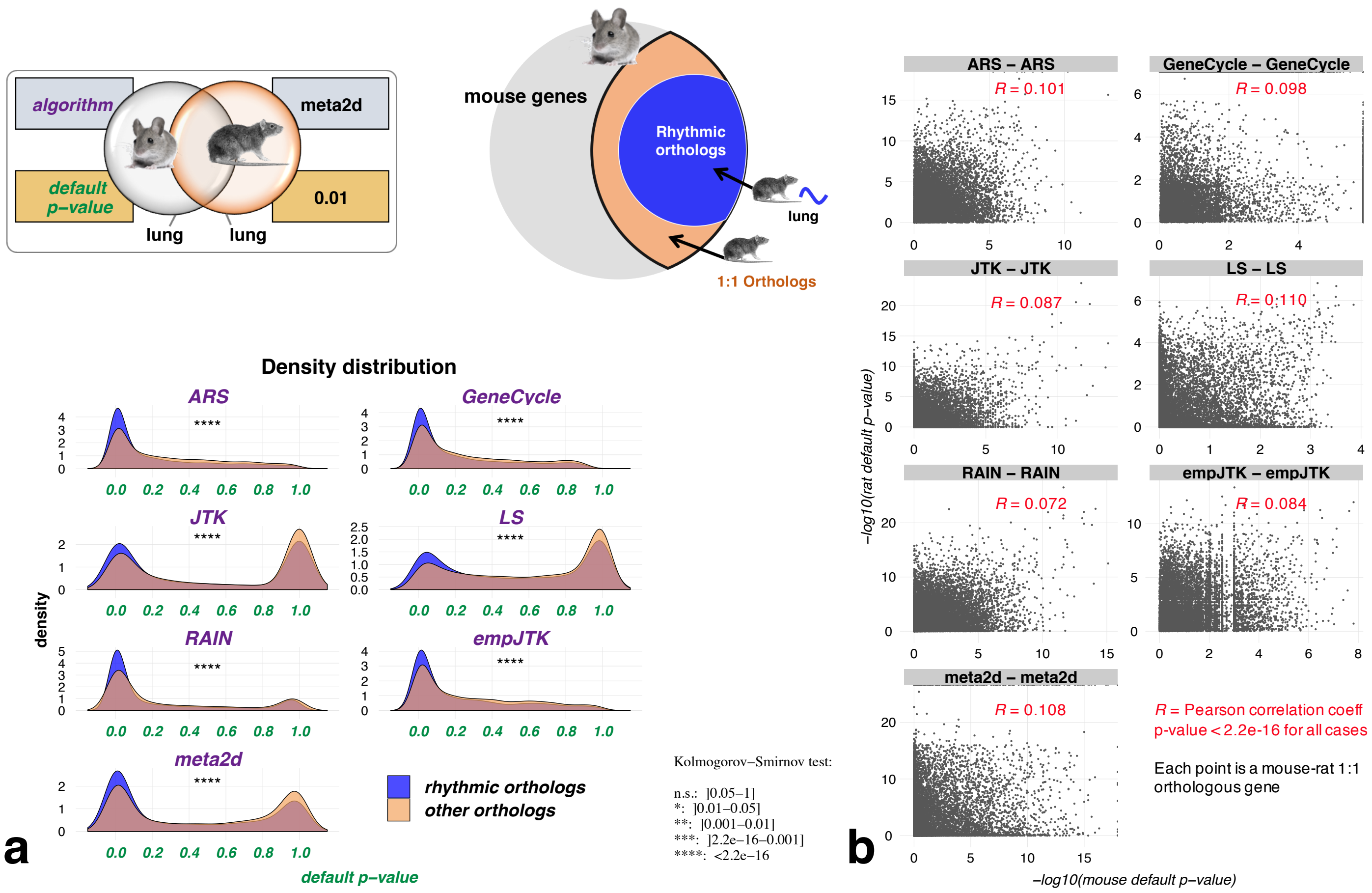


**Fig S12. a)** Distribution of default $p$-values of rhythmic and non-rhythmic mouse-rat orthologs obtained for the seven methods applied to mouse lung data. Rhythmic genes of rat lung are detected using the meta2d method and a $p$-value threshold of 0.01.
**b)** Scatter-plot comparing $p$-values for each mouse-rat one-to-one orthologous gene obtained for the seven methods (same method used for each comparison). $R$ is the Pearson correlation coefficient obtained with a significance $p$-value $< 2.2\text{e}{-16}$.

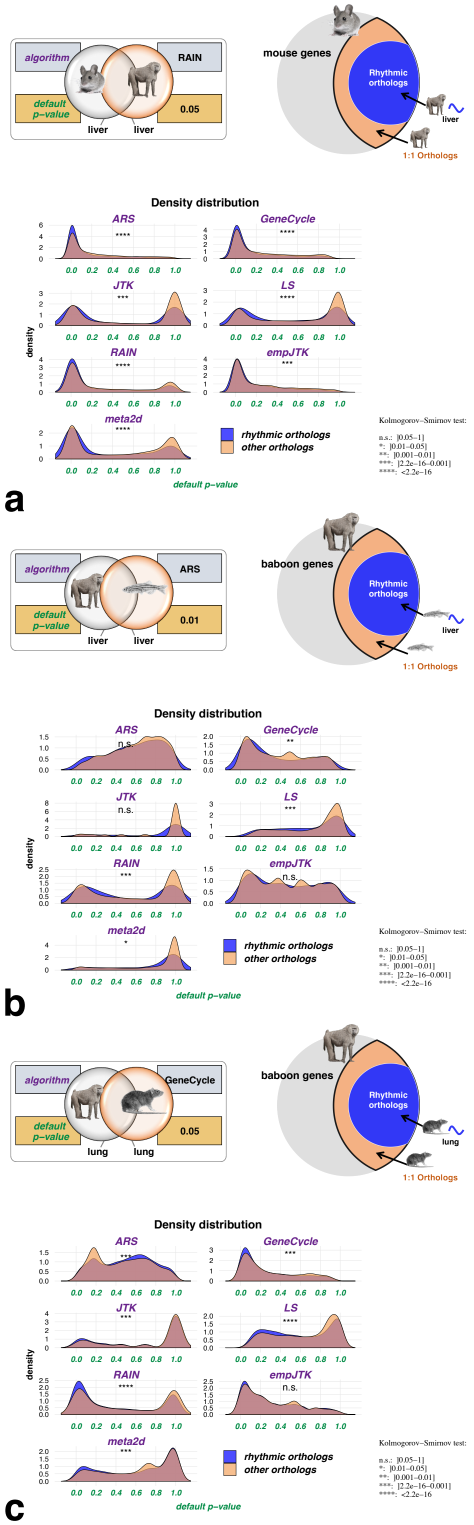


**Fig S13.** Distribution of default $p$-values of rhythmic and non-rhythmic mouse-baboon (**a**), baboon-zebrafish (**b**), and baboon-rat (**c**) orthologs obtained for the seven methods applied to the homologous tissue of species_1.

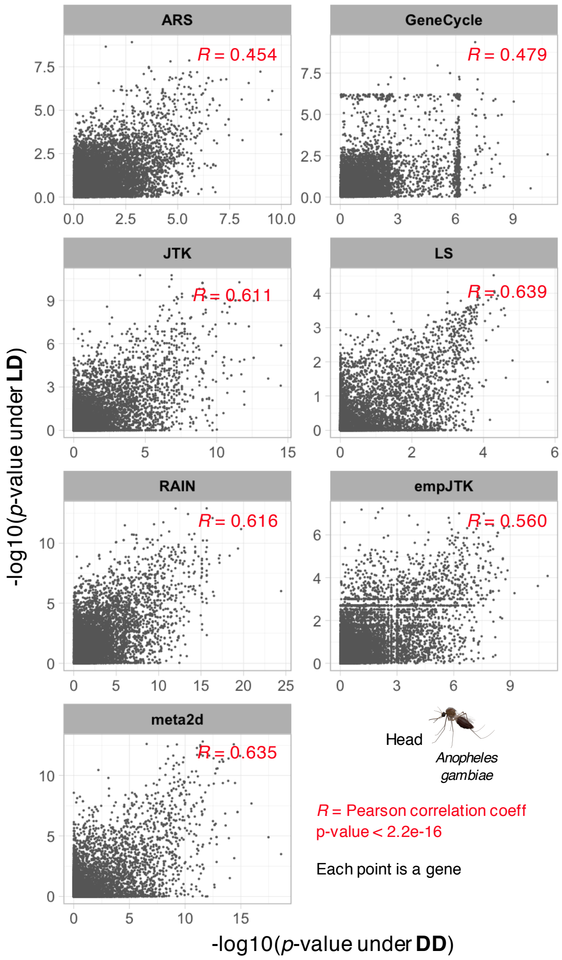


**Fig S14.** Scatter-plot comparing $p$-values obtained for the seven methods applied to the head dataset of *Anopheles gambiae* under light-dark (LD) or dark-dark (DD) conditions [1]. $R$ is the Pearson correlation coefficient obtained with a significance $p$-value $< 2.2\text{e}{-16}$.

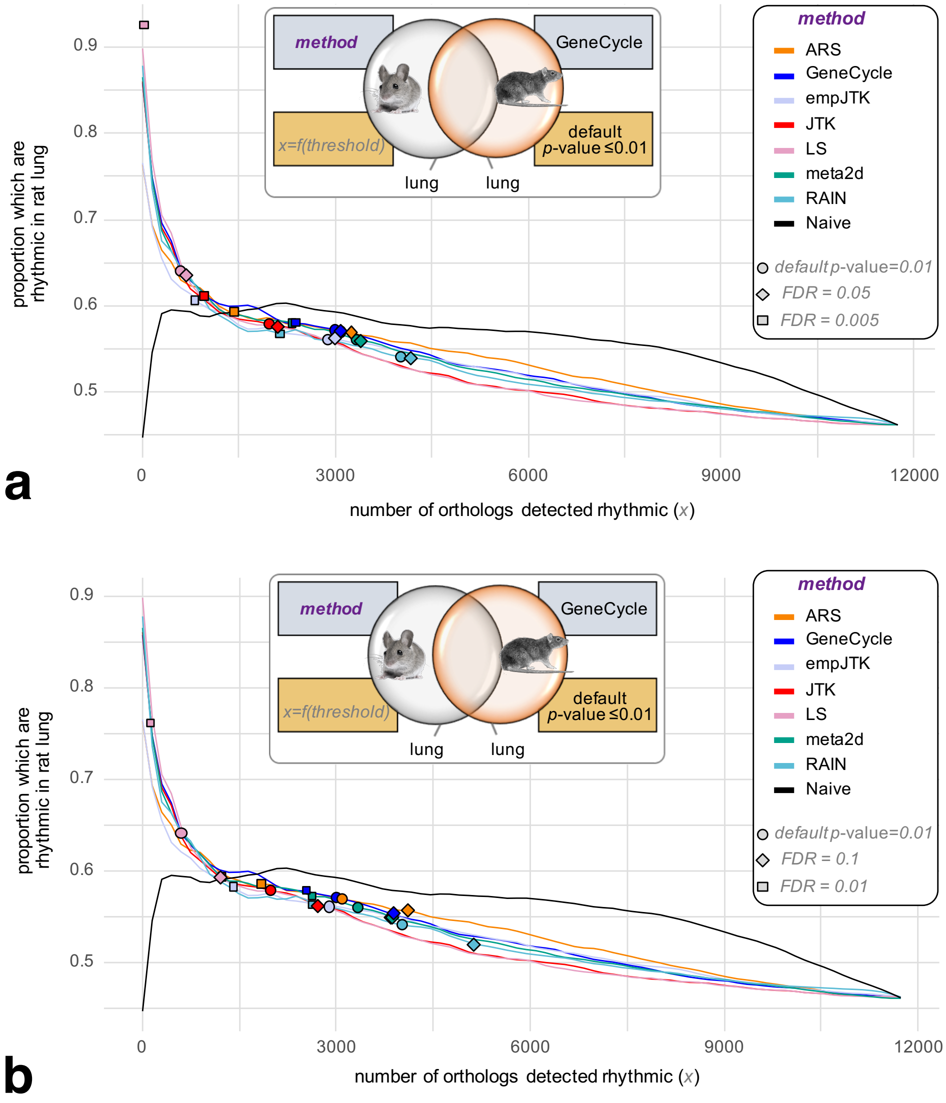


**Fig S15.**
Variation of the proportion rhythmic orthologs/all orthologs in mouse lung as a function of the number of mouse orthologs detected rhythmic, for each method applied to the mouse lung dataset. The benchmark gene set is composed of mouse-rat orthologs, detected rhythmic in rat lung by the GeneCycle method with default $p$-value $\leq 0.01$. The black line is the Naive method which orders genes according to their median expression levels (median of time-points), from highest expressed to lowest expressed gene, then, for each gene, calculates the proportion of rhythmic orthologs among those with higher expression. Rings correspond to a $p$-value threshold of 0.01, diamonds correspond to a FDR threshold of 0.05 (**a**) or 0.1 (**b**), and squares correspond to a FDR threshold of 0.005 (**a**) or 0.01 (**b**). FDR correspond to the false discovery rate adjustment of default $p$-values using `p.adjust` R function.

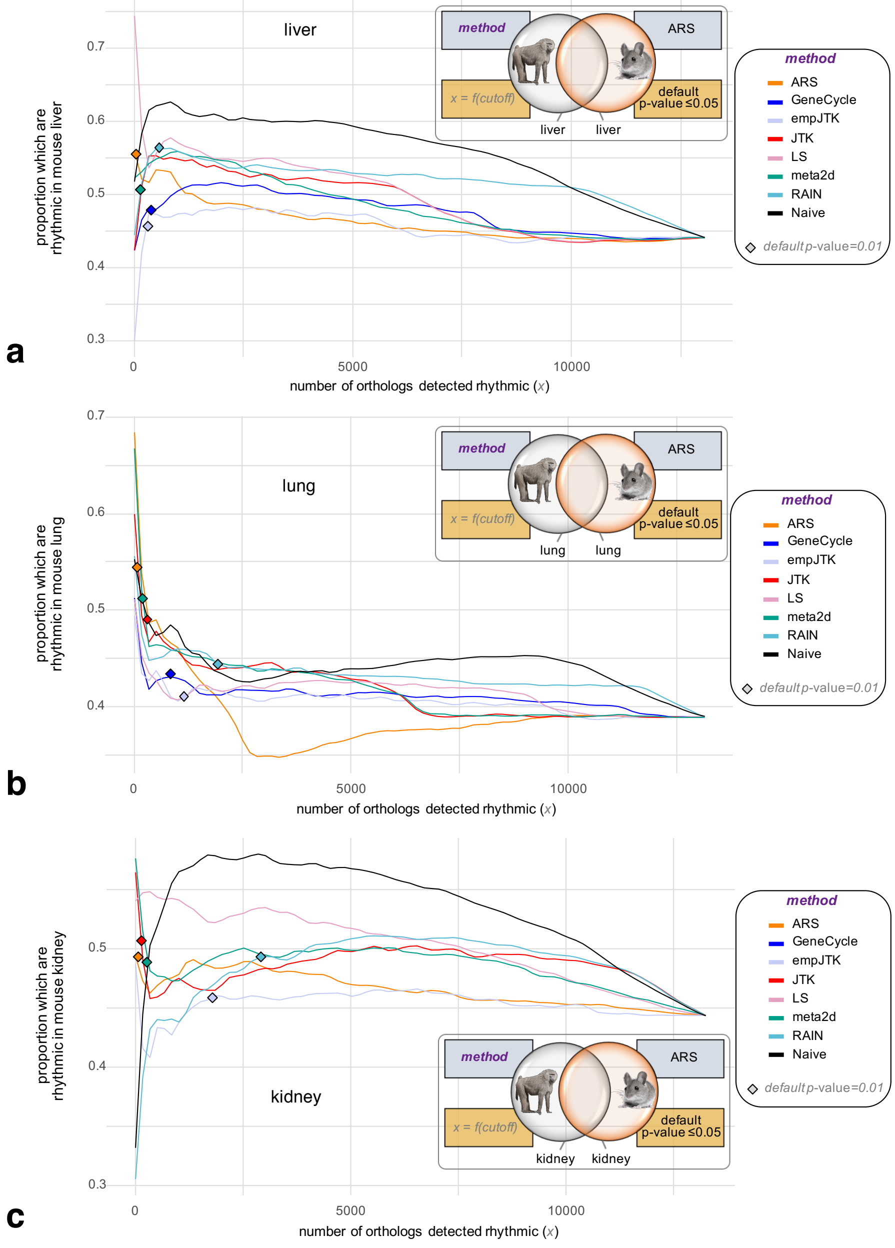


**Fig S16.** Variation of the proportion rhythmic orthologs/all orthologs in baboon as a function of the number of mouse orthologs detected rhythmic, for each method applied to the baboon lung (**a**), lung (**b**), and kidney (**c**) dataset. The benchmark gene set is composed of baboon-mouse orthologs, detected rhythmic in the homologous tissue of mouse by the ARS method with default $p$-value $\leq 0.05$.

### References


1. S. S. C. Rund, T. Y. Hou, S. M. Ward, F. H. Collins, and G. E. Duffield. Genome-wide profiling of diel and circadian gene expression in the malaria vector anopheles gambiae. *Proceedings of the National Academy of Sciences*, 108(32):E421–E430, 2011.

2. R. Zhang, N. F. Lahens, H. I. Ballance, M. E. Hughes, and J. B. Hogenesch. A circadian gene expression atlas in

mammals: Implications for biology and medicine. *Proceedings of the National Academy of Sciences*, 111(45):16219–16224, 2014.
