## Additional file 2 for "Methods detecting rhythmic gene expression are biologically relevant only for strong signal"

### VERTEBRATES

#### default *p*-values

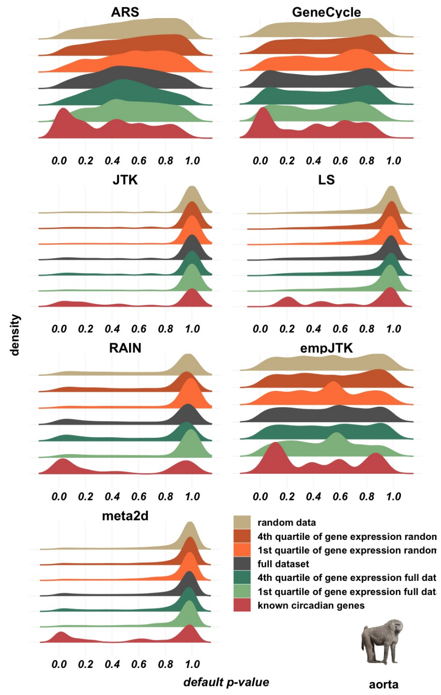


**Fig. S1**

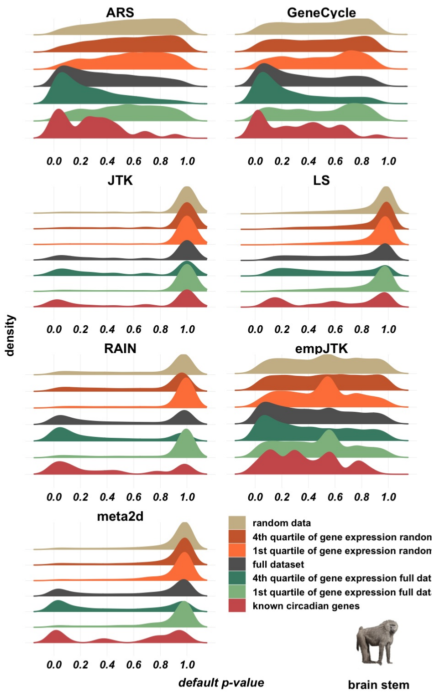

ARS
GeneCycle
JTK
LS
RAIN
empJTK
meta2d
0.0 0.2 0.4 0.6 0.8 1.0
density
default p-value
random data
4th quartile of gene expression randon
1st quartile of gene expression randon
full dataset
4th quartile of gene expression full dat
1st quartile of gene expression full dat
known circadian genes
brain stem


Fig. S2

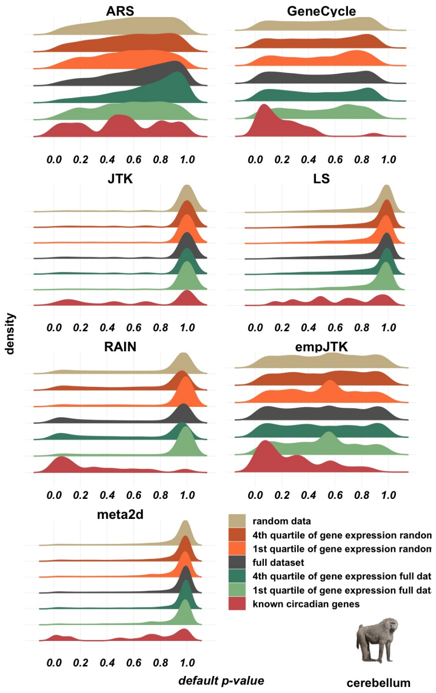

ARS
GeneCycle
JTK
LS
RAIN
empJTK
meta2d
0.0 0.2 0.4 0.6 0.8 1.0
density
default p-value
random data
4th quartile of gene expression randon
1st quartile of gene expression randon
full dataset
4th quartile of gene expression full dat
1st quartile of gene expression full dat
known circadian genes
cerebellum


Fig. S3

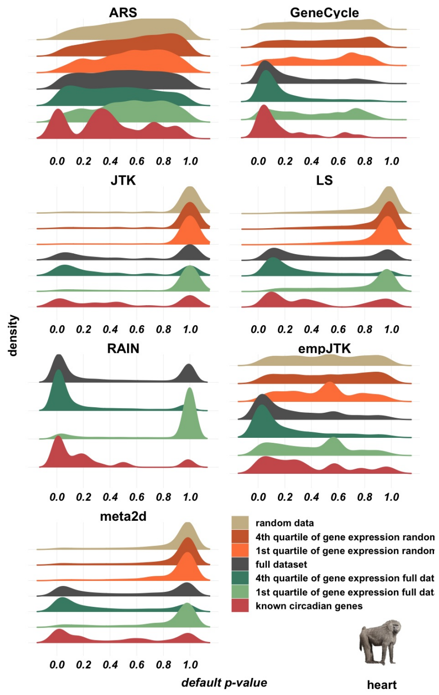
ARS
GeneCycle
JTK
LS
RAIN
empJTK
meta2d
density
0.0 0.2 0.4 0.6 0.8 1.0
random data
4th quartile of gene expression randon
1st quartile of gene expression randon
full dataset
4th quartile of gene expression full dat
1st quartile of gene expression full dat
known circadian genes
default p-value
heart

### Fig. S4

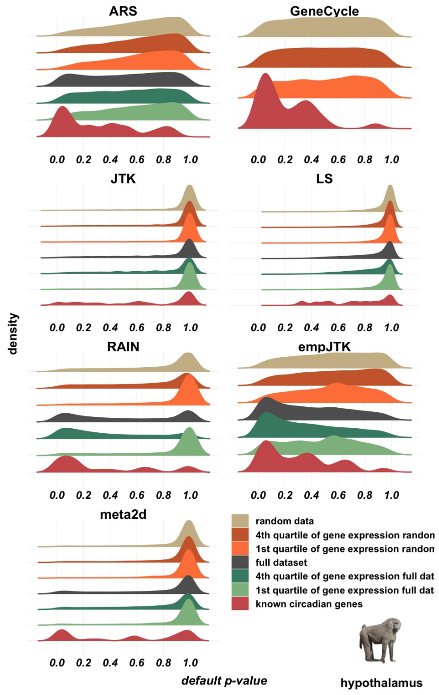
ARS
GeneCycle
JTK
LS
RAIN
empJTK
meta2d
density
0.0 0.2 0.4 0.6 0.8 1.0
random data
4th quartile of gene expression randon
1st quartile of gene expression randon
full dataset
4th quartile of gene expression full dat
1st quartile of gene expression full dat
known circadian genes
default p-value
hypothalamus

### Fig. S5

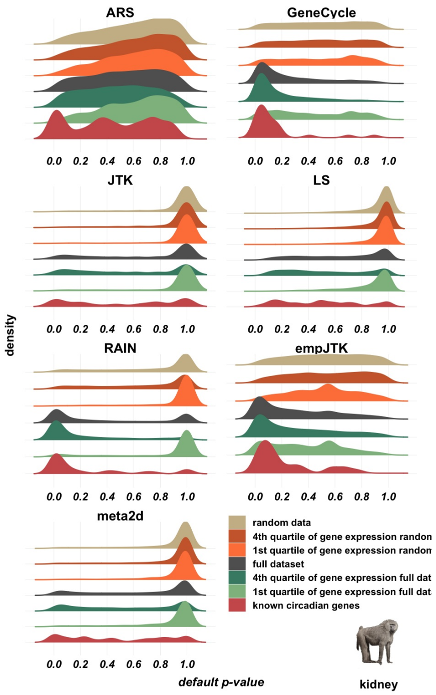
ARS
GeneCycle
JTK
LS
RAIN
empJTK
meta2d
density
0.0 0.2 0.4 0.6 0.8 1.0
default p-value
random data
4th quartile of gene expression randon
1st quartile of gene expression randon
full dataset
4th quartile of gene expression full dat
1st quartile of gene expression full dat
known circadian genes
kidney

**Fig. S6**

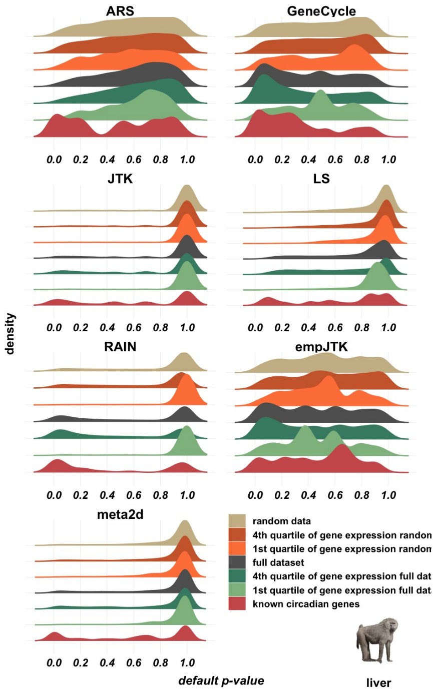
ARS
GeneCycle
JTK
LS
RAIN
empJTK
meta2d
density
0.0 0.2 0.4 0.6 0.8 1.0
default p-value
random data
4th quartile of gene expression randon
1st quartile of gene expression randon
full dataset
4th quartile of gene expression full dat
1st quartile of gene expression full dat
known circadian genes
liver

**Fig. S7**

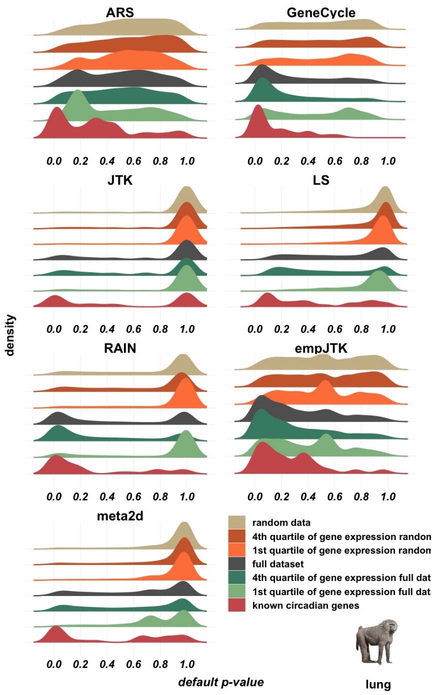
ARS
GeneCycle
JTK
LS
RAIN
empJTK
meta2d
0.0 0.2 0.4 0.6 0.8 1.0
density
default p-value
random data
4th quartile of gene expression randon
1st quartile of gene expression randon
full dataset
4th quartile of gene expression full dat
1st quartile of gene expression full dat
known circadian genes
lung

Fig. S8

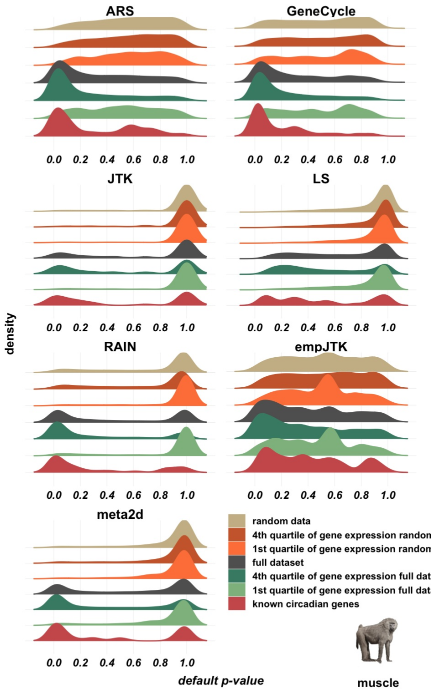
ARS
GeneCycle
JTK
LS
RAIN
empJTK
meta2d
0.0 0.2 0.4 0.6 0.8 1.0
density
default p-value
random data
4th quartile of gene expression randon
1st quartile of gene expression randon
full dataset
4th quartile of gene expression full dat
1st quartile of gene expression full dat
known circadian genes
muscle

Fig. S9

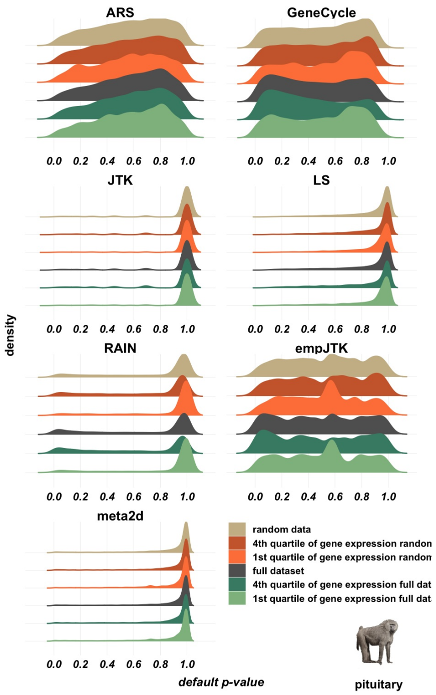


### Fig. S10

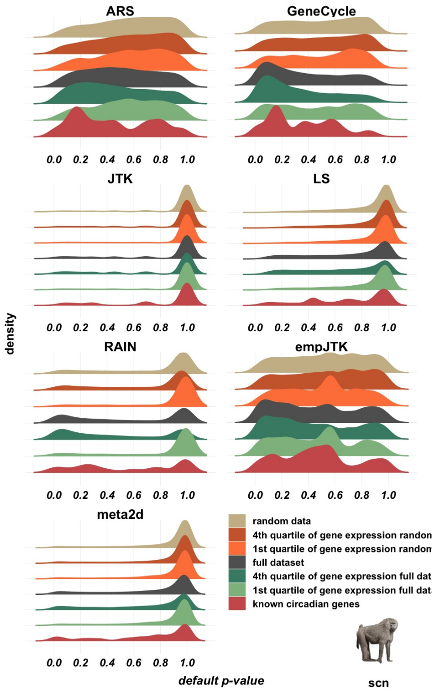


### Fig. S11

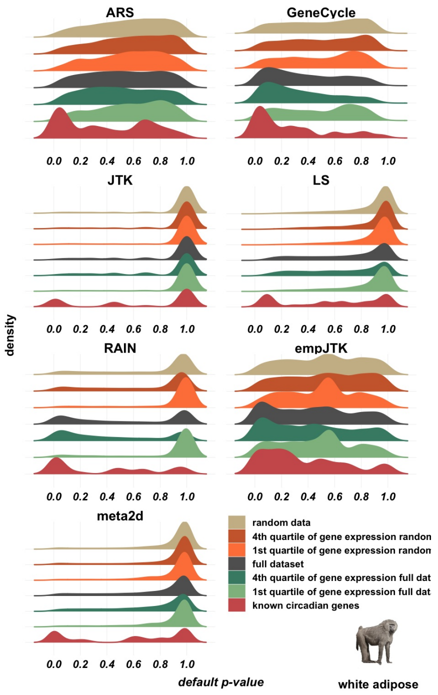


### Fig. S12

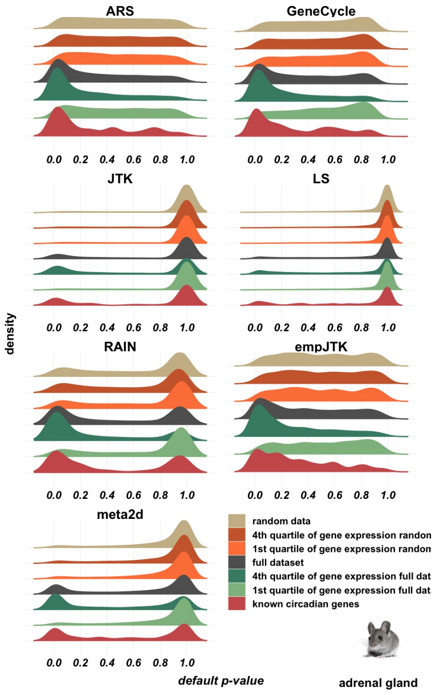


### Fig. S13

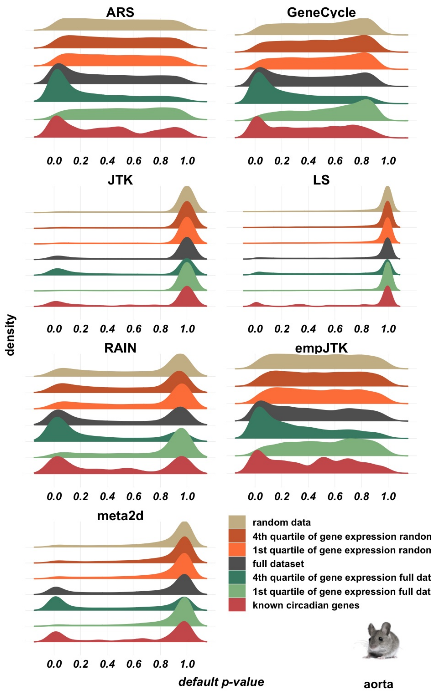


### Fig. S14

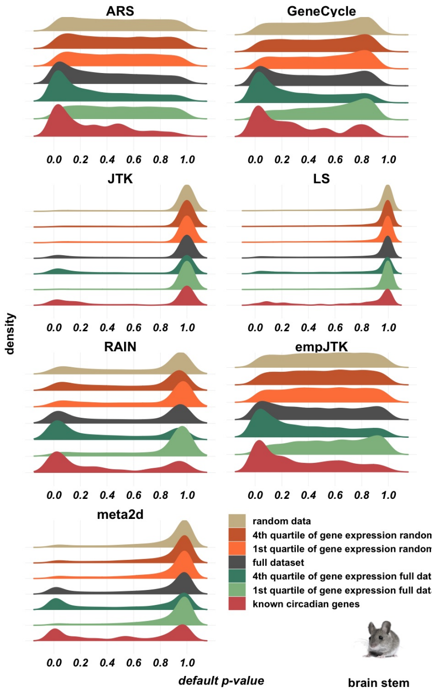


### Fig. S15

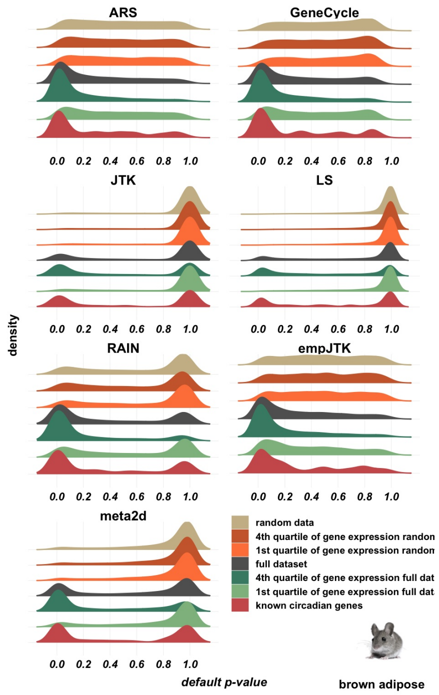


### Fig. S16

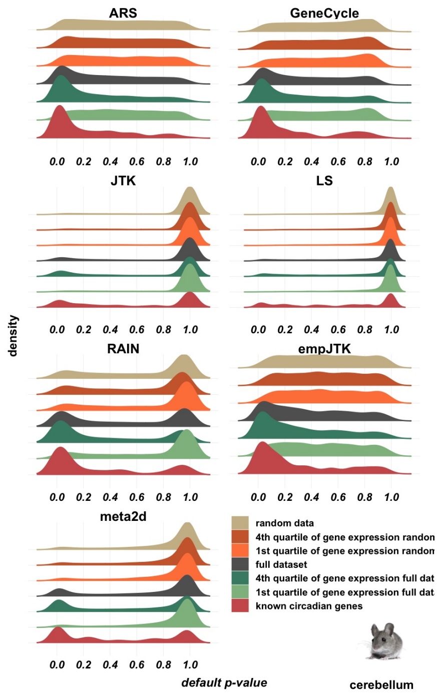


### Fig. S17

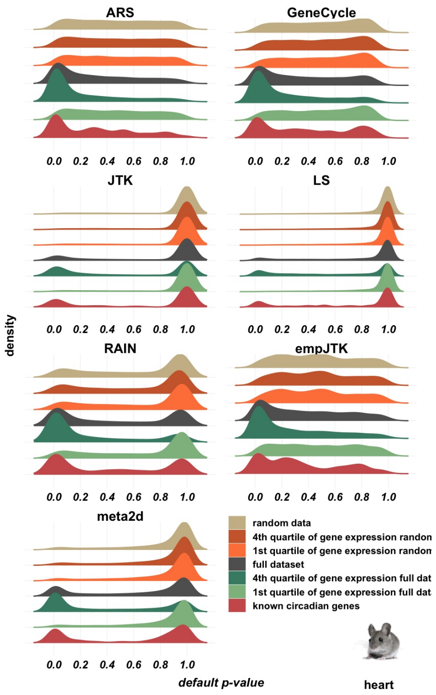


### Fig. S18

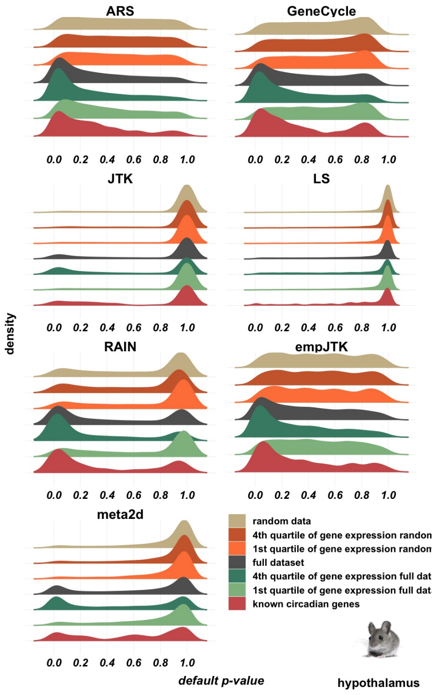


### Fig. S19

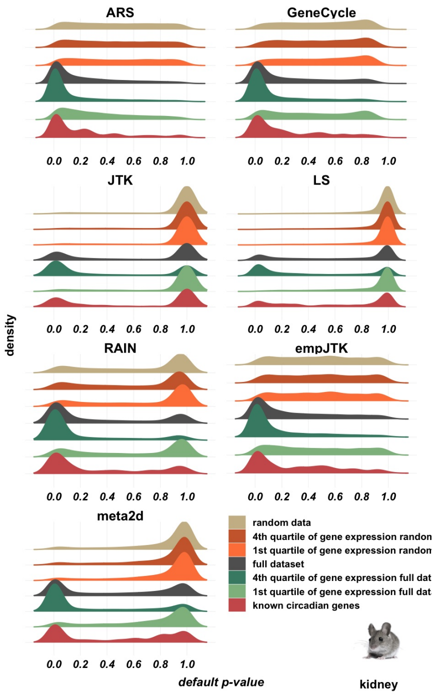


### Fig. S20

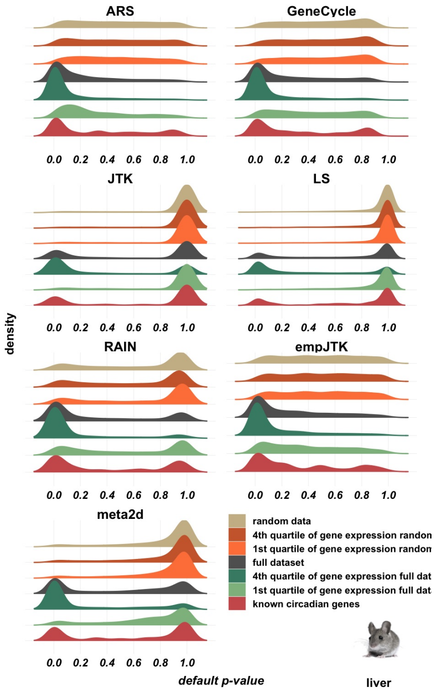


### Fig. S21

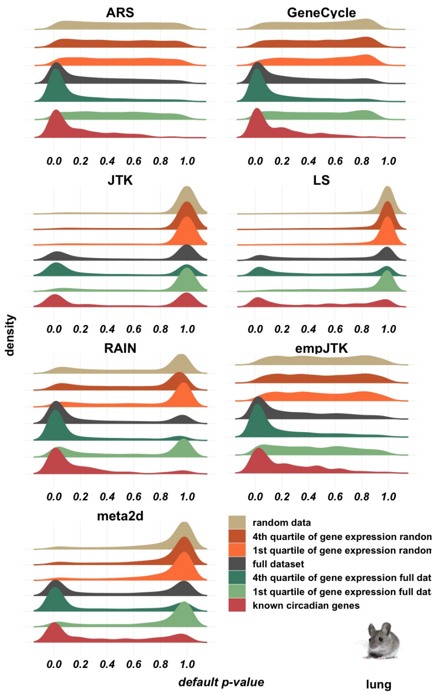


### Fig. S22

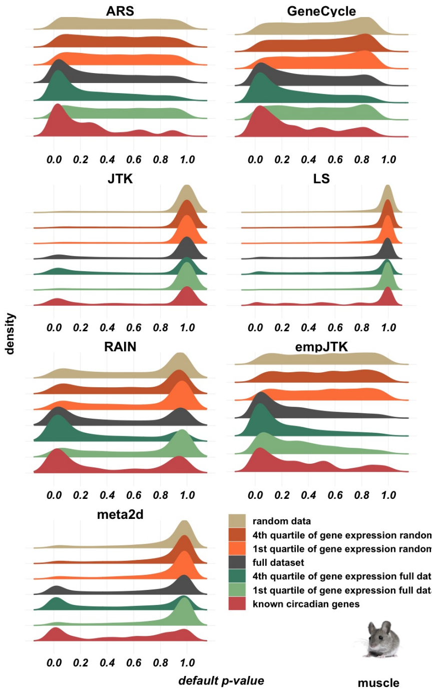


### Fig. S23

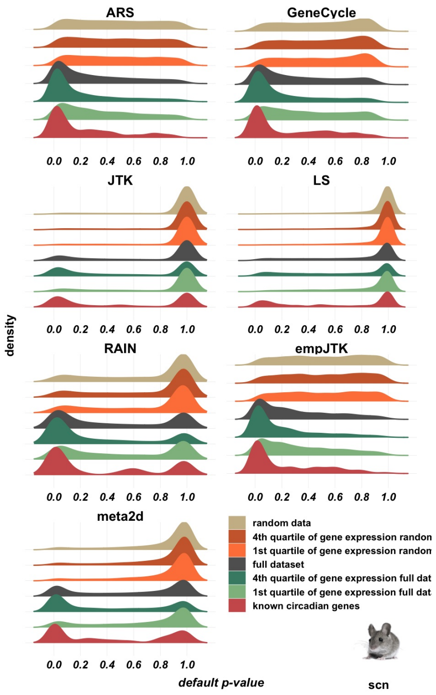


### Fig. S24

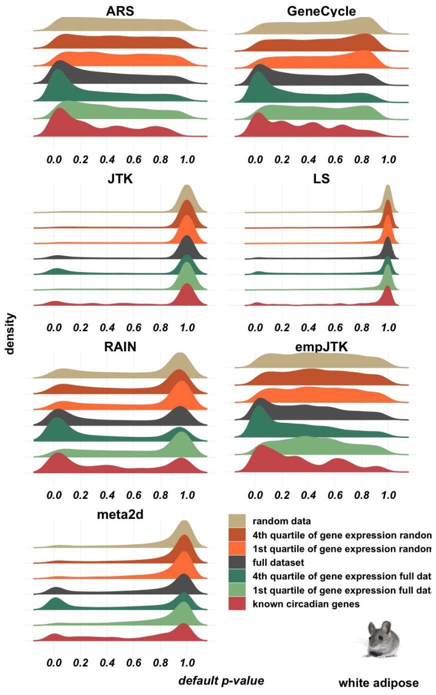


### Fig. S25

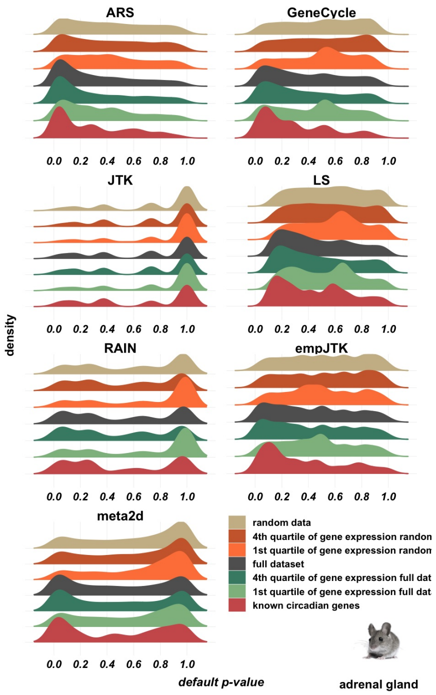


### Fig. S26

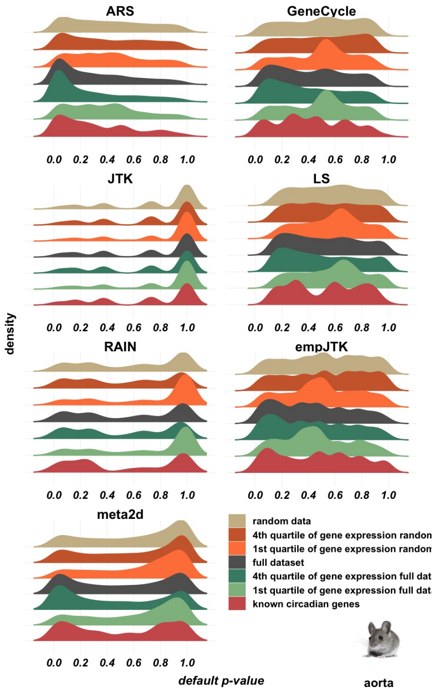


### Fig. S27

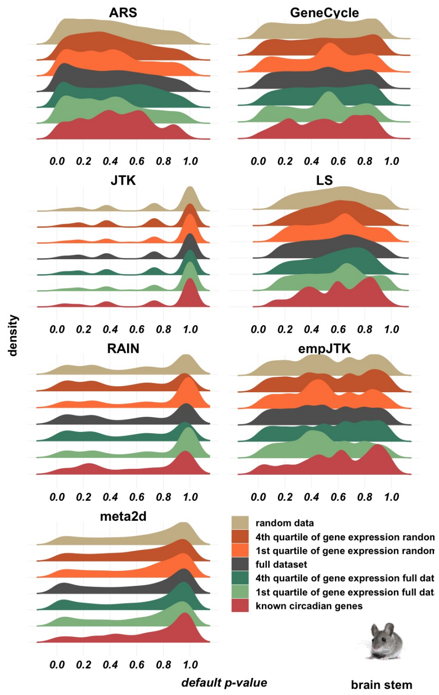


### Fig. S28

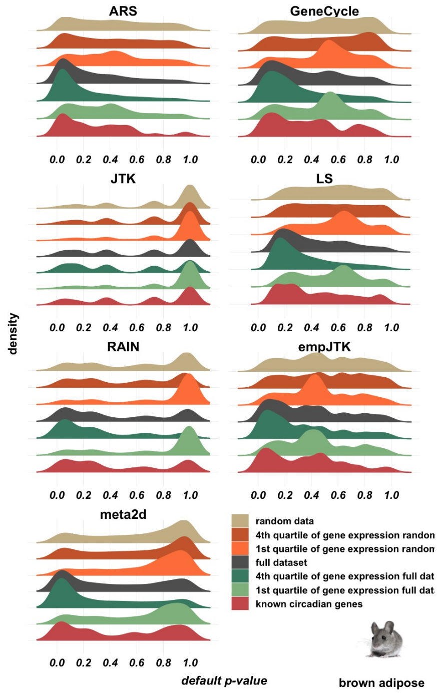


### Fig. S29

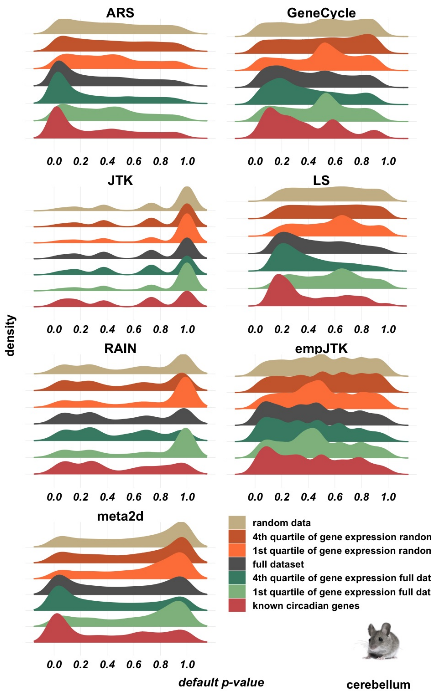


**Fig. S30**

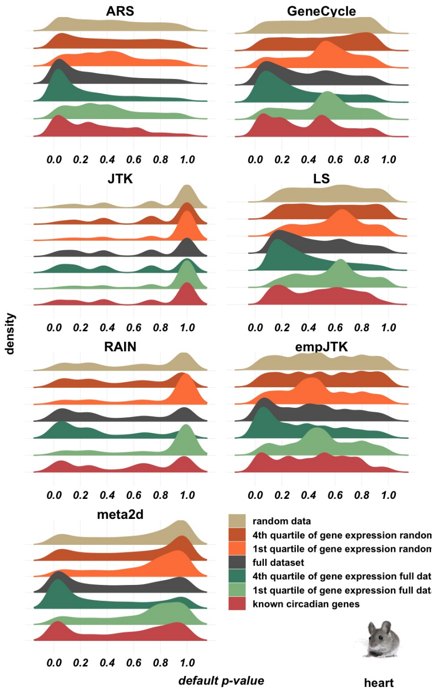


**Fig. S31**

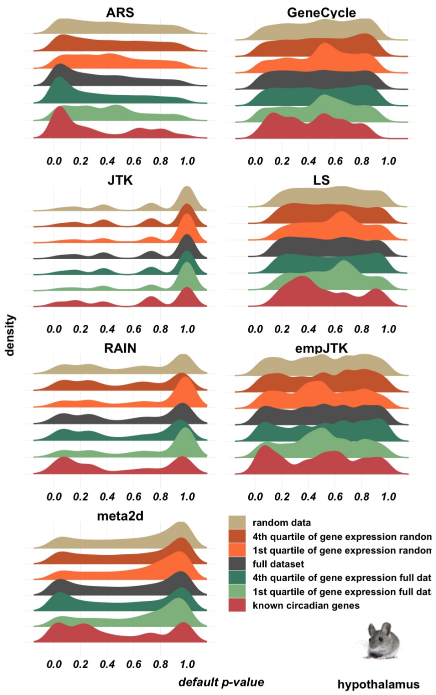


**Fig. S32**

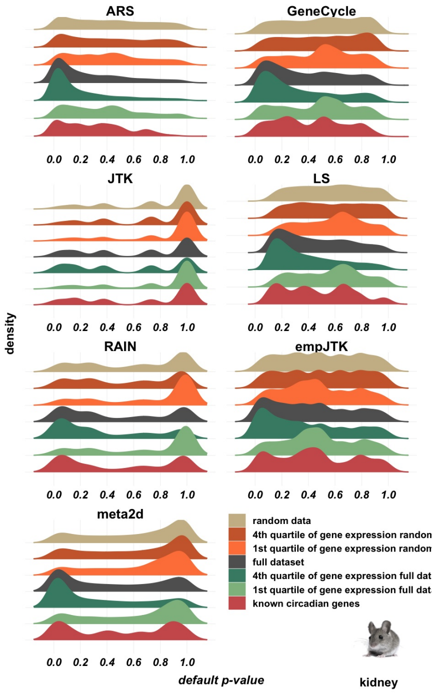


**Fig. S33**

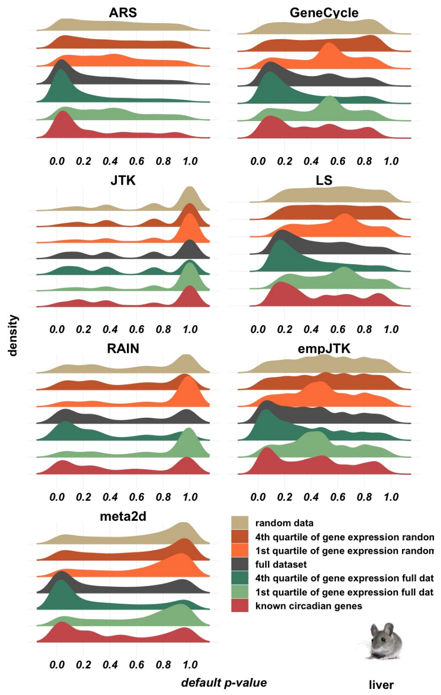


### Fig. S34

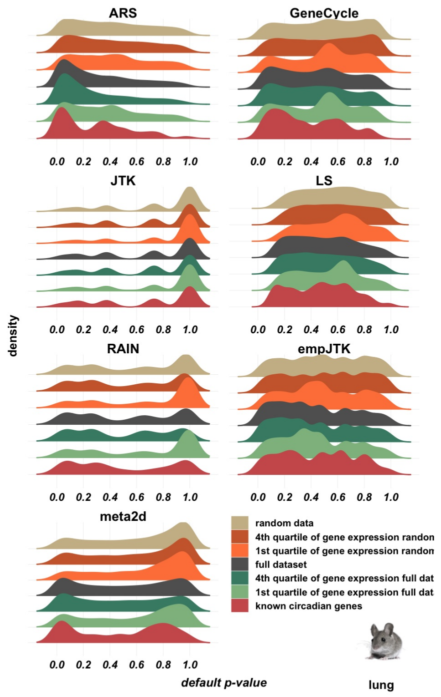


### Fig. S35

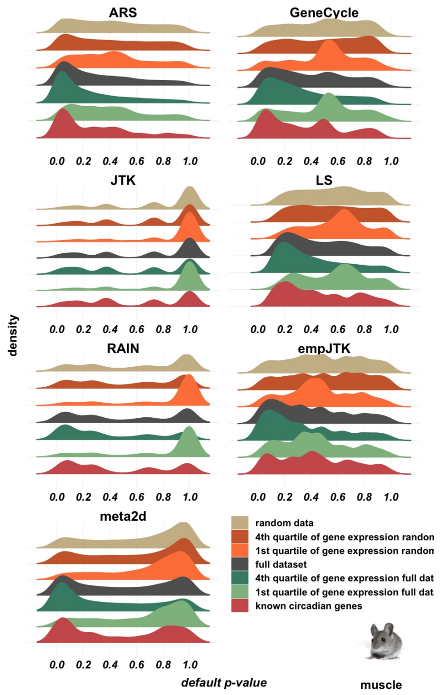


### Fig. S36

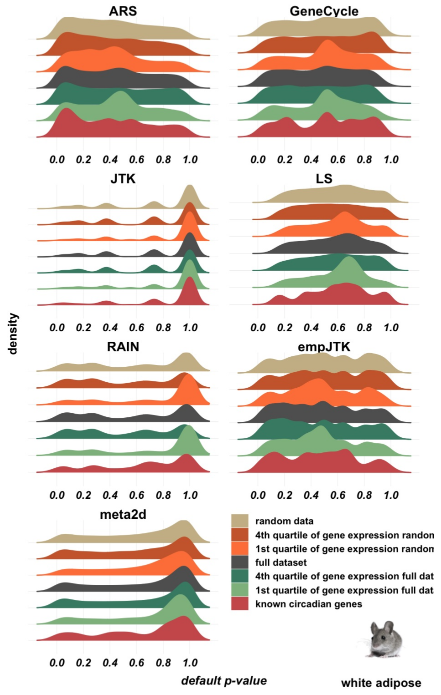


### Fig. S37

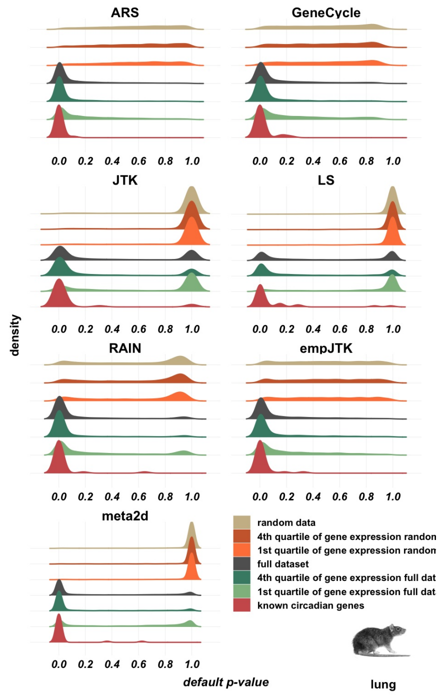


### Fig. S38

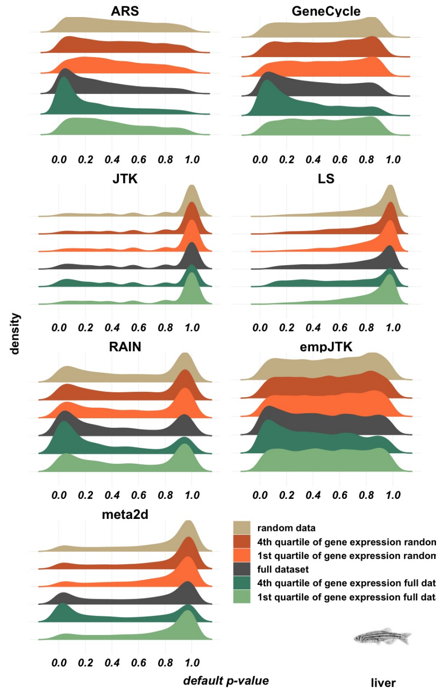


### Fig. S39

### raw *p*-values

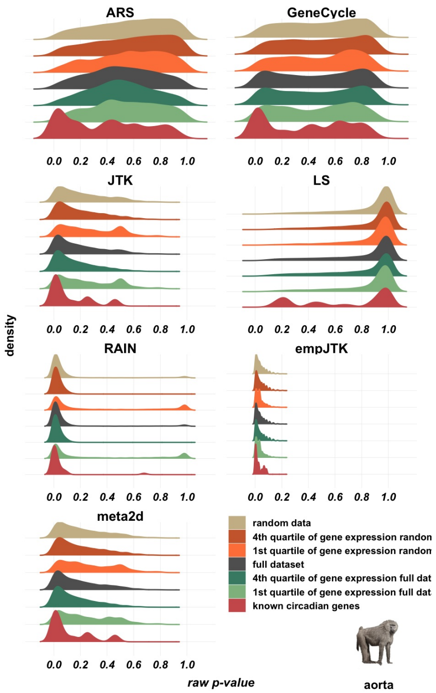


### Fig. S40

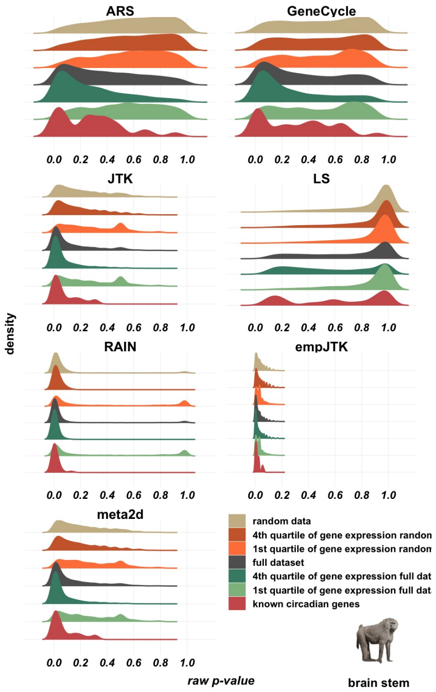


### Fig. S41

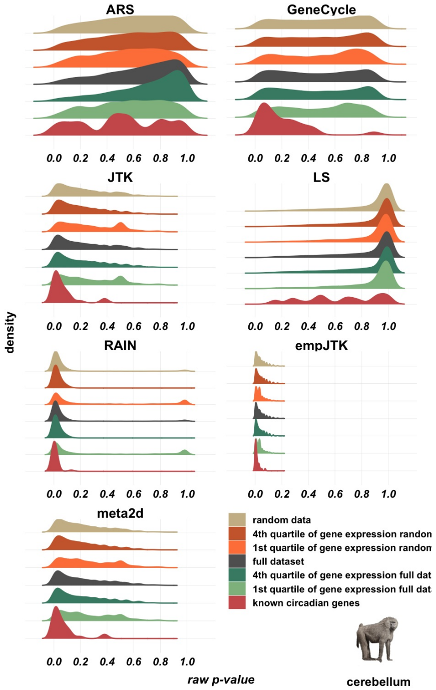


### Fig. S42

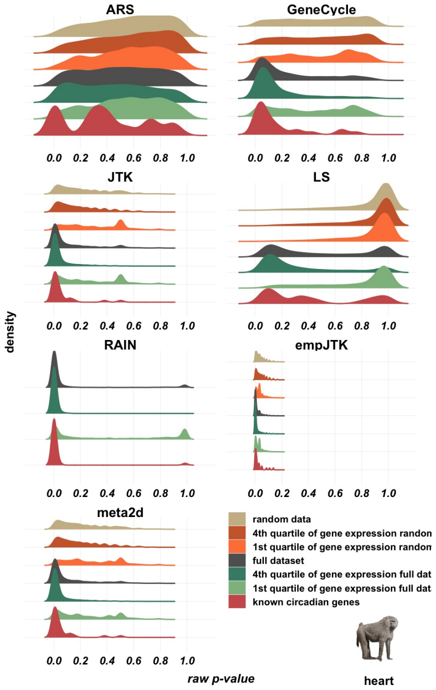


### Fig. S43

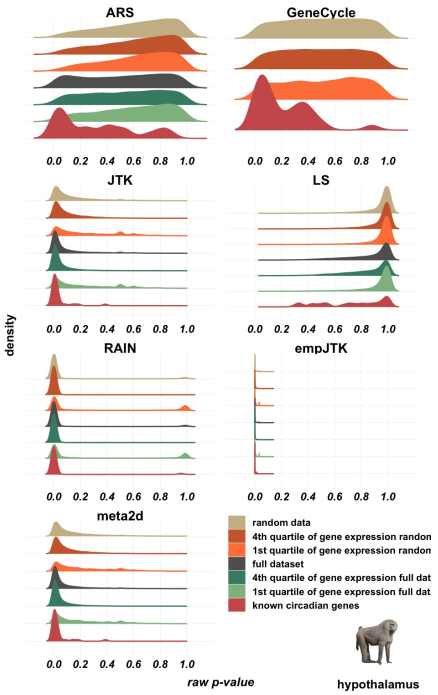


### Fig. S44

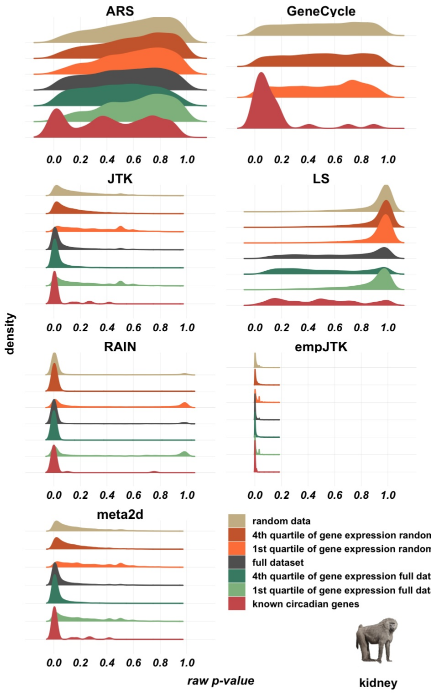


### Fig. S45

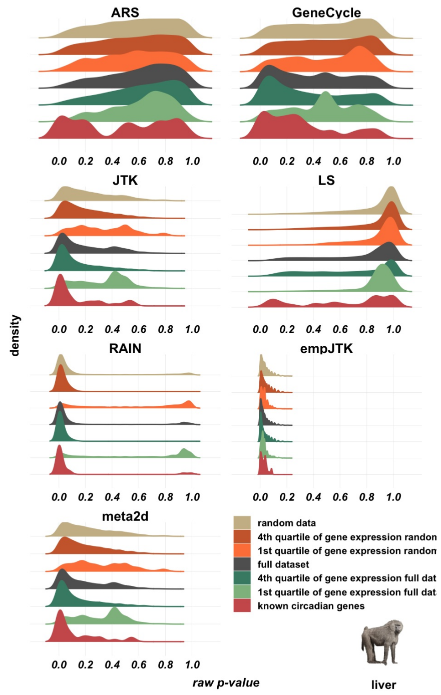


### Fig. S46

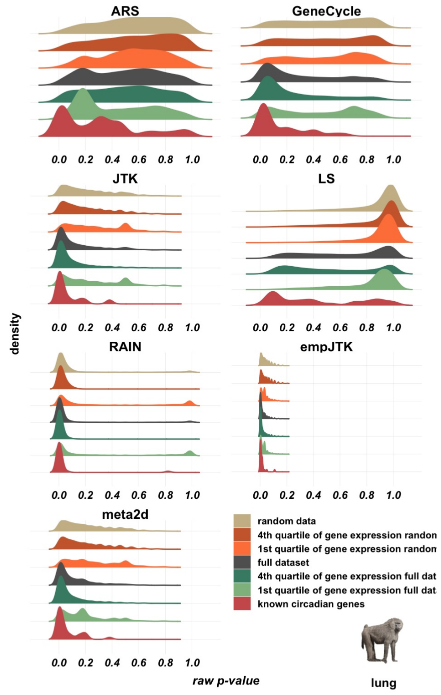


### Fig. S47

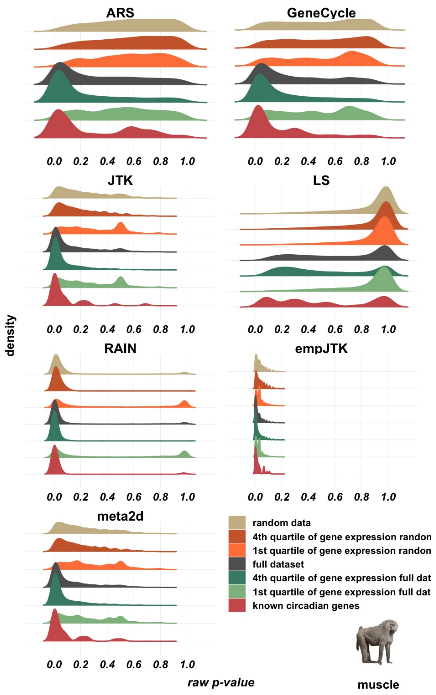


### Fig. S48

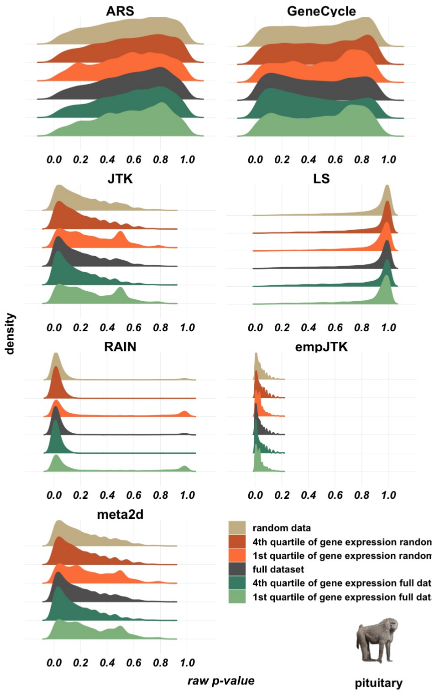


### Fig. S49

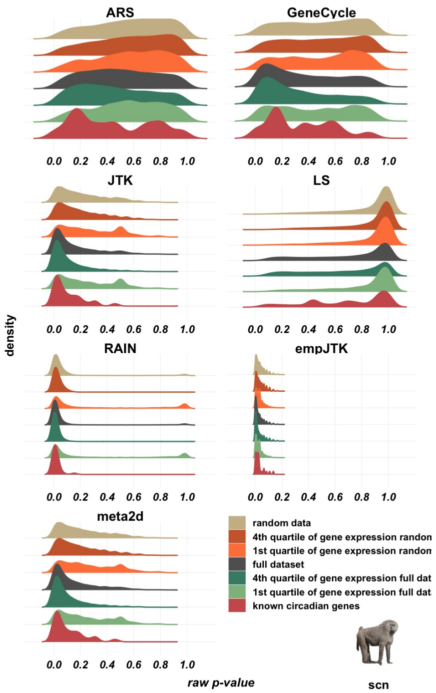


### Fig. S50

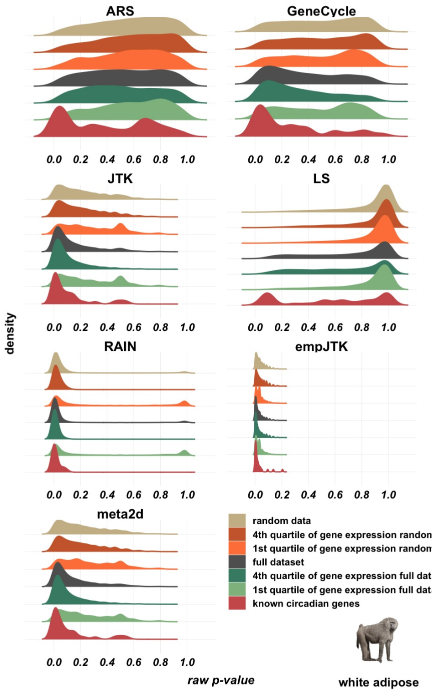


### Fig. S51

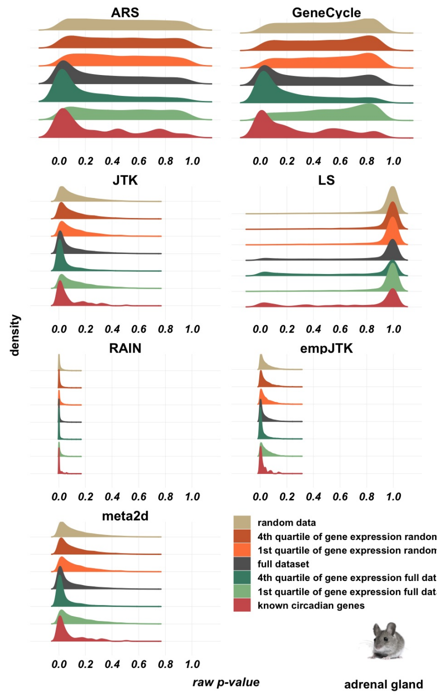


**Fig. S52**

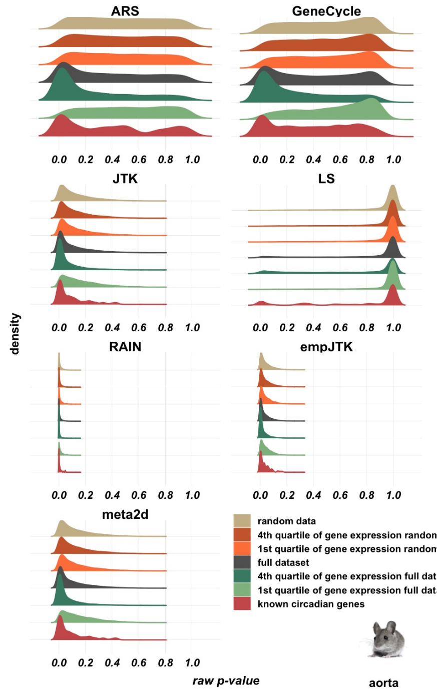


**Fig. S53**

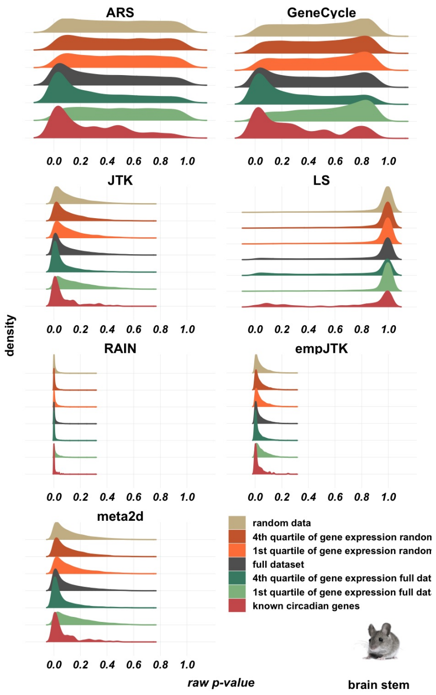


### Fig. S54

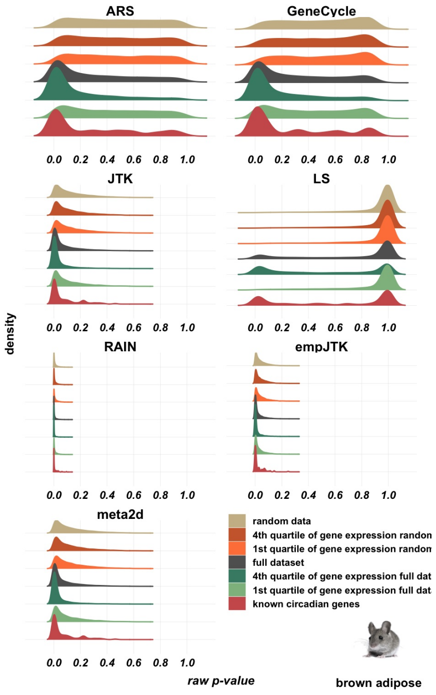


### Fig. S55

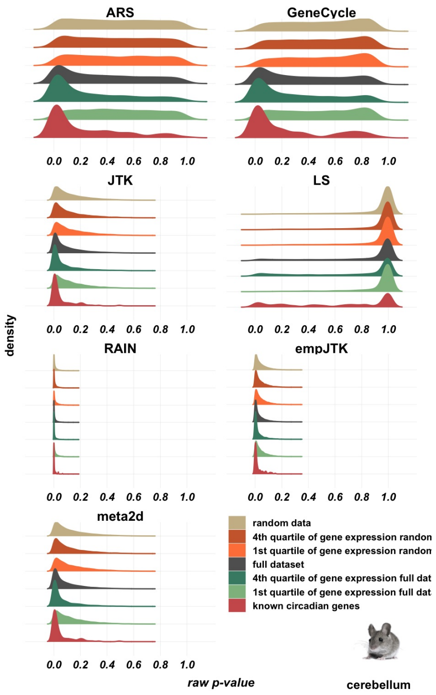


### Fig. S56

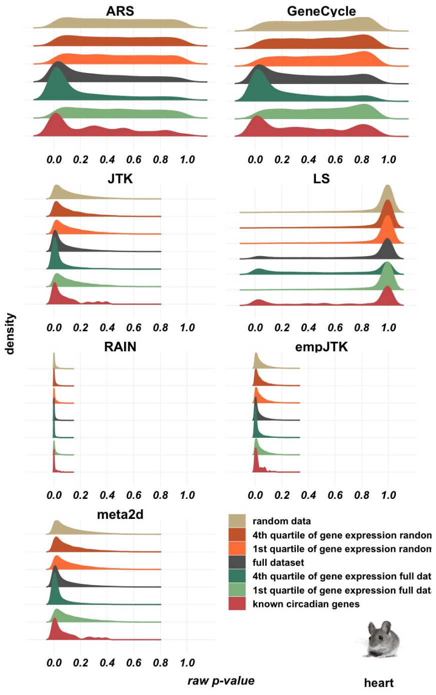


### Fig. S57

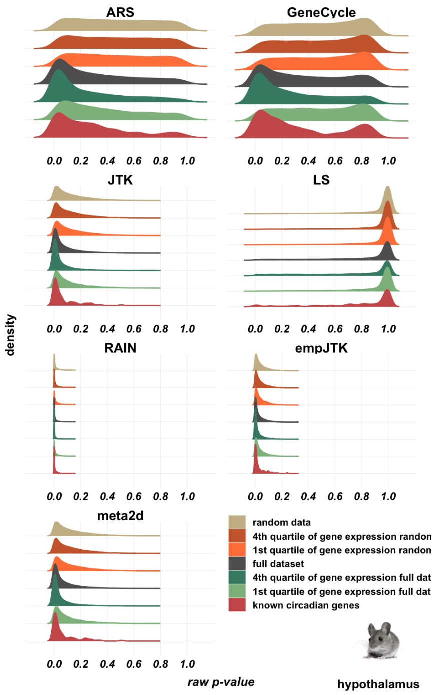


### Fig. S58

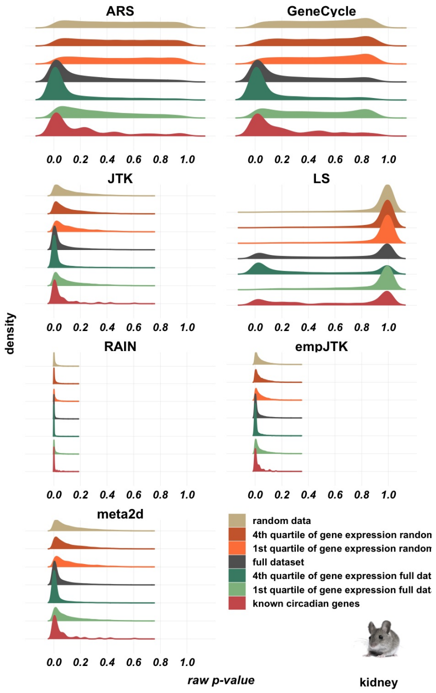


### Fig. S59

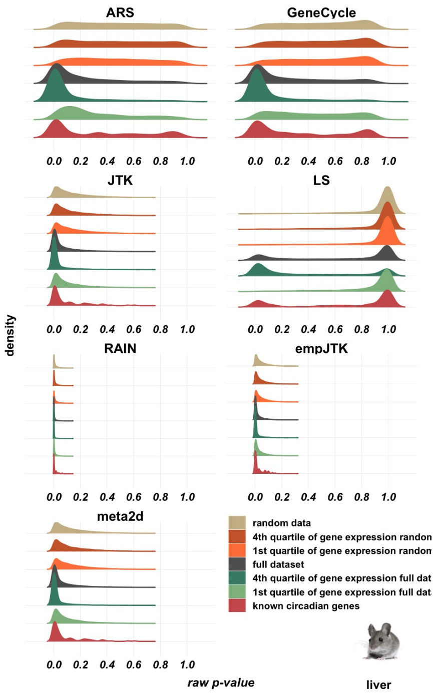


### Fig. S60

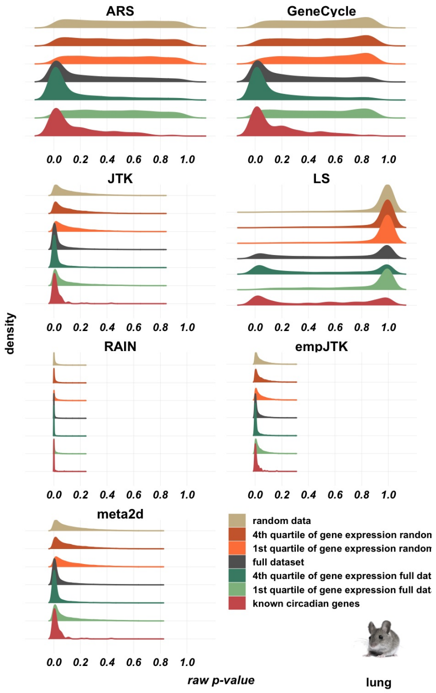


### Fig. S61

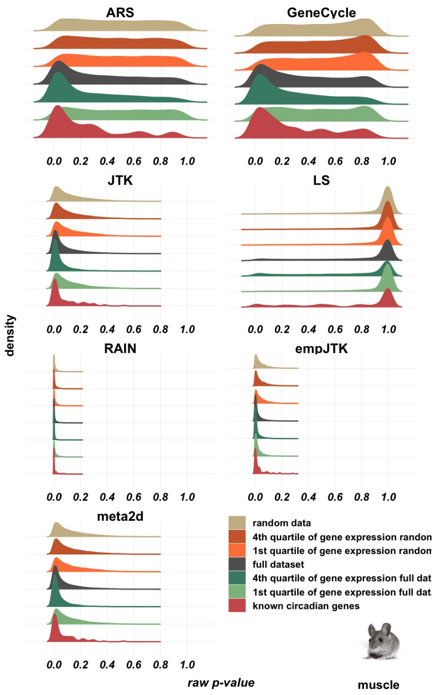


**Fig. S62**

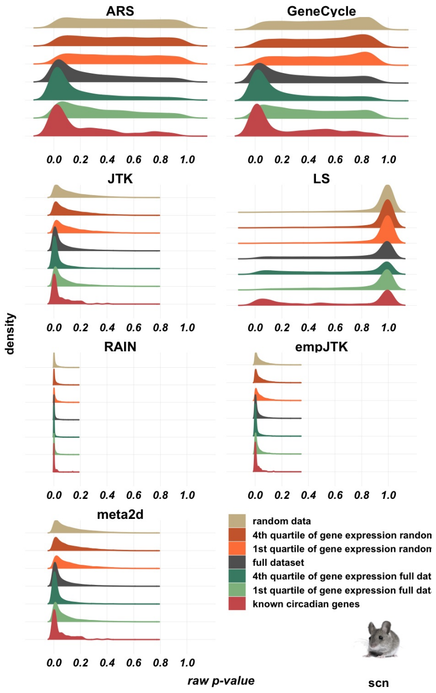


**Fig. S63**

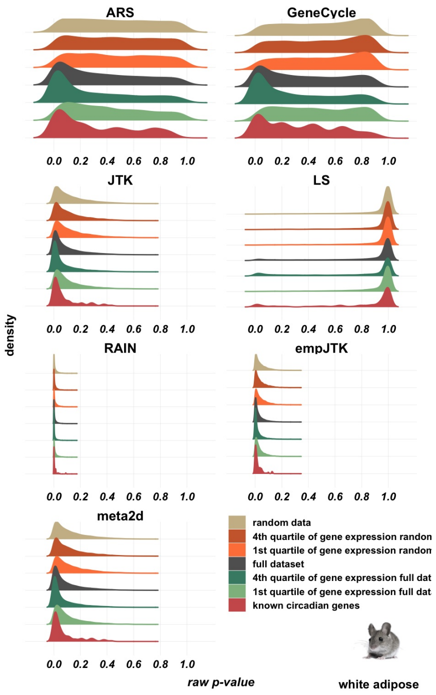


### Fig. S64

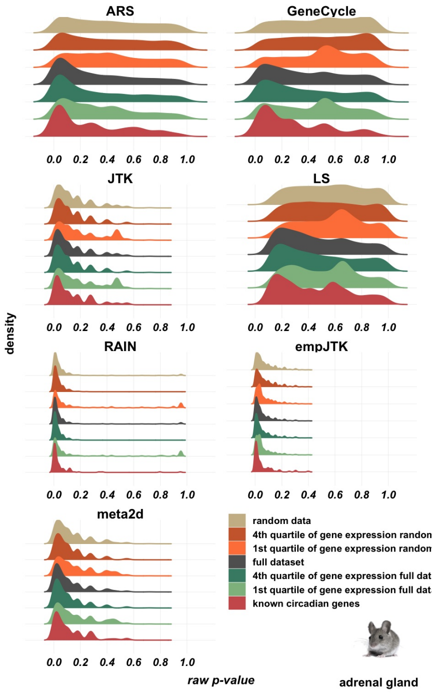


### Fig. S65

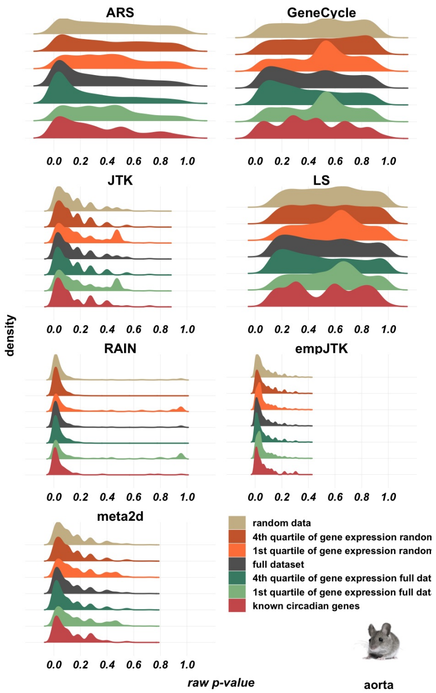


### Fig. S66

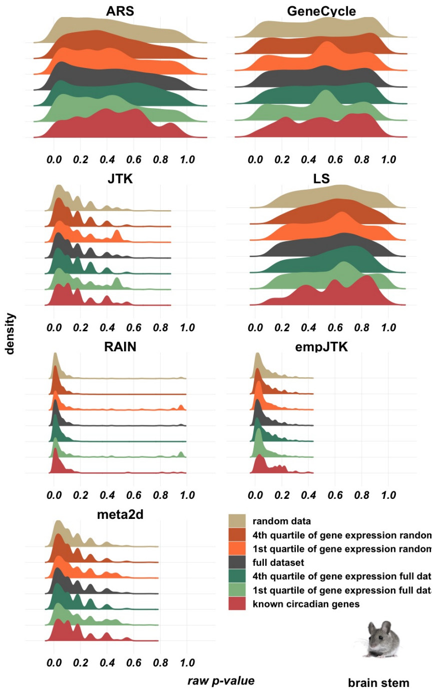


### Fig. S67

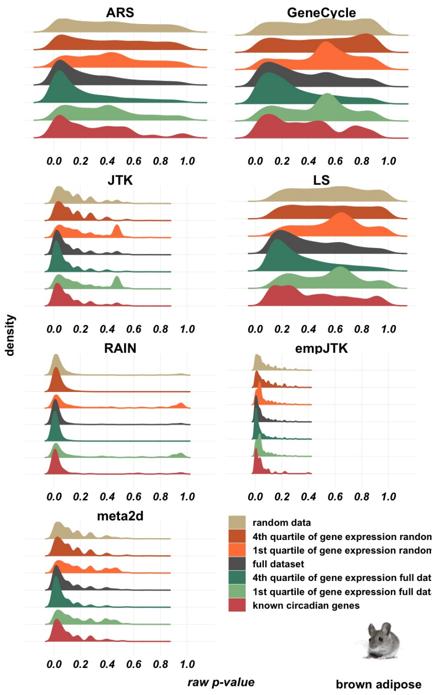


### Fig. S68

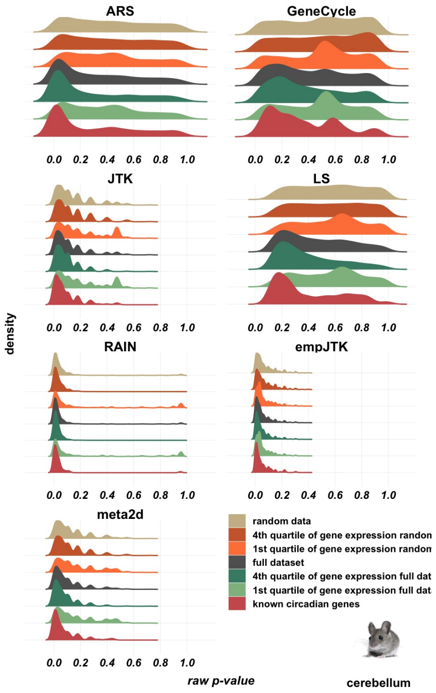


### Fig. S69

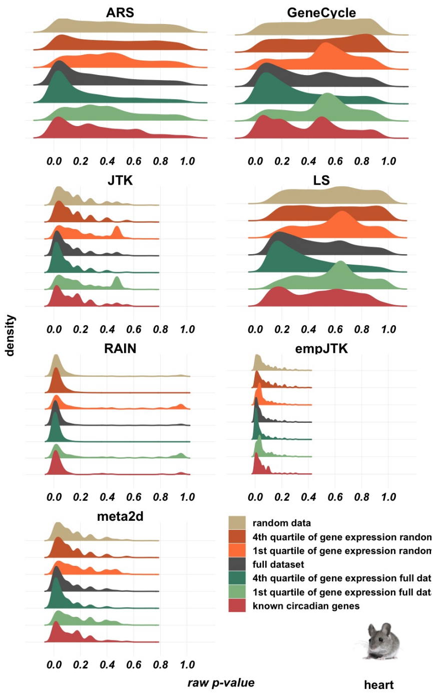


### Fig. S70

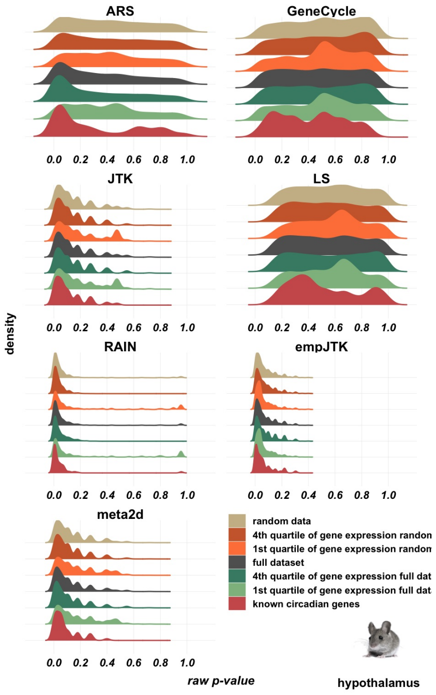


### Fig. S71

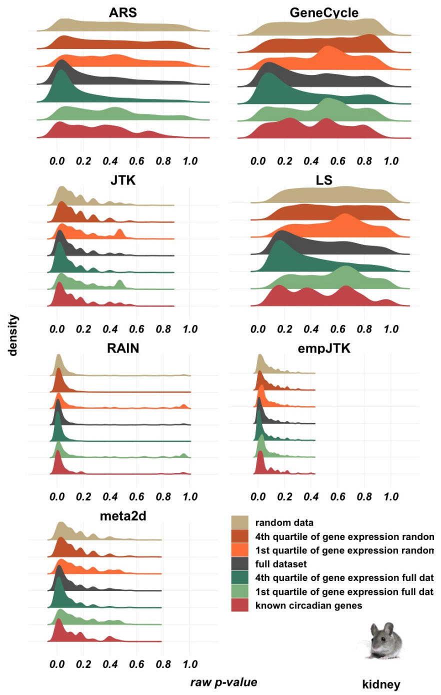


**Fig. S72**

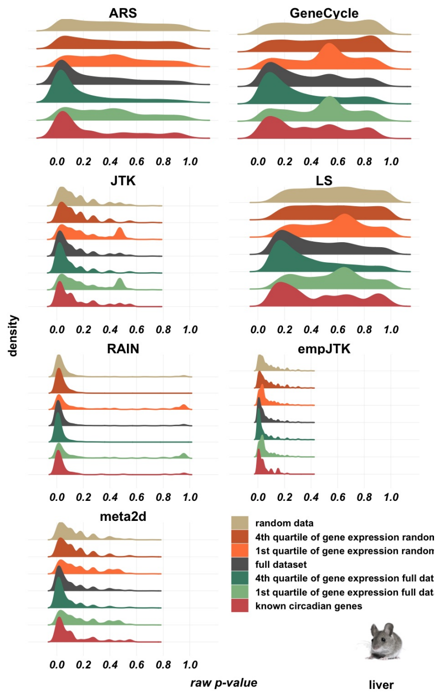


**Fig. S73**

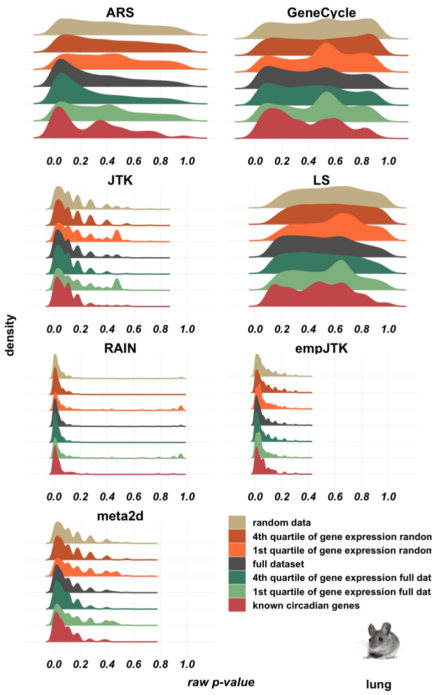


### Fig. S74

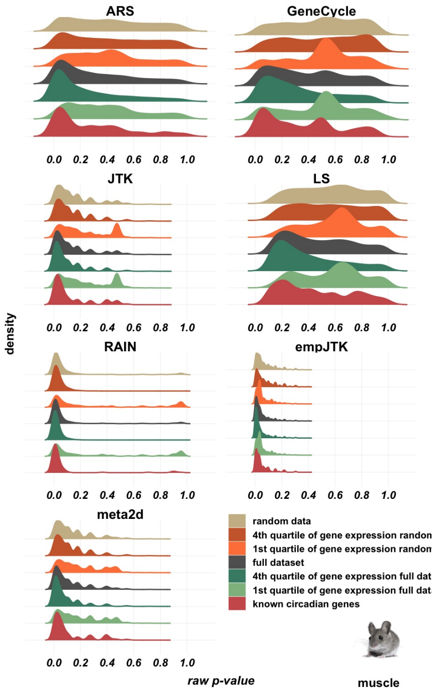


### Fig. S75

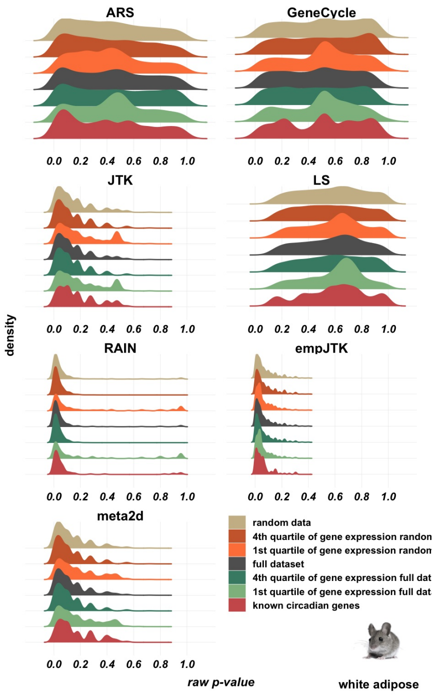


### Fig. S76

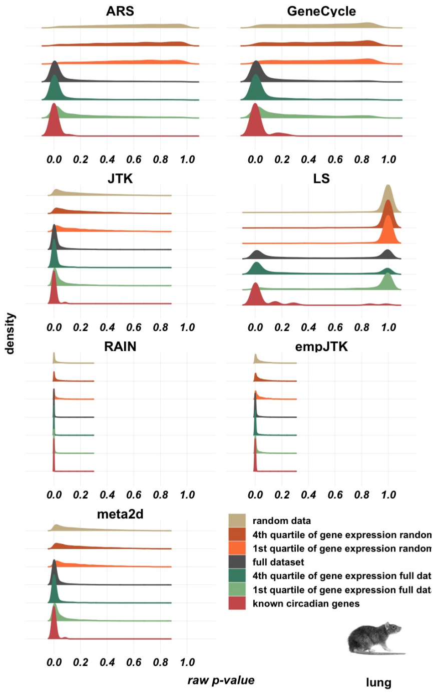


### Fig. S77

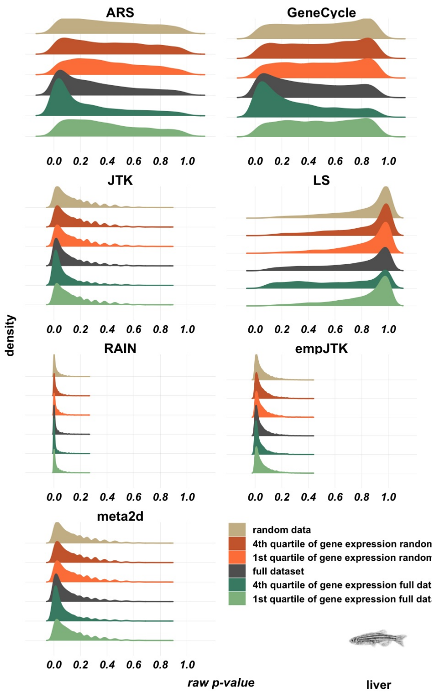


### Fig. S78

# BH.Q

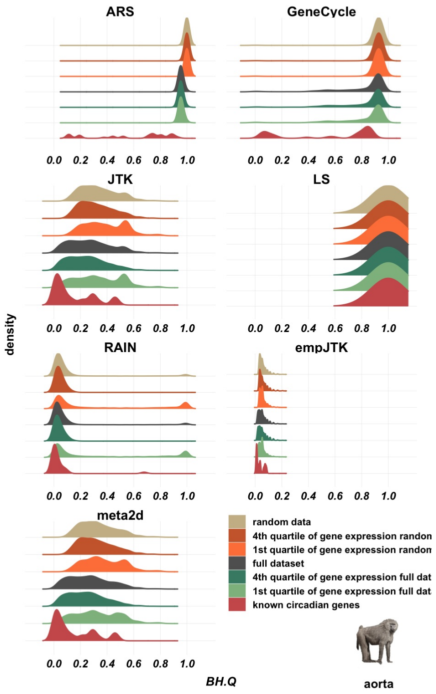


### Fig. S78

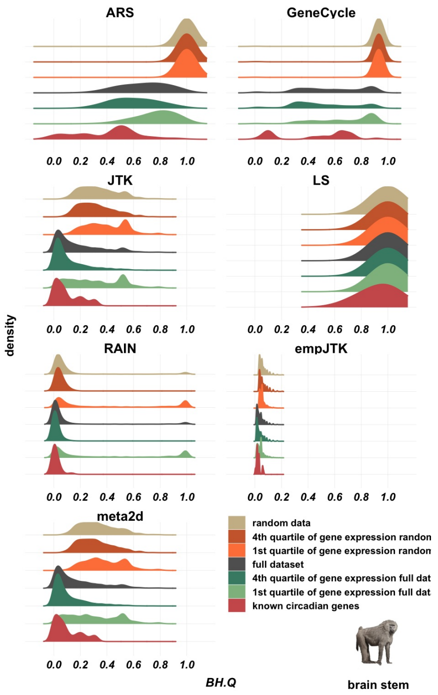


### Fig. S78

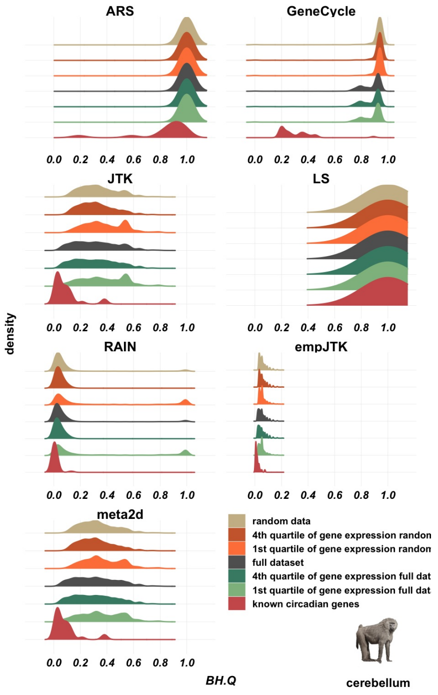


### Fig. S78

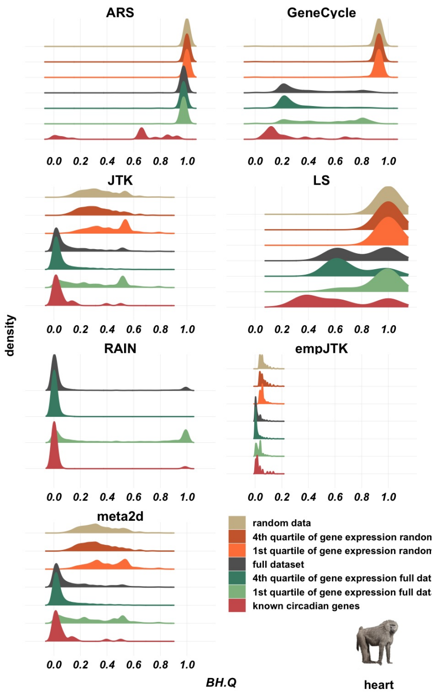


### Fig. S78

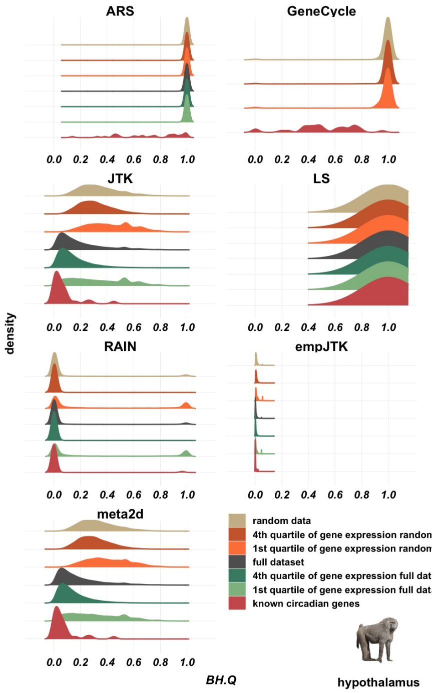


### Fig. S78

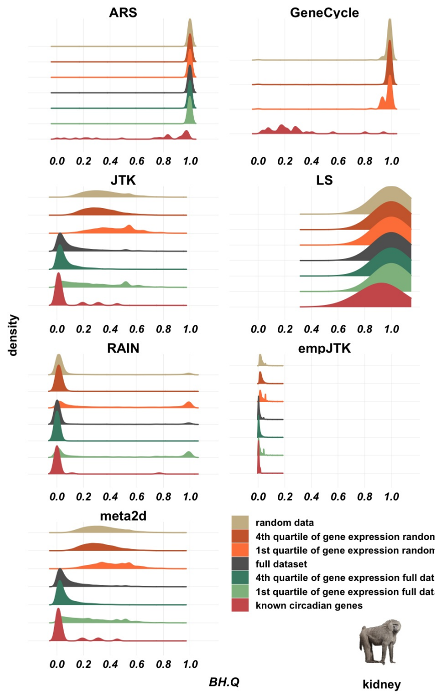


**Fig. S78**

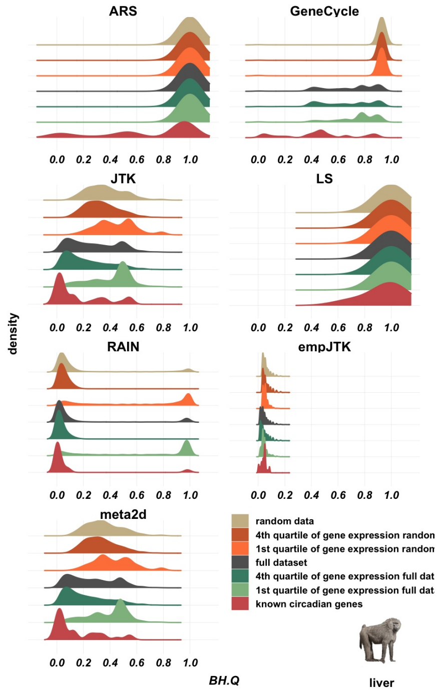


**Fig. S78**

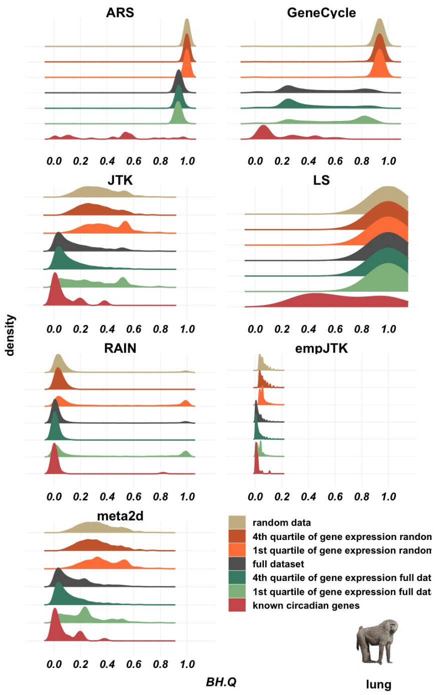


Fig. S78

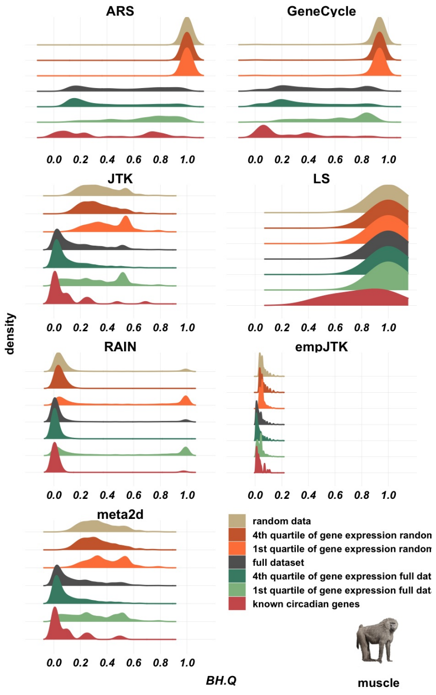


Fig. S78

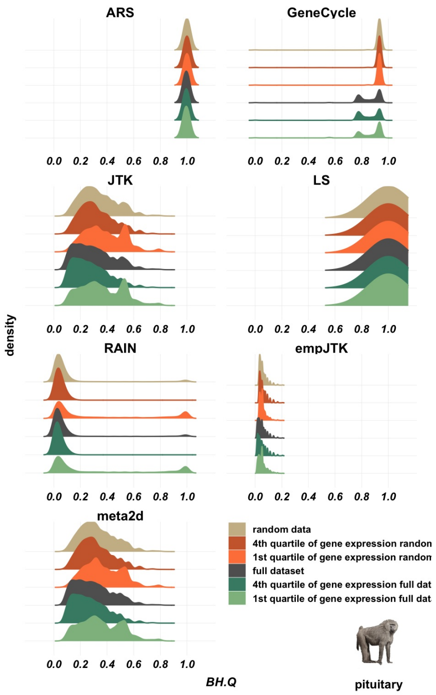


**Fig. S78**

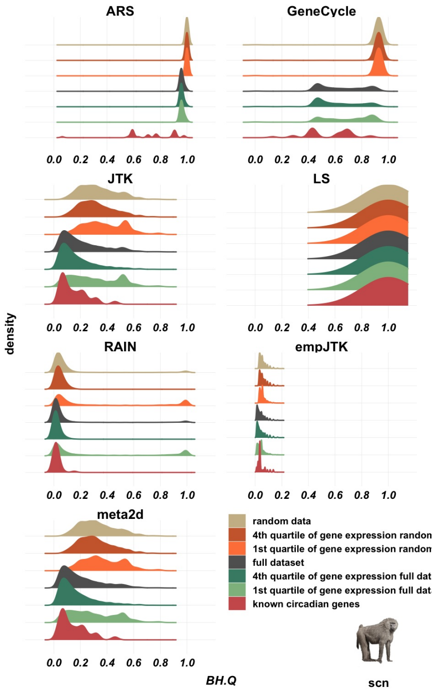


**Fig. S78**

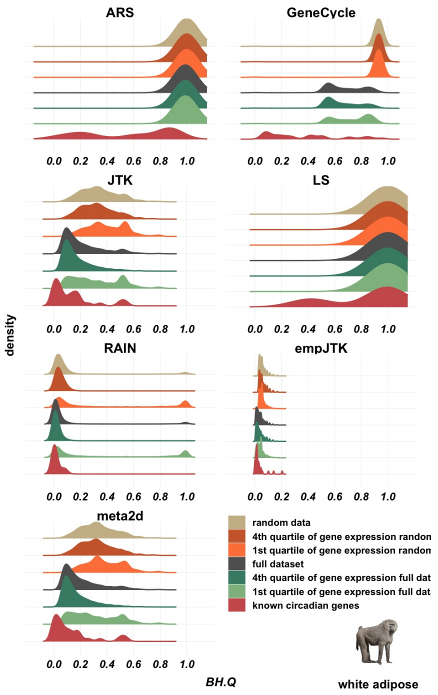


### Fig. S78

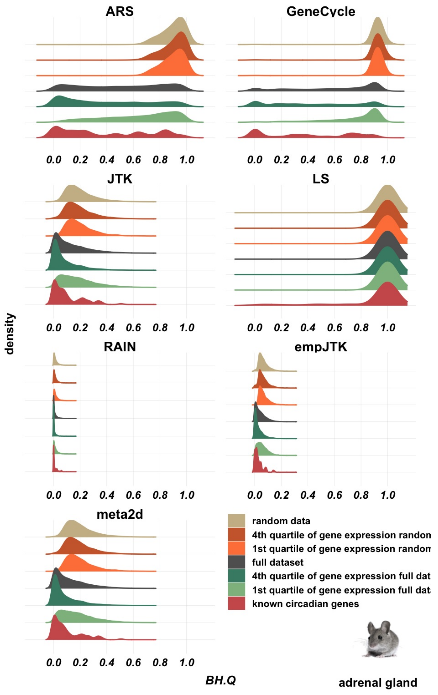


### Fig. S78

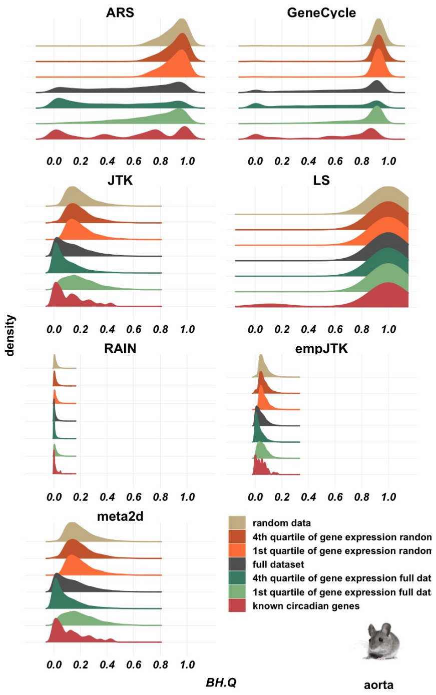


### Fig. S78

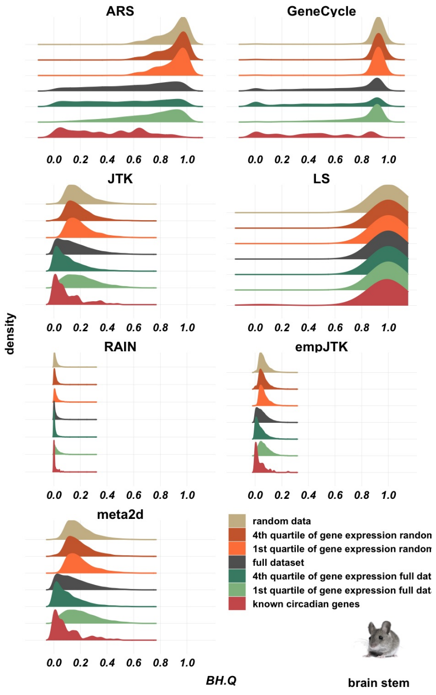


### Fig. S78

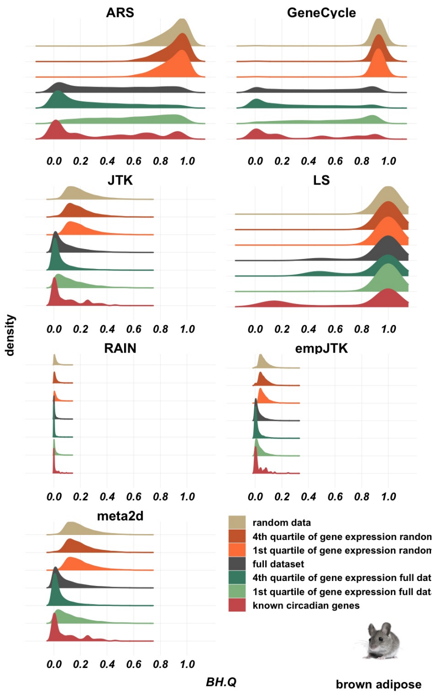


### Fig. S78

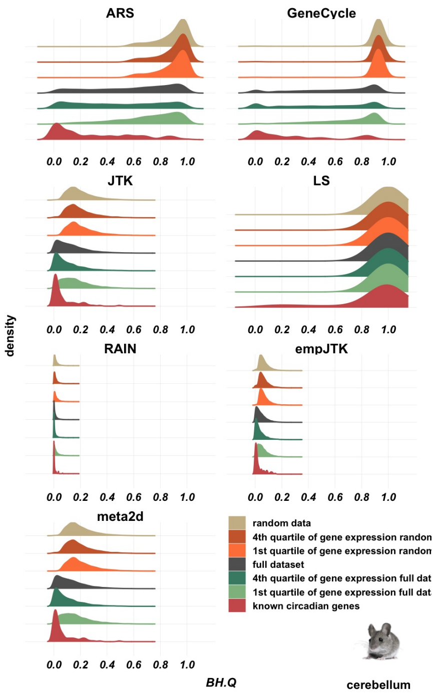


### Fig. S78

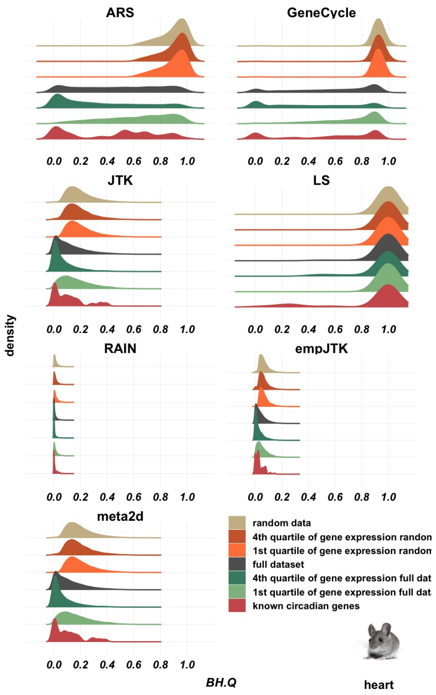


Fig. S78

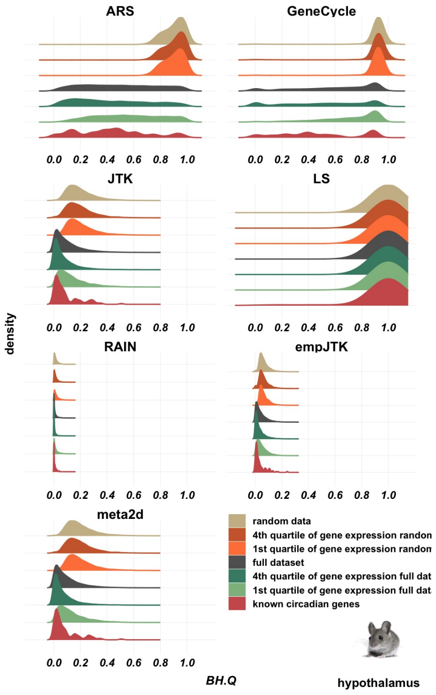


Fig. S78

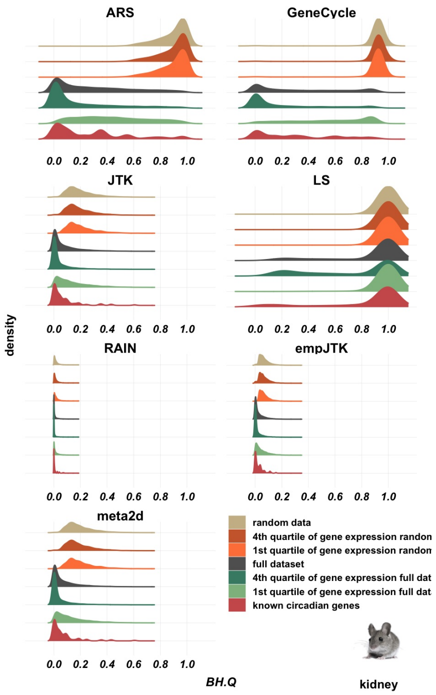


### Fig. S78

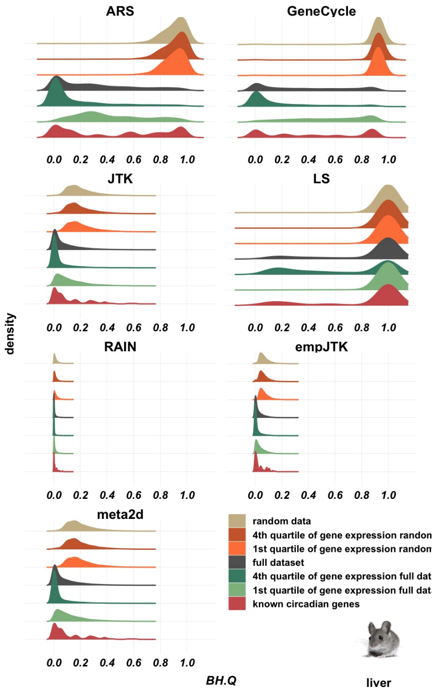


### Fig. S78

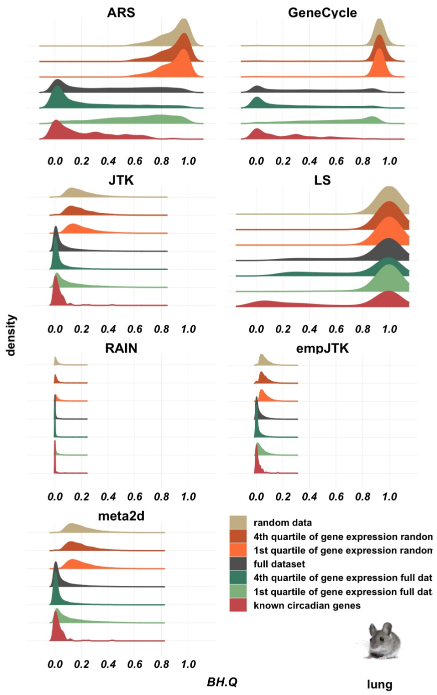


### Fig. S78

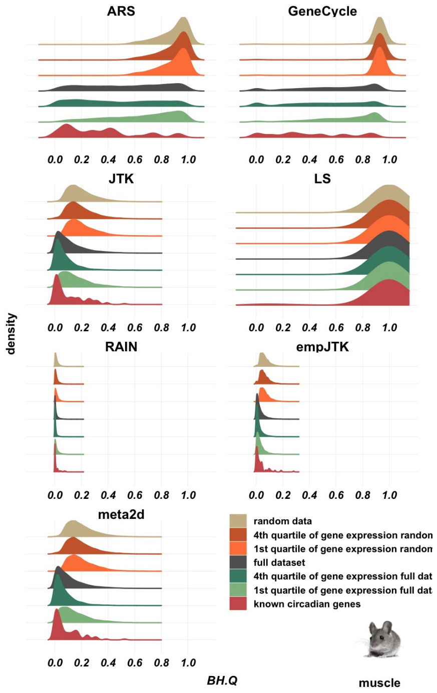


### Fig. S78

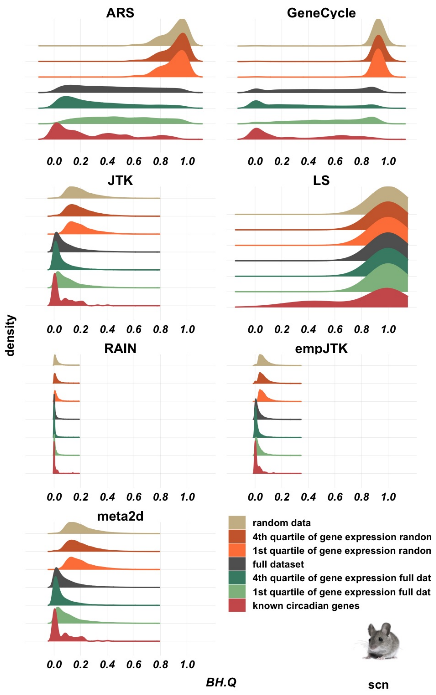


### Fig. S78

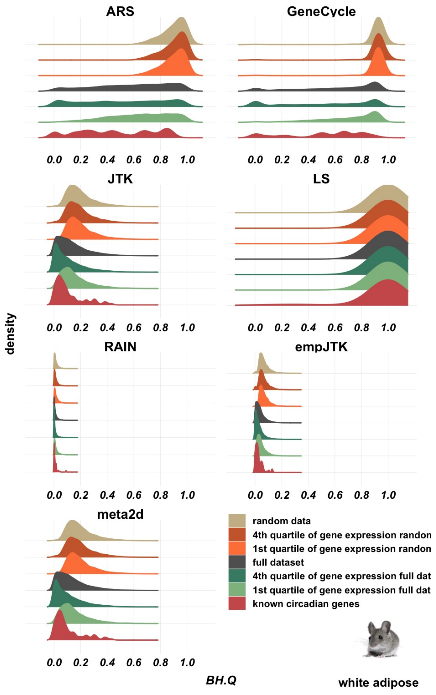


### Fig. S78

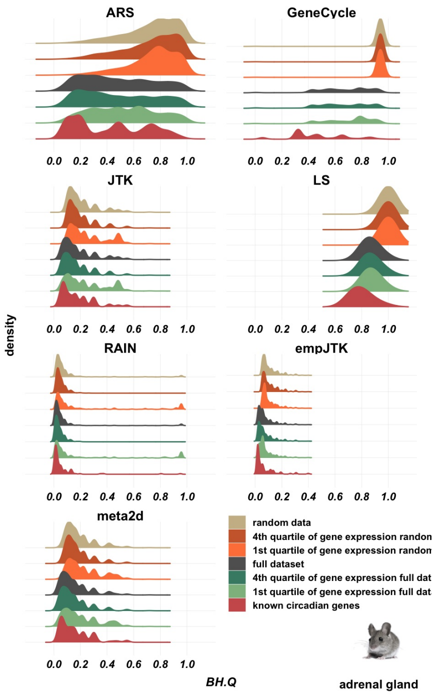


**Fig. S78**

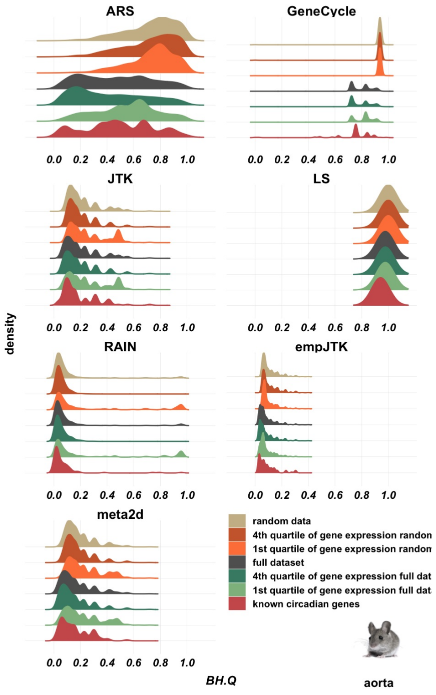


**Fig. S78**

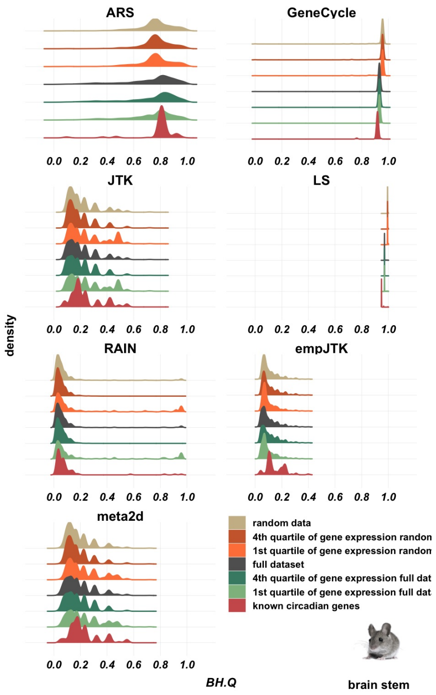


### Fig. S78

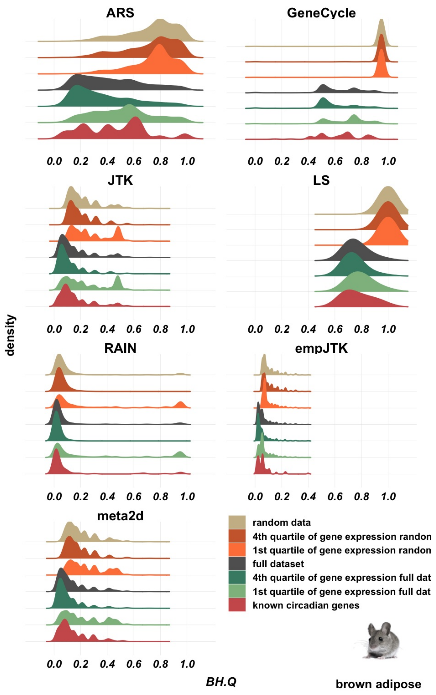


### Fig. S78

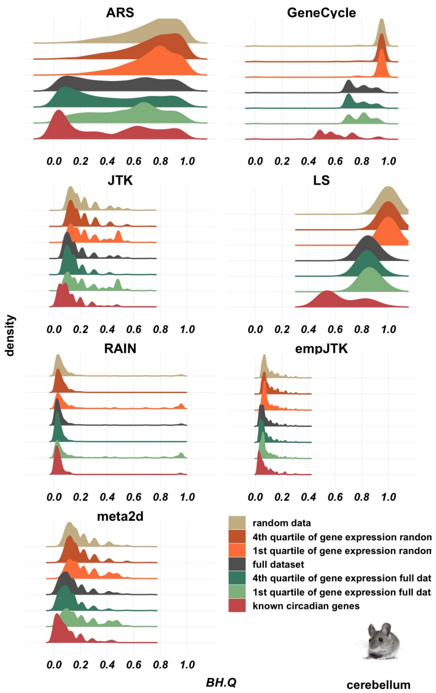


### Fig. S78

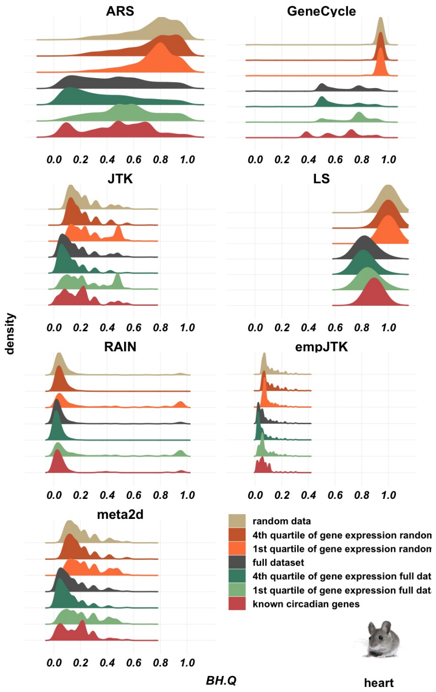


### Fig. S78

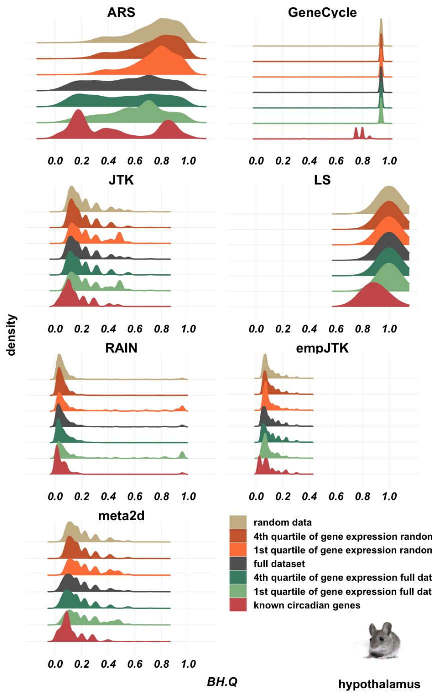


### Fig. S78

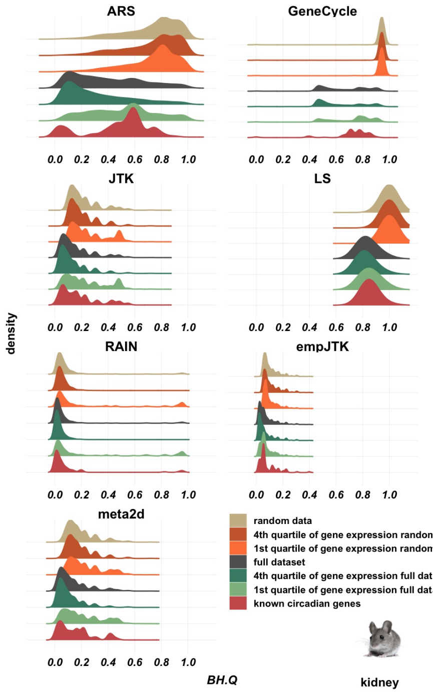


### Fig. S78

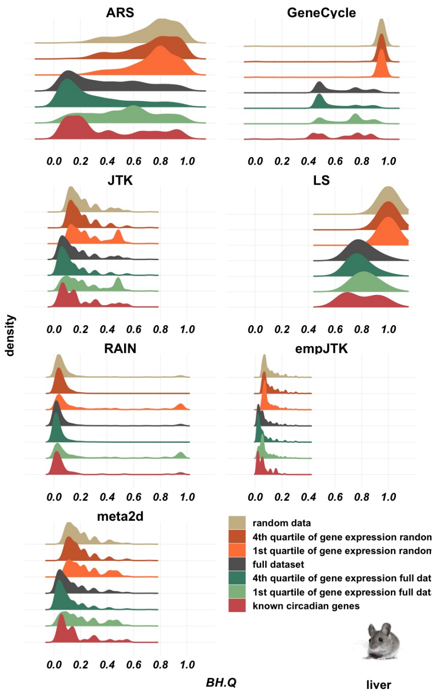


### Fig. S78

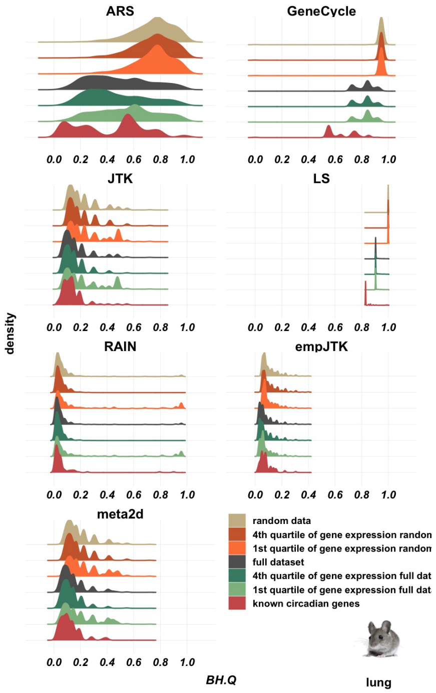


### Fig. S78

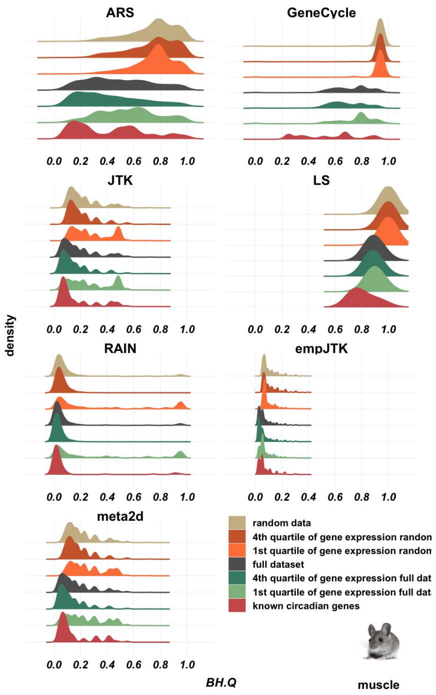


### Fig. S78

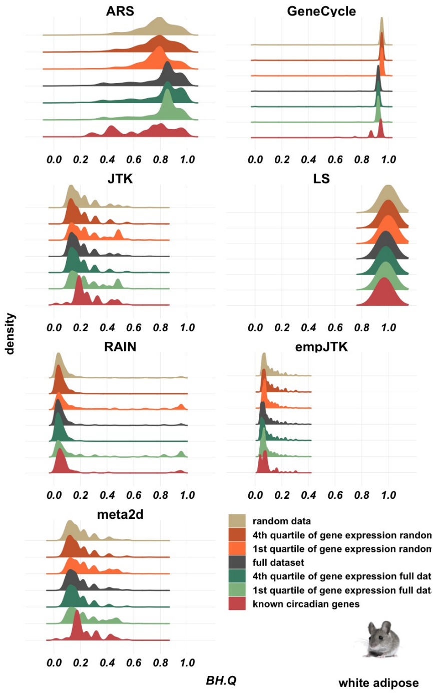


### Fig. S78

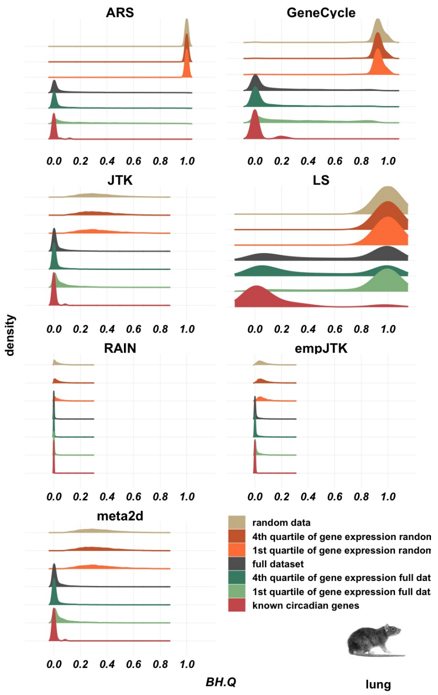


Fig. S78

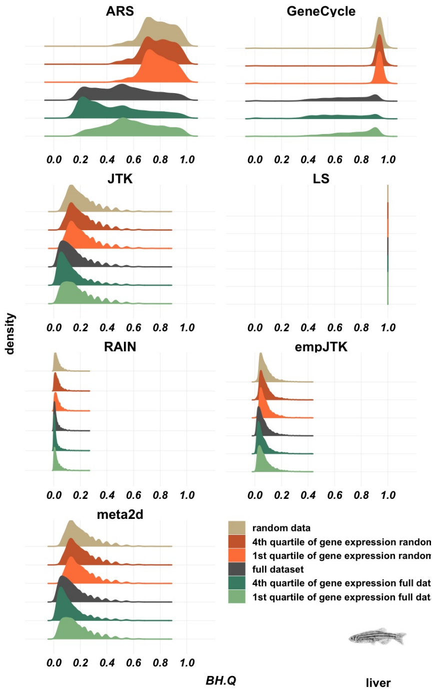


Fig. S78
