## Supplementary figures and images for "Methods detecting rhythmic gene expression are biologically relevant only for strong signal"

### Additional file 3

#### VERTEBRATES

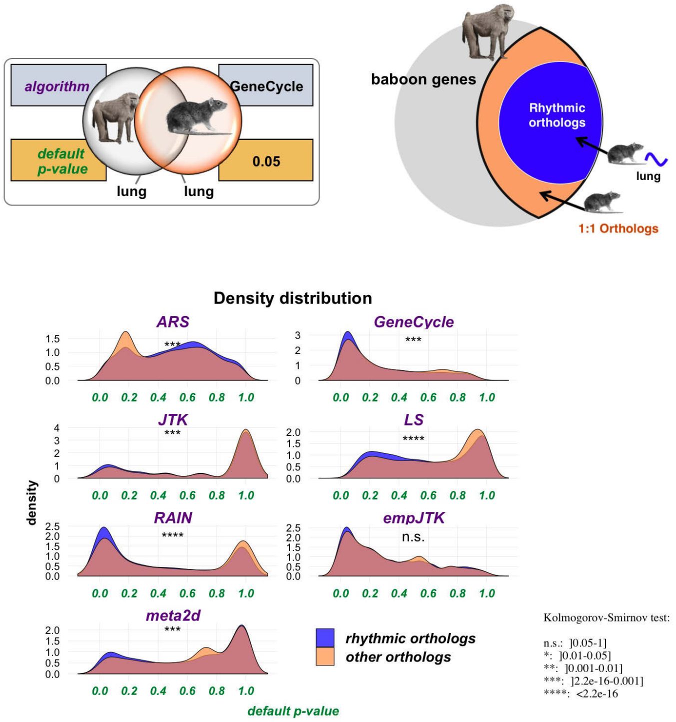


**Fig. S1**

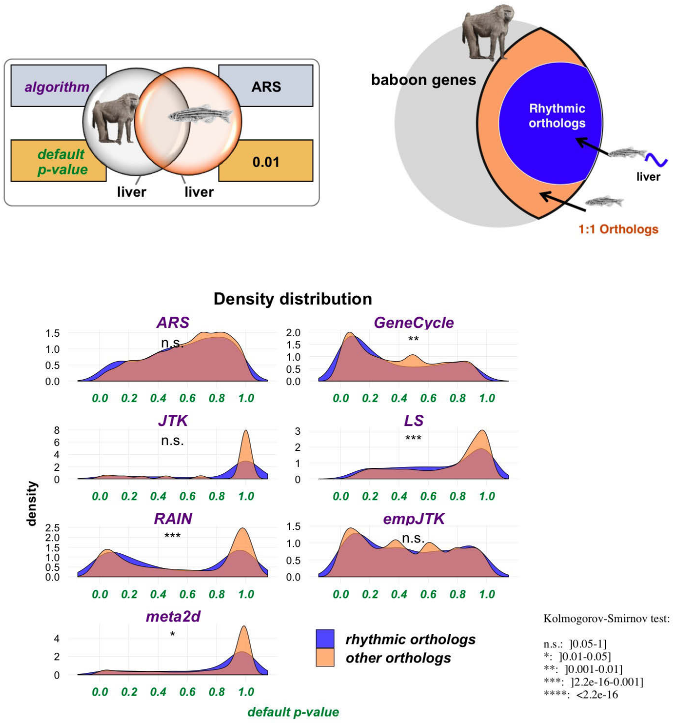

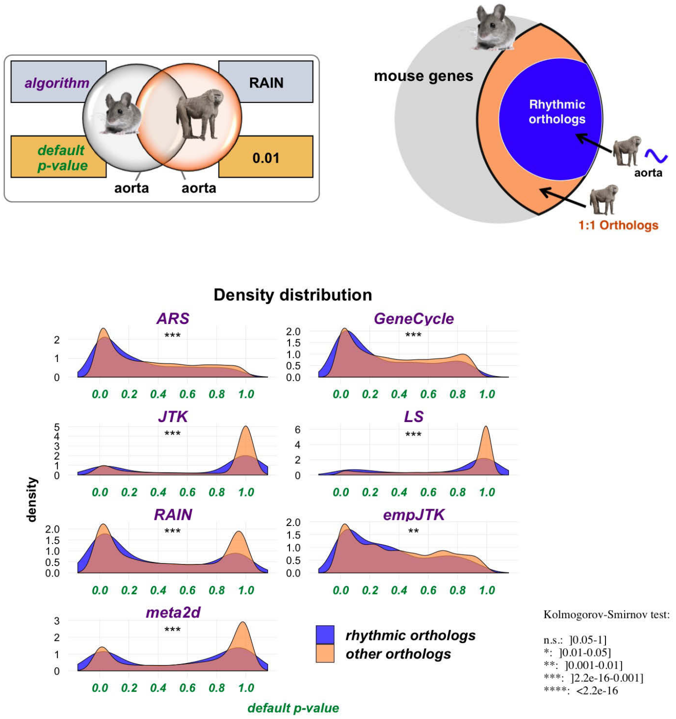

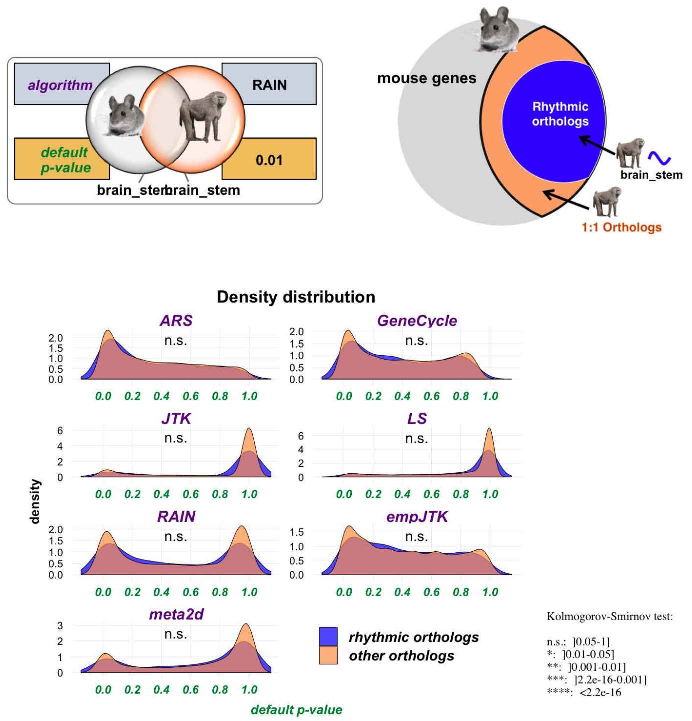

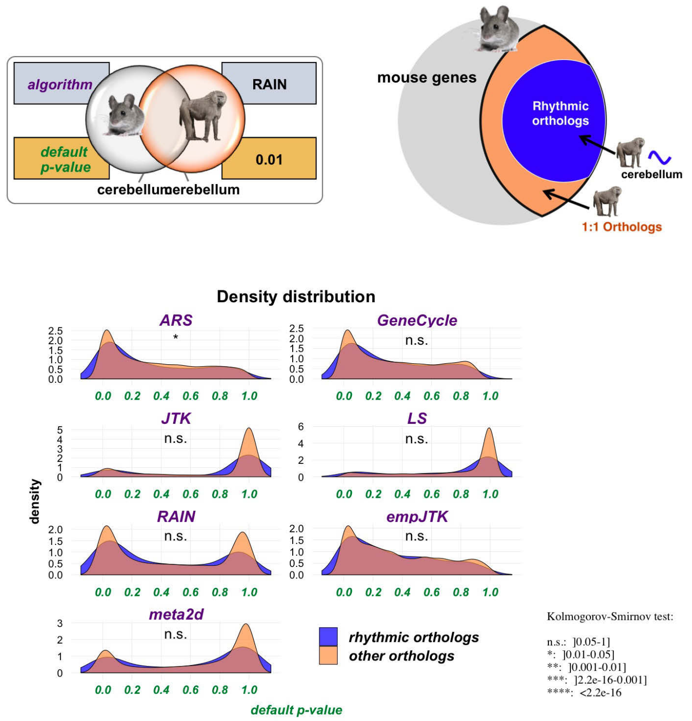

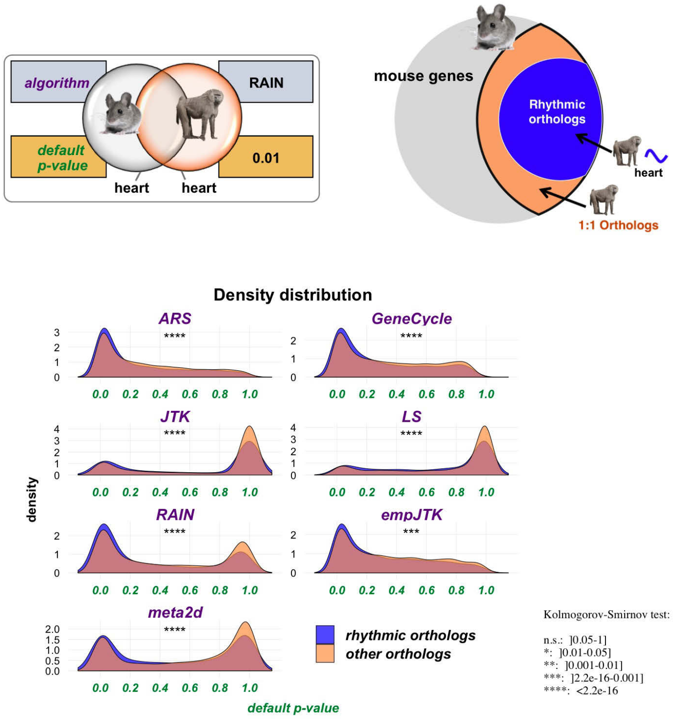

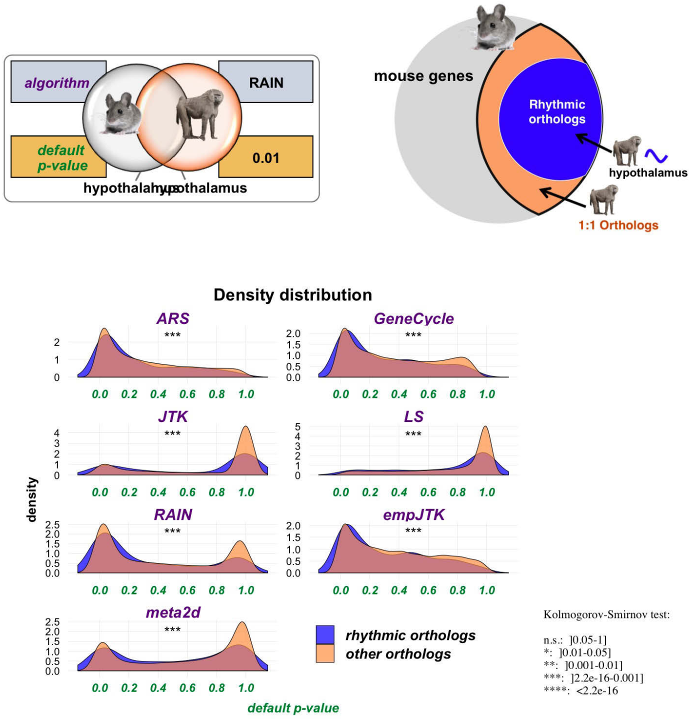

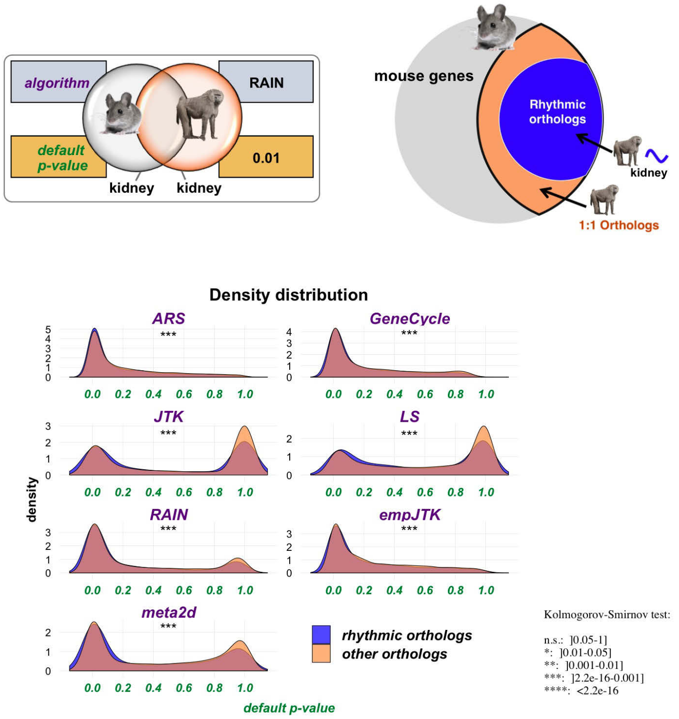

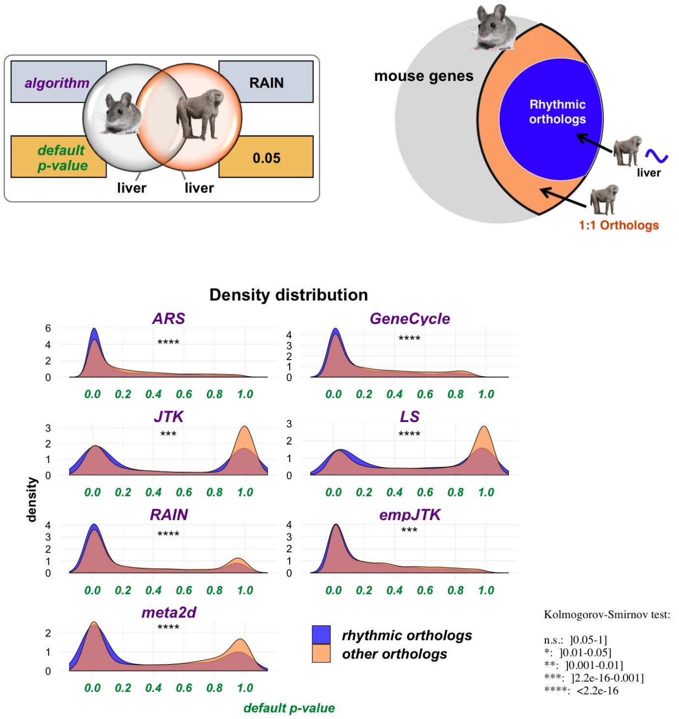

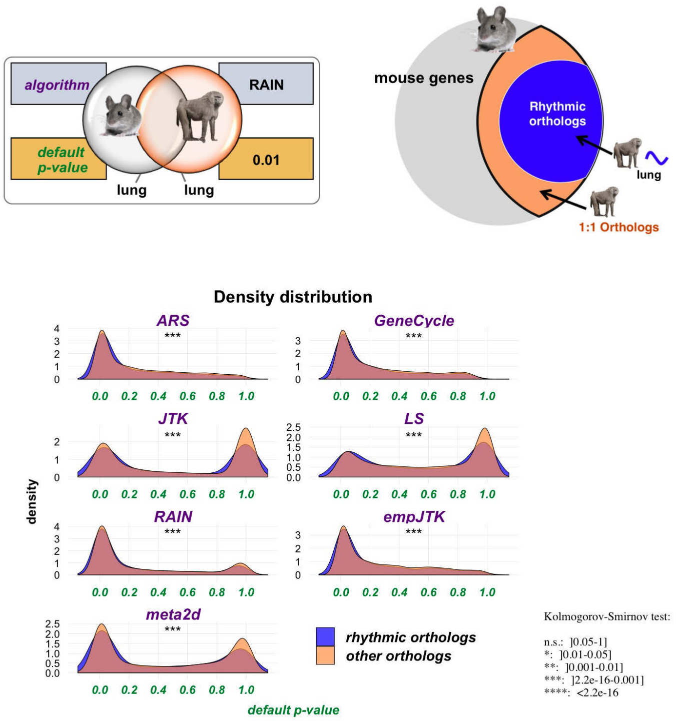


#### Fig. S10

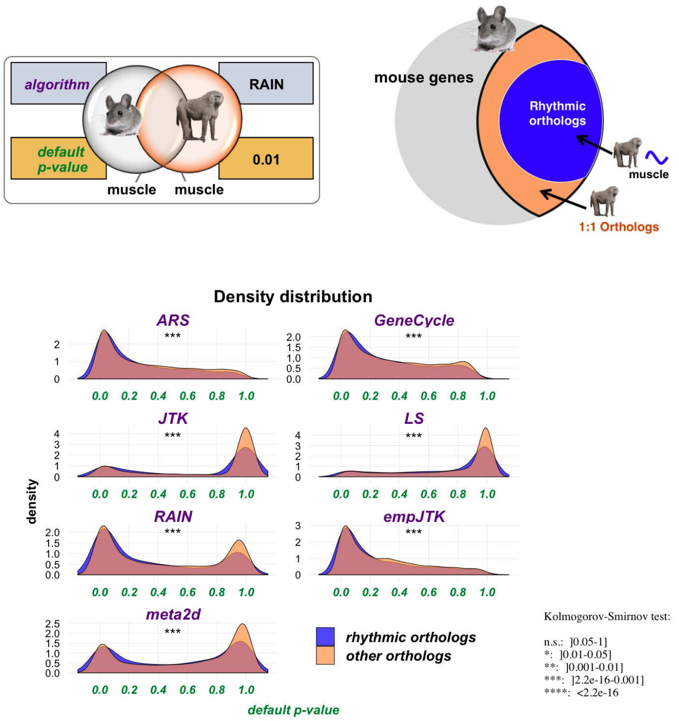


#### Fig. S11

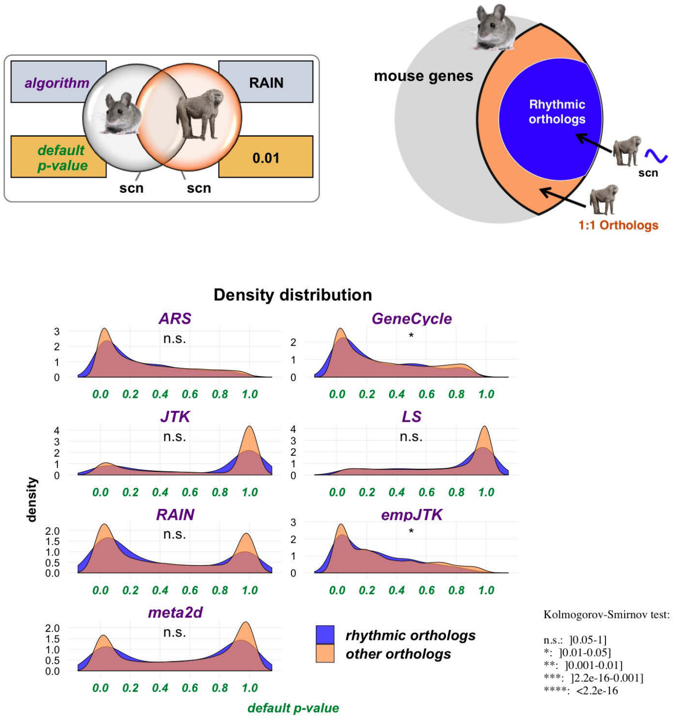


#### Fig. S12

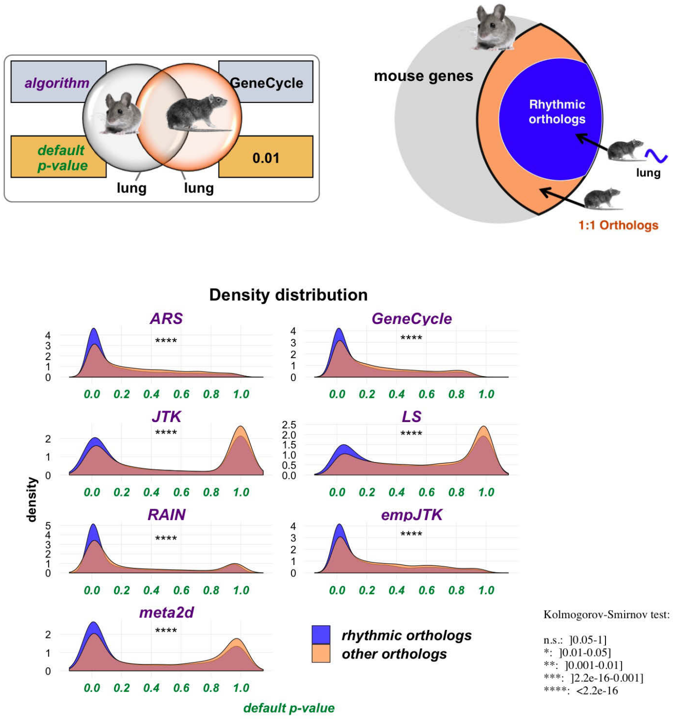


#### Fig. S13

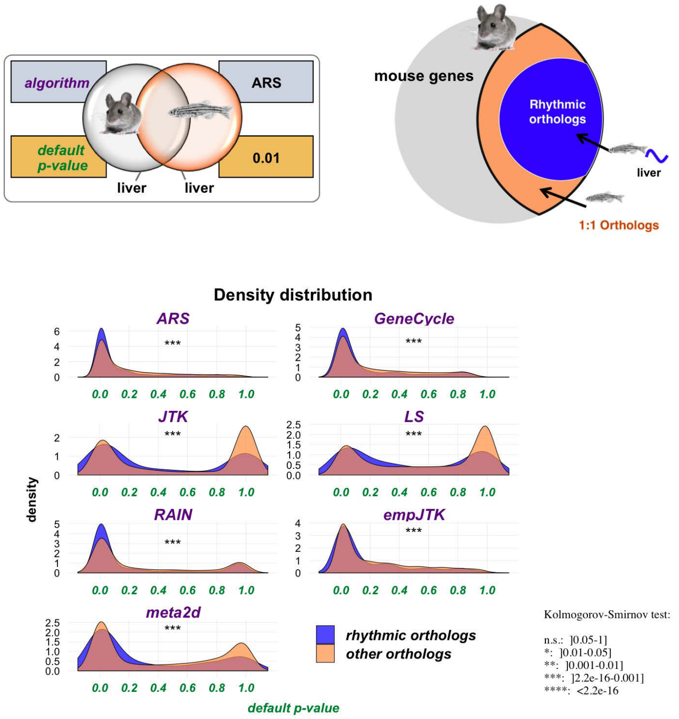
