## Additional file 4 for "Methods detecting rhythmic gene expression are biologically relevant only for strong signal"

### VERTEBRATES

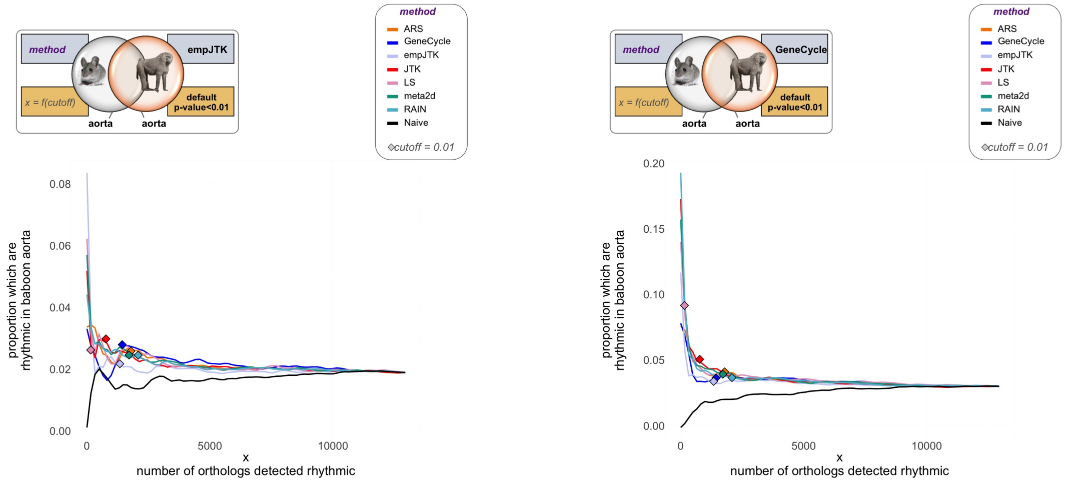


**Fig. S1**

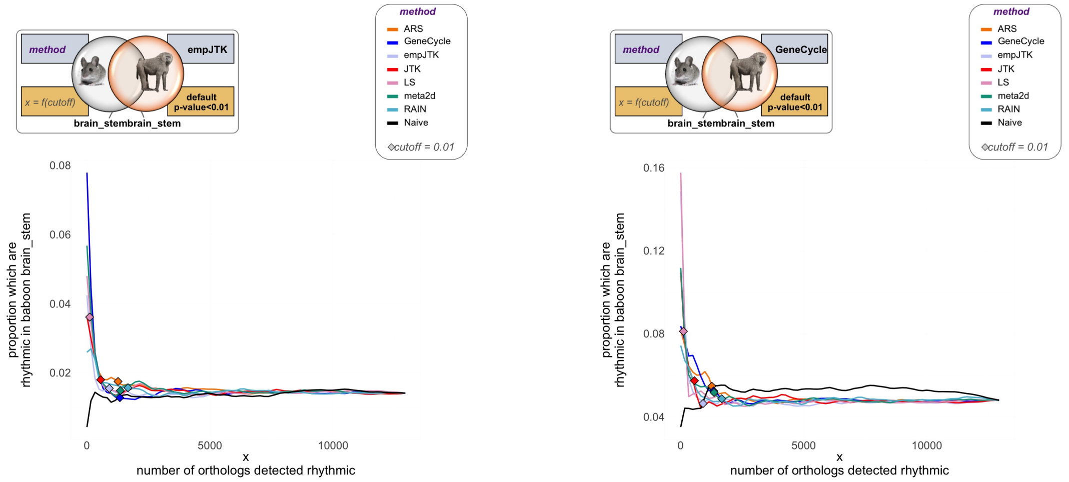


**Fig. S2**

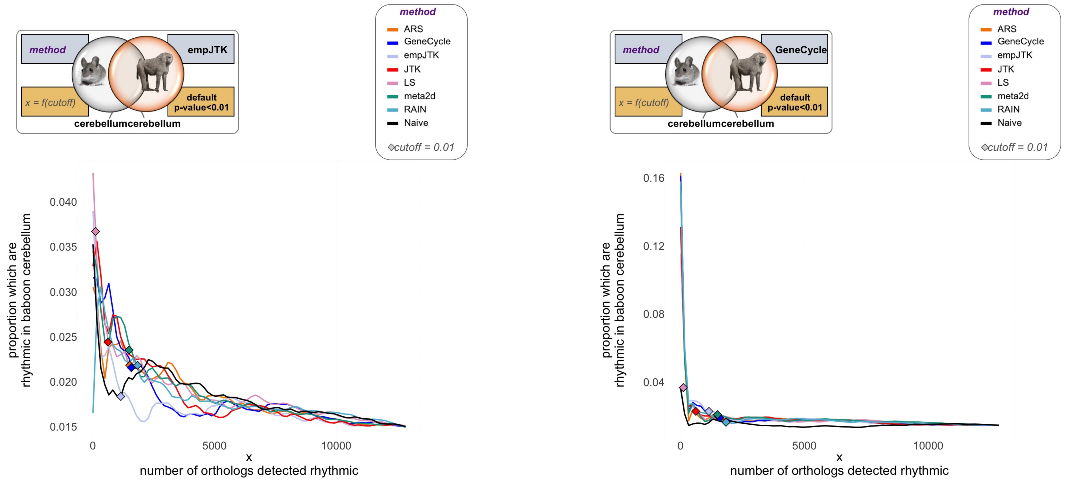


### Fig. S3

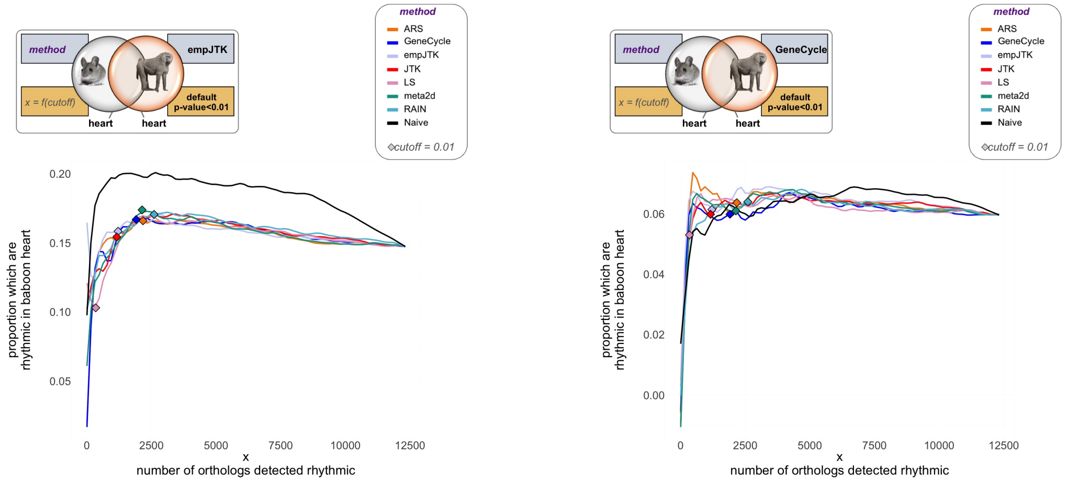


### Fig. S4

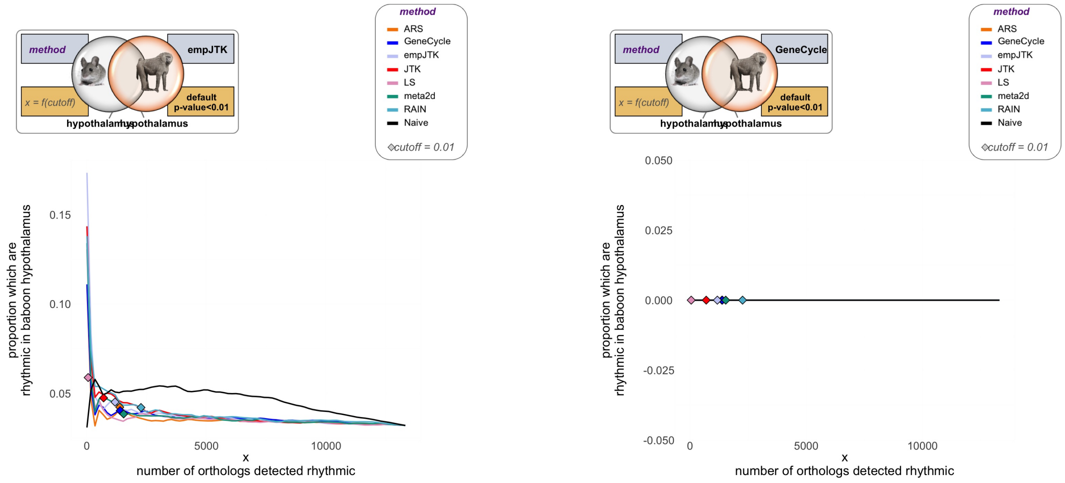


### Fig. S5

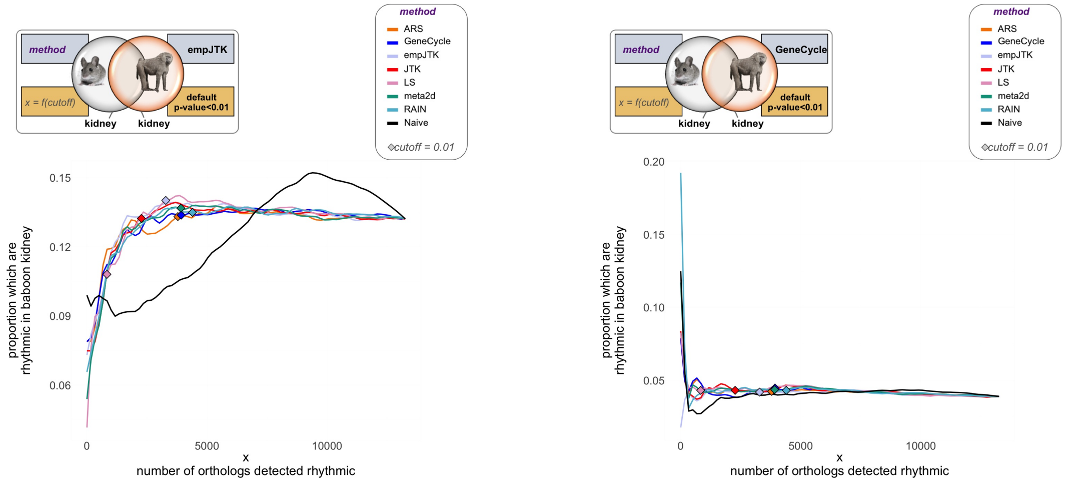


### Fig. S6

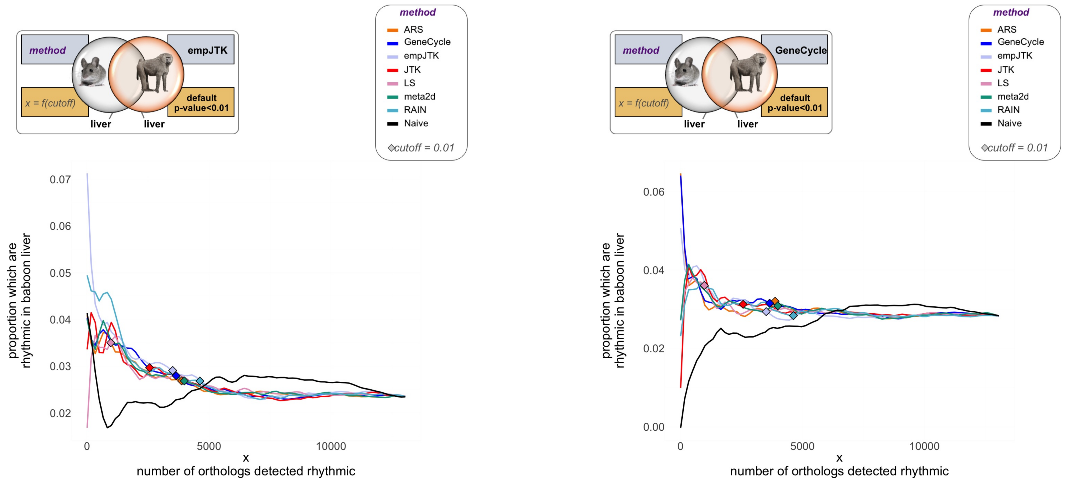


### Fig. S7

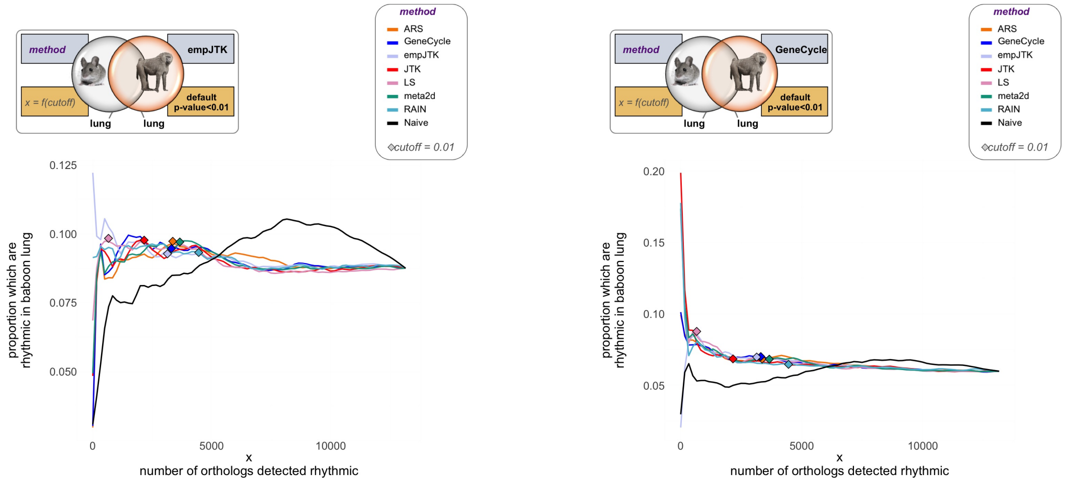


### Fig. S8

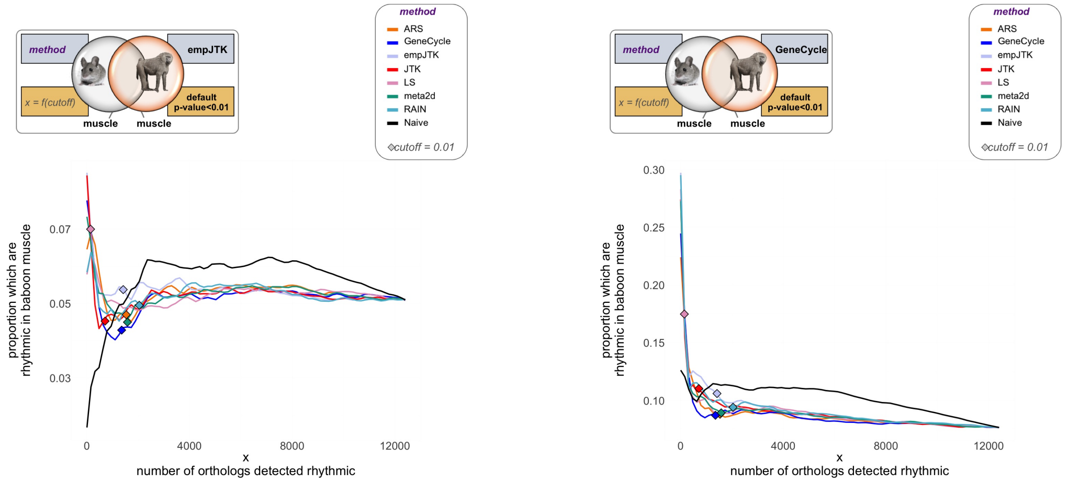


### Fig. S9

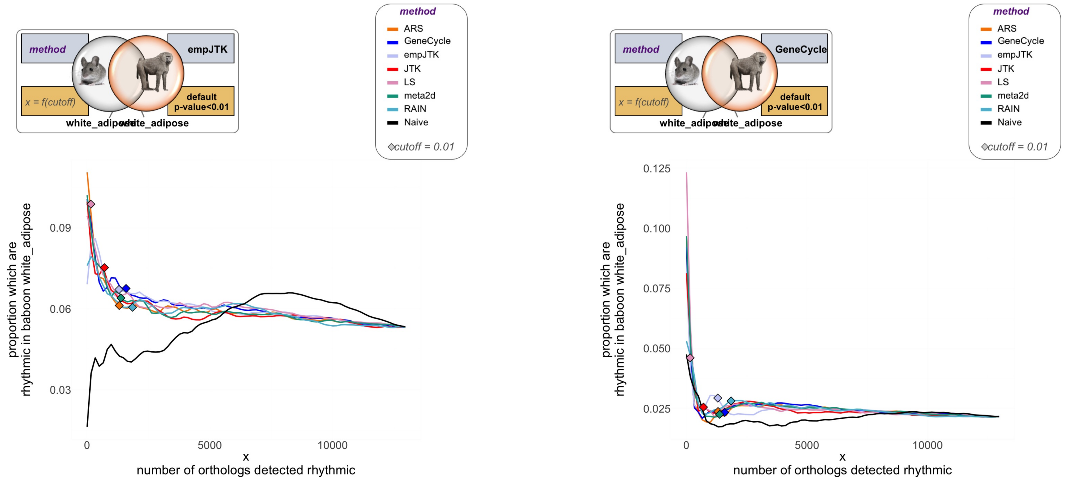


### Fig. S10

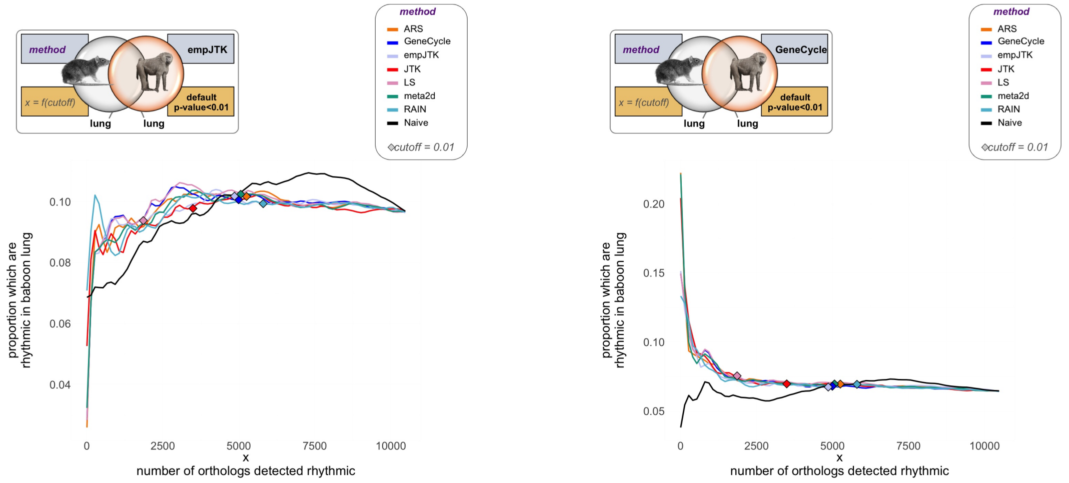


### Fig. S11

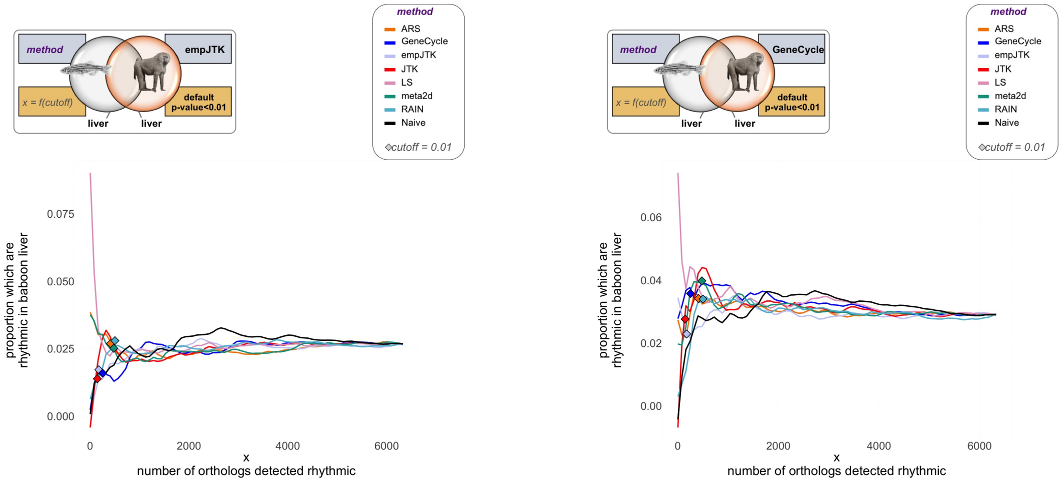


### Fig. S12

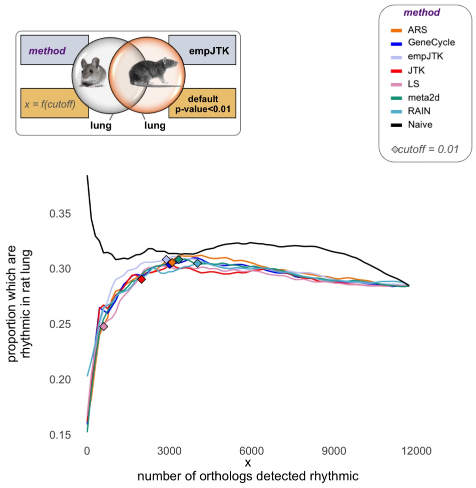


### Fig. S13

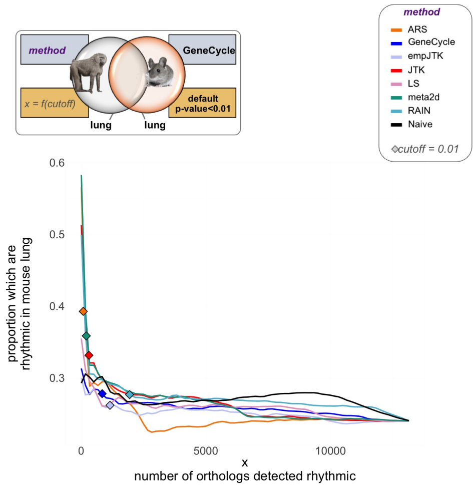

### Fig. S14

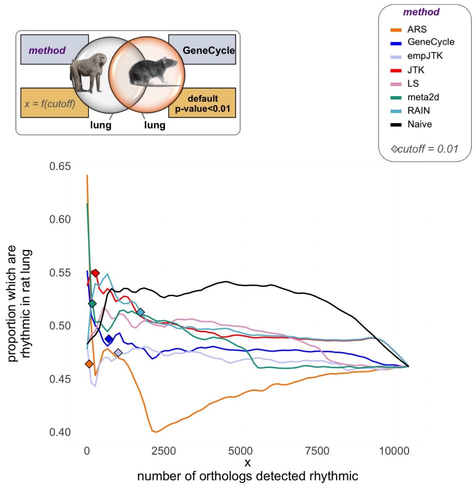


### Fig. S15

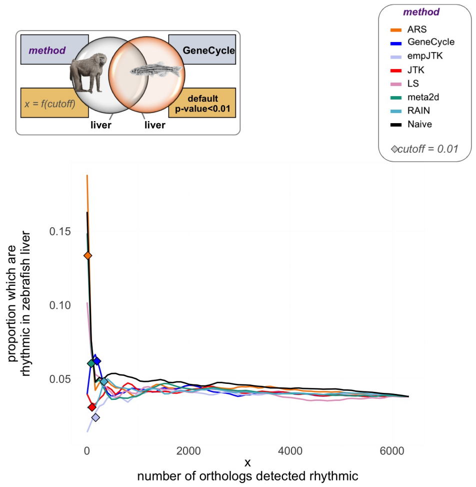


### Fig. S16

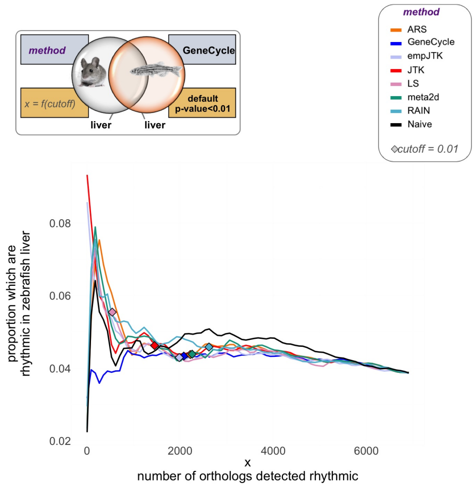

### Fig. S17

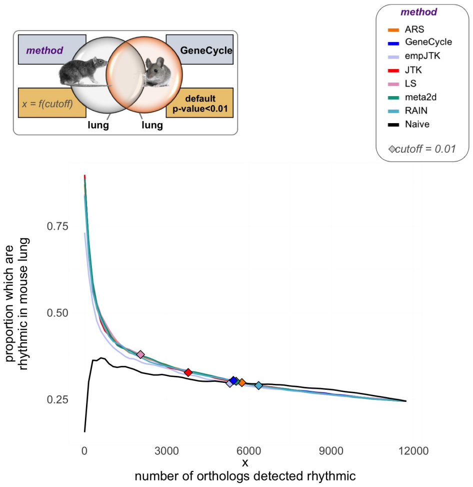


### Fig. S18

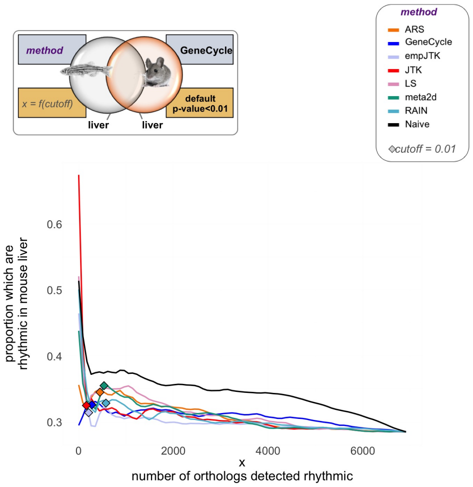


### Fig. S19
