## Additional file 5 for "Methods detecting rhythmic gene expression are biologically relevant only for strong signal"

### INSECTS

### Density distribution of raw and default *p*-values

#### default *p*-values

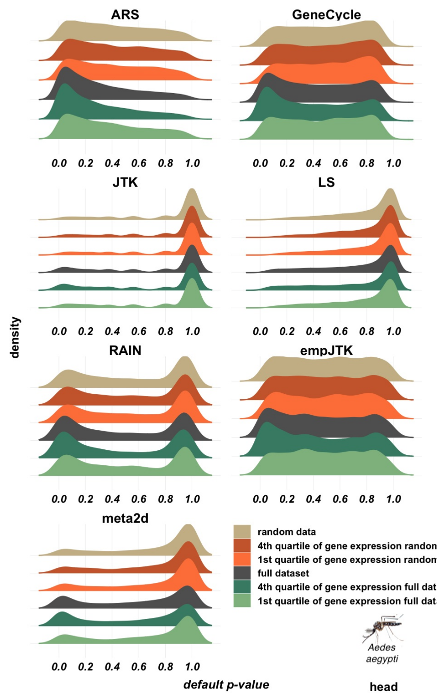


**Fig. S1**

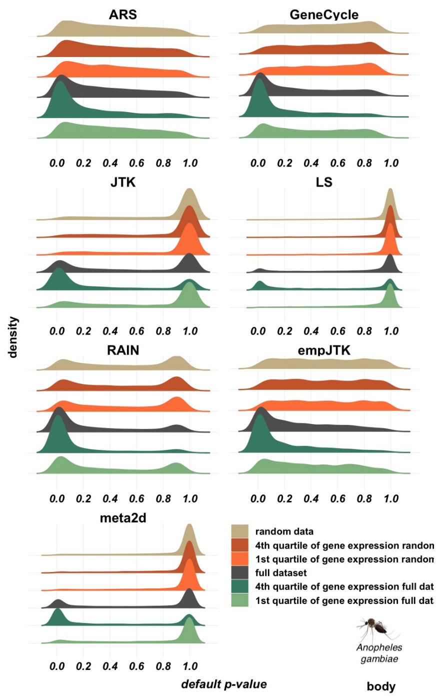
ARS
GeneCycle
JTK
LS
RAIN
empJTK
meta2d
density
0.0 0.2 0.4 0.6 0.8 1.0
default p-value
random data
4th quartile of gene expression randon
1st quartile of gene expression randon
full dataset
4th quartile of gene expression full dat
1st quartile of gene expression full dat
Anopheles gambiae
body

### Fig. S2

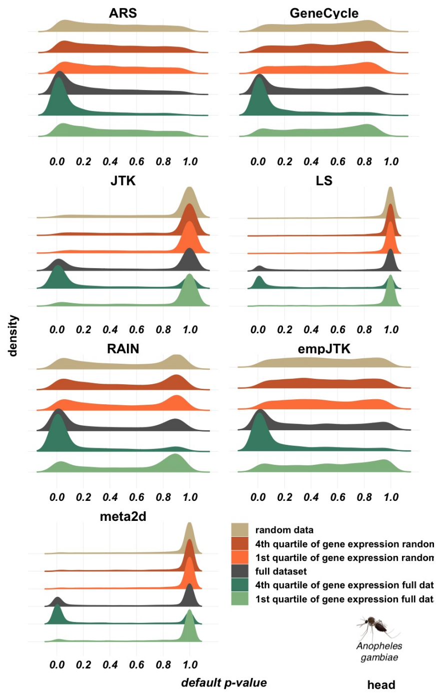
ARS
GeneCycle
JTK
LS
RAIN
empJTK
meta2d
density
0.0 0.2 0.4 0.6 0.8 1.0
default p-value
random data
4th quartile of gene expression randon
1st quartile of gene expression randon
full dataset
4th quartile of gene expression full dat
1st quartile of gene expression full dat
Anopheles gambiae
head

### Fig. S3

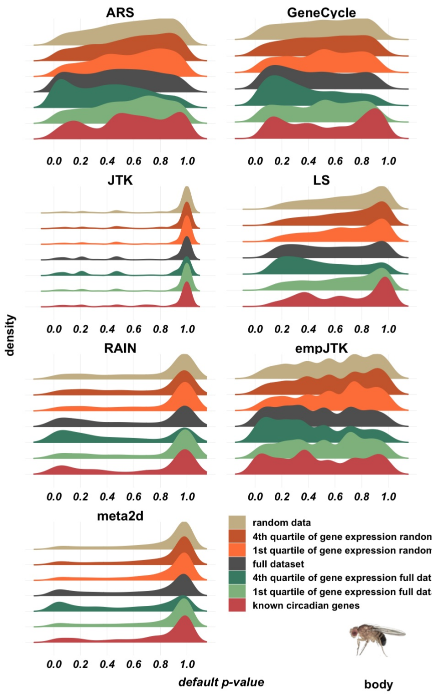

ARS
GeneCycle
JTK
LS
RAIN
empJTK
meta2d
density
0.0 0.2 0.4 0.6 0.8 1.0
default p-value
random data
4th quartile of gene expression randon
1st quartile of gene expression randon
full dataset
4th quartile of gene expression full dat
1st quartile of gene expression full dat
known circadian genes
body


### Fig. S4

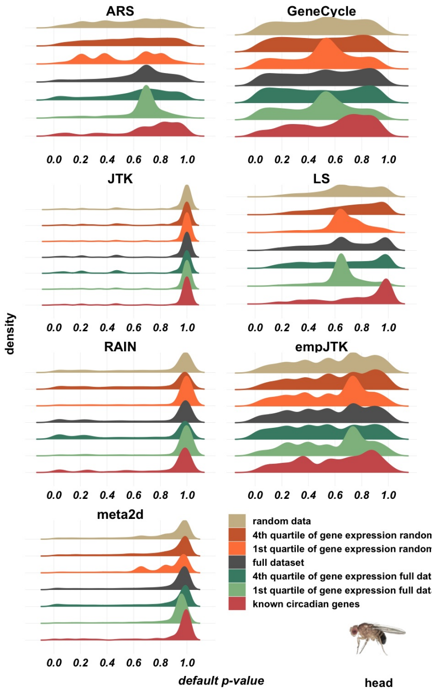

ARS
GeneCycle
JTK
LS
RAIN
empJTK
meta2d
density
0.0 0.2 0.4 0.6 0.8 1.0
default p-value
random data
4th quartile of gene expression randon
1st quartile of gene expression randon
full dataset
4th quartile of gene expression full dat
1st quartile of gene expression full dat
known circadian genes
head


### Fig. S5

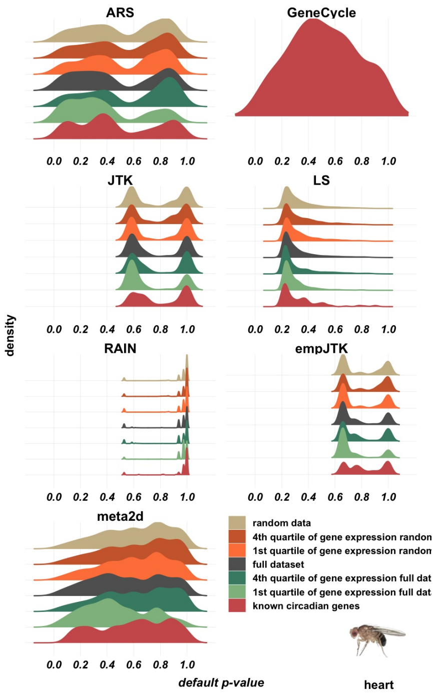


### Fig. S6

### raw *p*-values

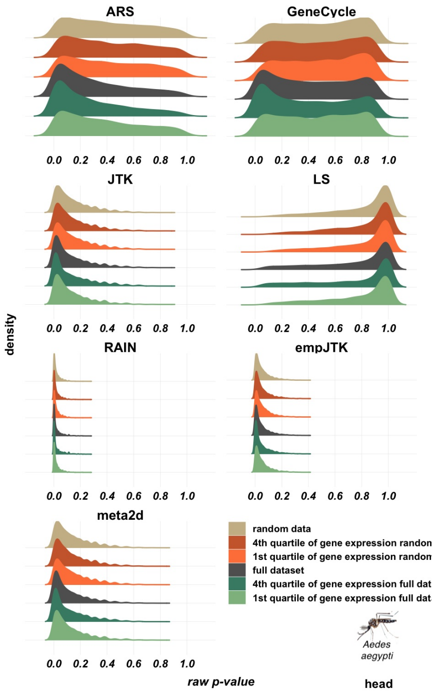


### Fig. S7

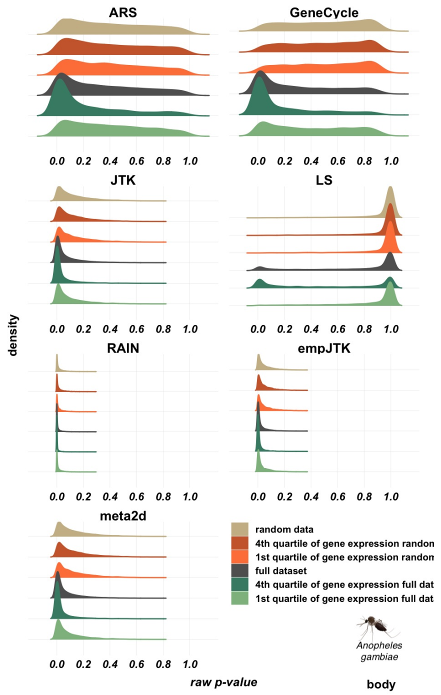
ARS
GeneCycle
JTK
LS
RAIN
empJTK
meta2d
0.0 0.2 0.4 0.6 0.8 1.0
density
raw p-value
random data
4th quartile of gene expression randon
1st quartile of gene expression randon
full dataset
4th quartile of gene expression full dat
1st quartile of gene expression full dat
Anopheles gambiae
body

### Fig. S8

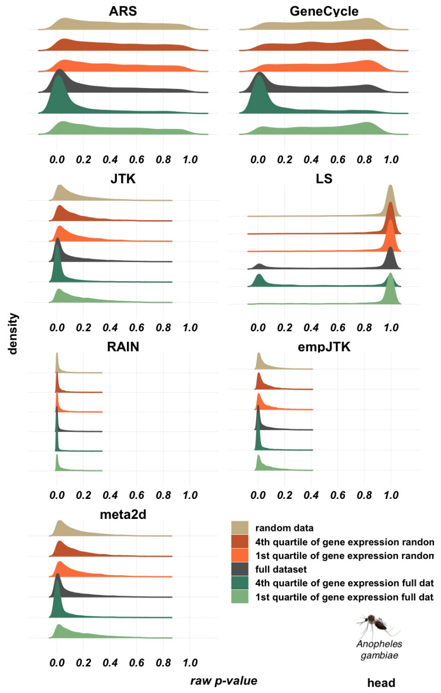
ARS
GeneCycle
JTK
LS
RAIN
empJTK
meta2d
0.0 0.2 0.4 0.6 0.8 1.0
density
raw p-value
random data
4th quartile of gene expression randon
1st quartile of gene expression randon
full dataset
4th quartile of gene expression full dat
1st quartile of gene expression full dat
Anopheles gambiae
head

### Fig. S9

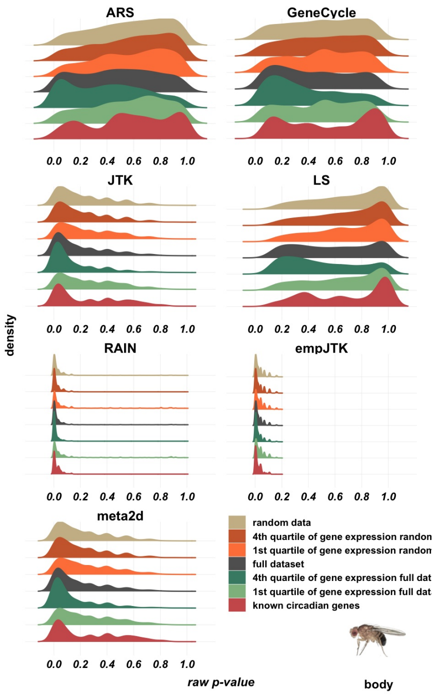


### Fig. S10

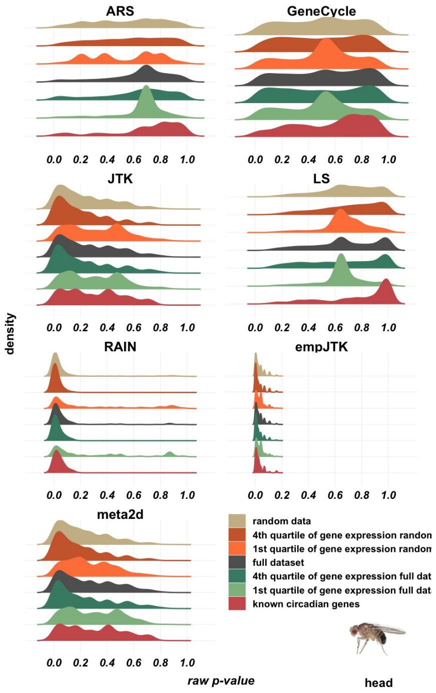


### Fig. S11

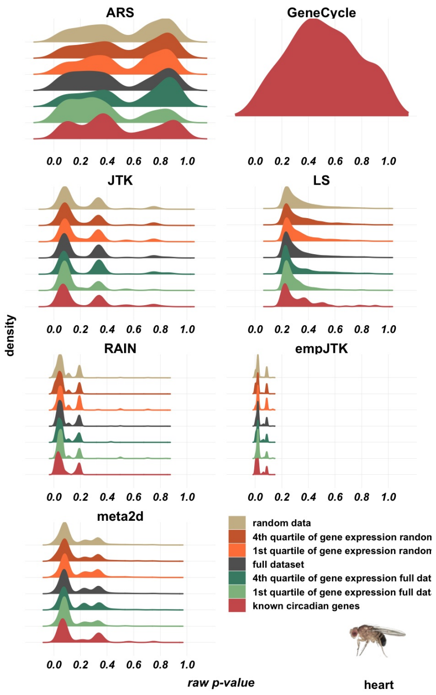
ARS
GeneCycle
JTK
LS
RAIN
empJTK
meta2d
0.0 0.2 0.4 0.6 0.8 1.0
density
raw p-value
random data
4th quartile of gene expression randon
1st quartile of gene expression randon
full dataset
4th quartile of gene expression full dat
1st quartile of gene expression full dat
known circadian genes
heart

### Fig. S12

### Density distribution of *p*-values: rhythmic vs non-rhythmic orthologs

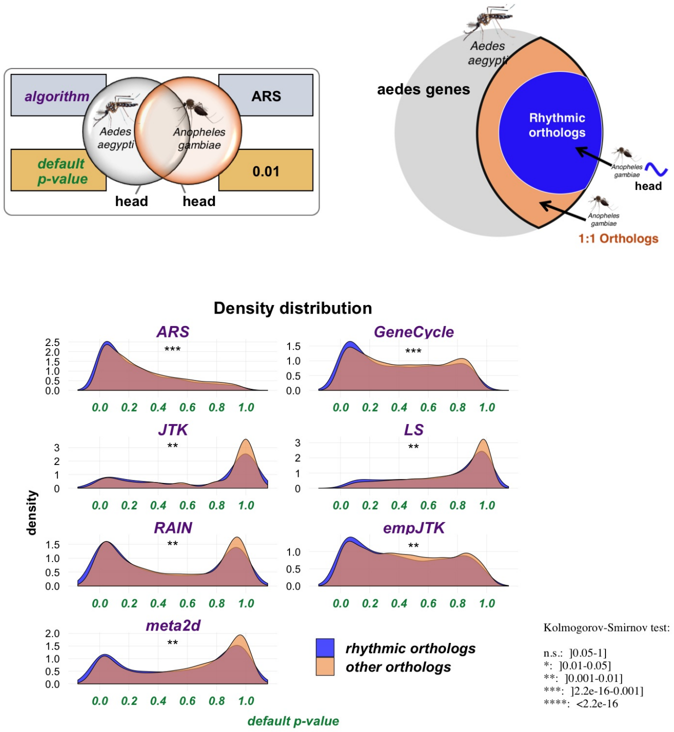


**Fig. S13**

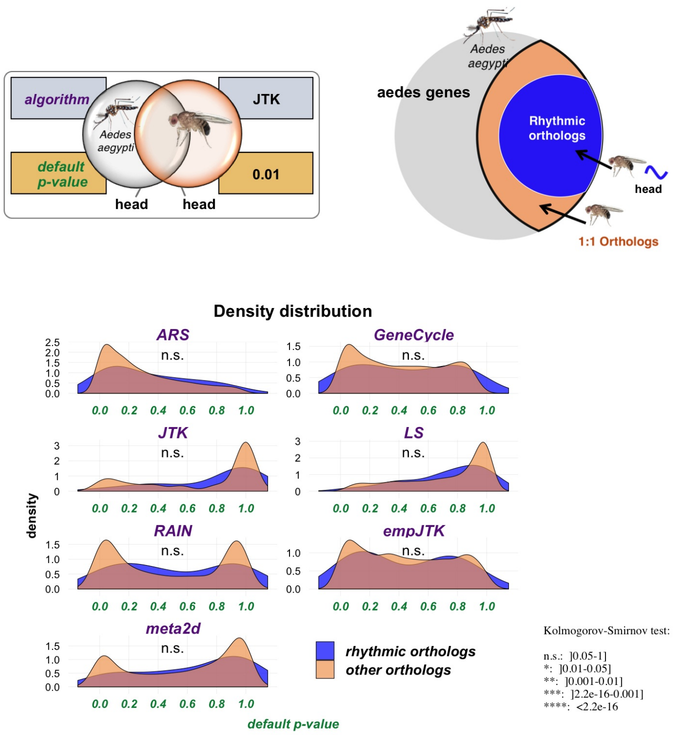


**Fig. S14**

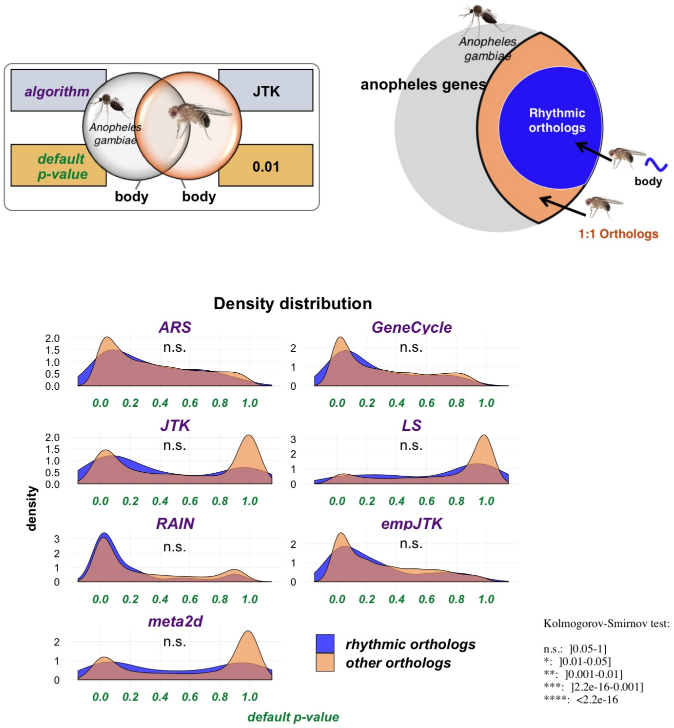


**Fig. S15**

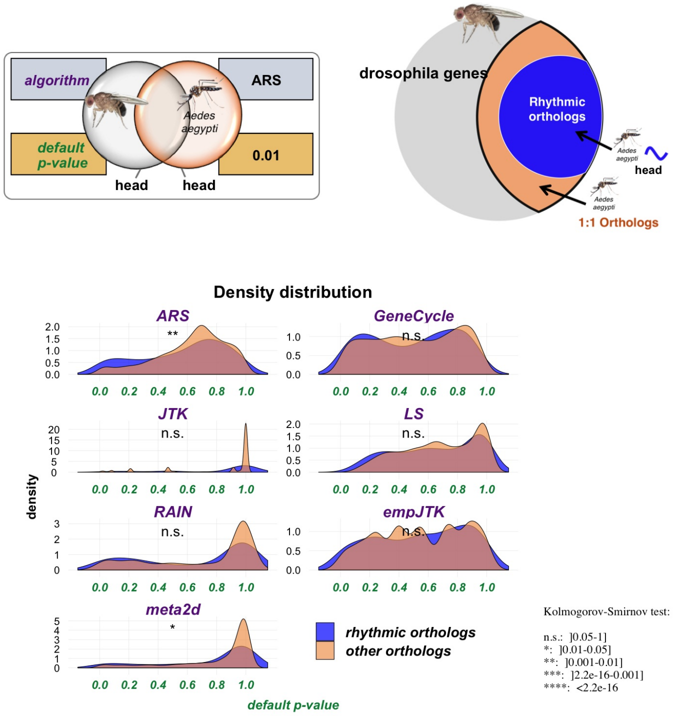


### Fig. S16

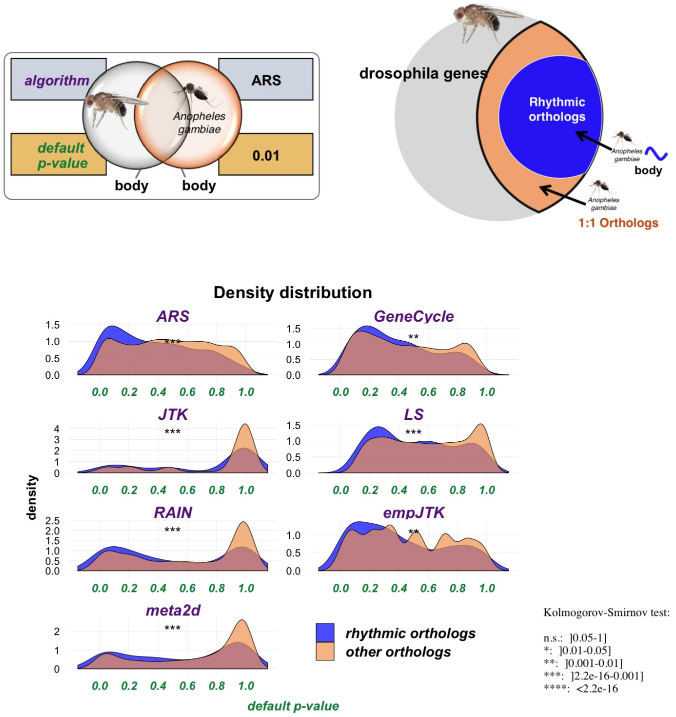


### Fig. S17

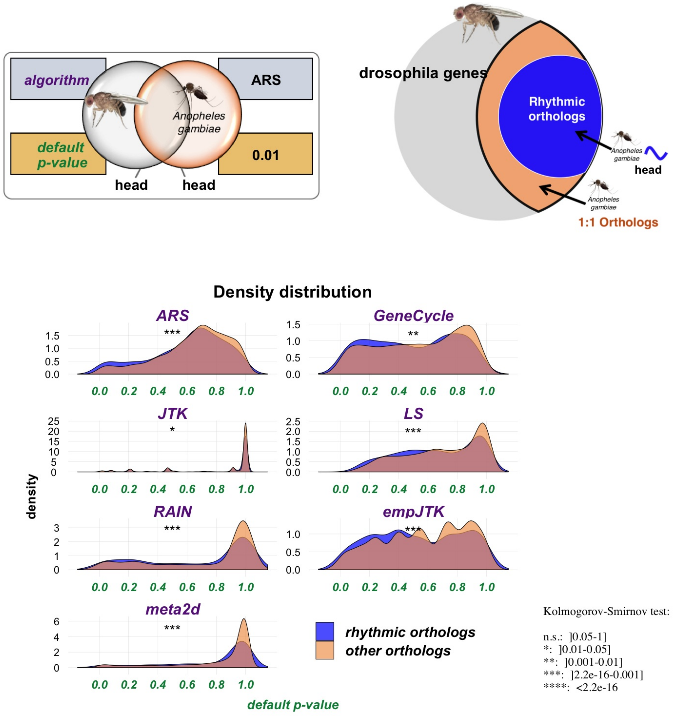

algorithm
ARS
Anopheles gambiae
default p-value
0.01
head
head
drosophila genes
Rhythmic orthologs
Anopheles gambiae
head
1:1 Orthologs
Density distribution
ARS
GeneCycle
JTK
LS
RAIN
empJTK
meta2d
density
default p-value
rhythmic orthologs
other orthologs
Kolmogorov-Smirnov test:
n.s.: ]0.05-1]
*: ]0.01-0.05]
**: ]0.001-0.01]
***: ]2.2e-16-0.001]
****: <2.2e-16


### Fig. S18

### Variation of the proportion A/B as a function of the number of orthologs detected rhythmic

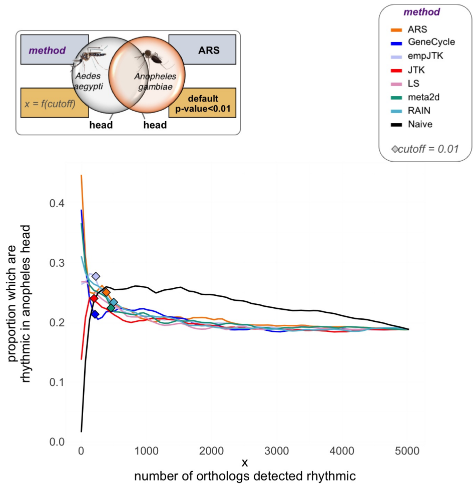


**Fig. S19**

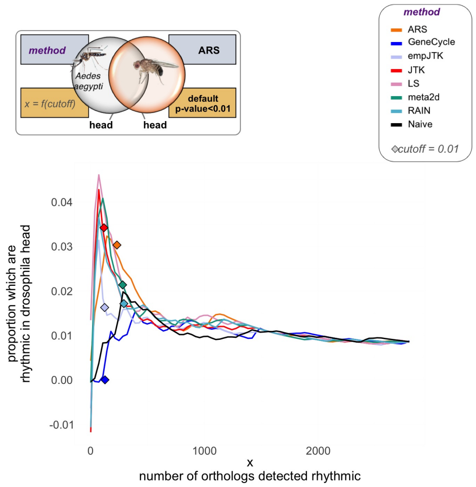


**Fig. S20**

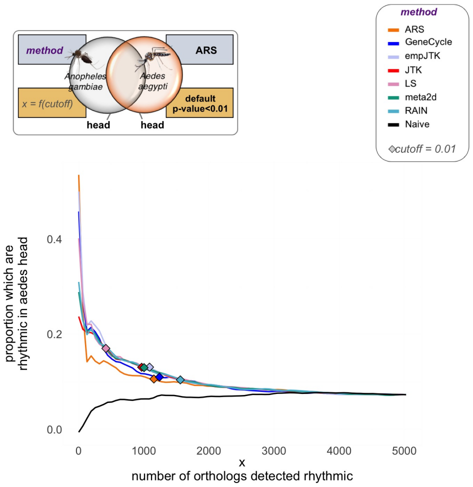


### Fig. S21

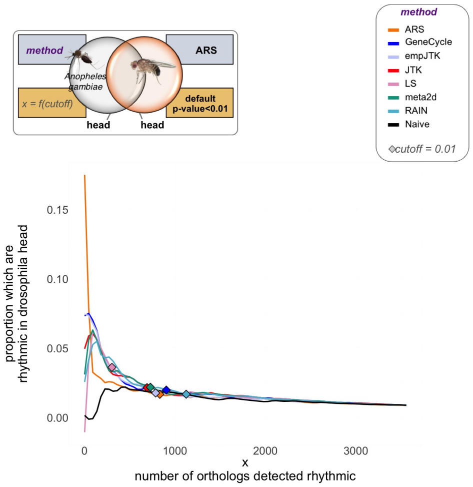


### Fig. S22

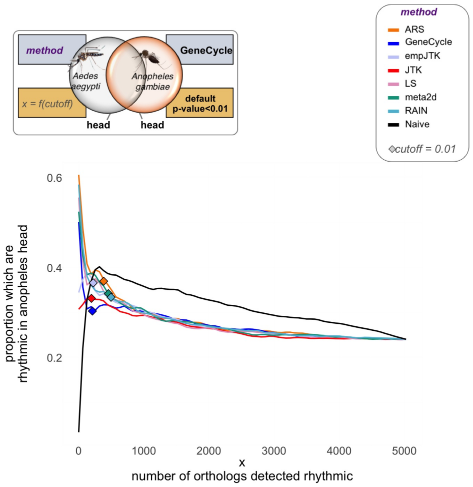


### Fig. S23

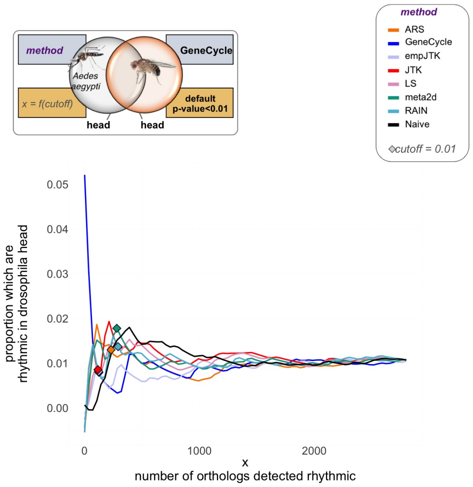


### Fig. S24

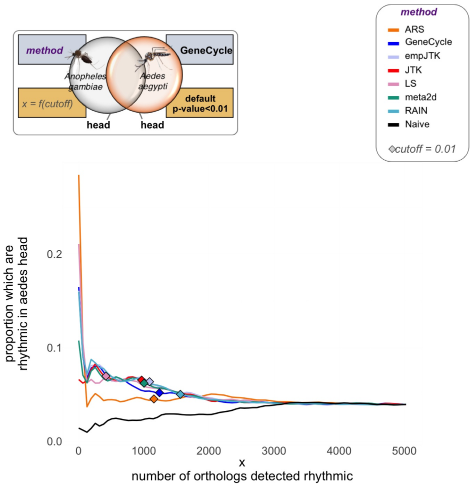


### Fig. S25

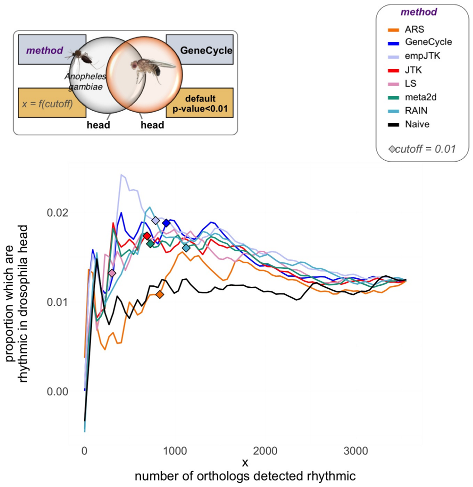


### Fig. S26

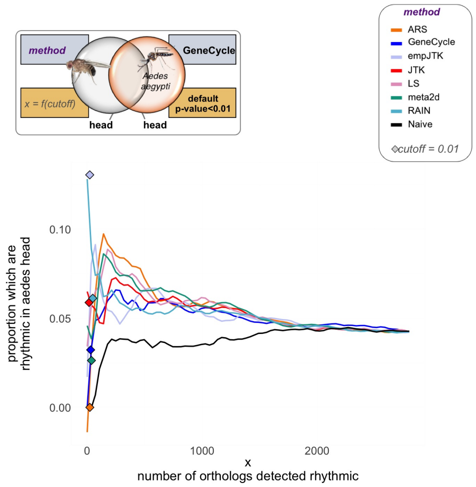


### Fig. S27

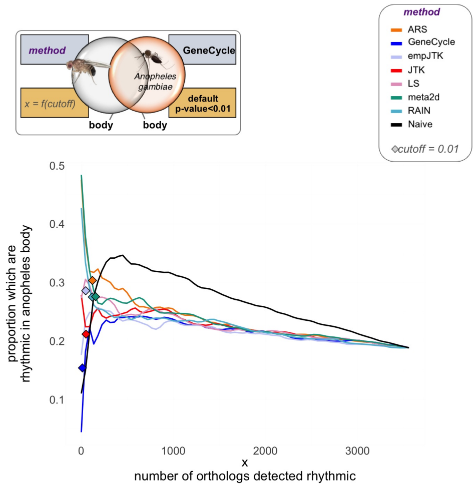


### Fig. S28

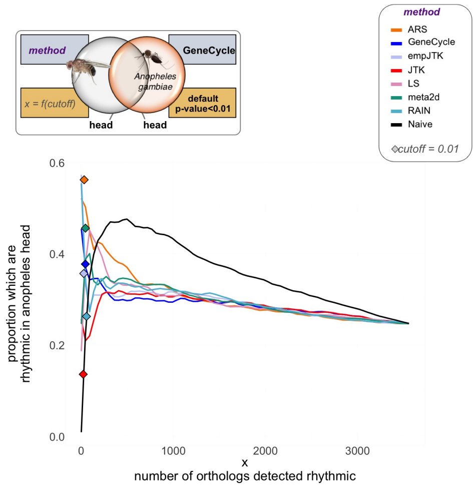


### Fig. S29
